# The SjD Map: An interactive pathway tour into Sjögren’s disease signalling mechanisms

**DOI:** 10.1101/2025.09.08.674876

**Authors:** Sacha E Silva-Saffar, Xavier Mariette, Jacques-Eric Gottenberg, Michele Bombardieri, Divi Cornec, Marta E. Alarcon-Riquelme, PRECISESADS Clinical Consortium, Michael Barnes, Sandra Ng, Wan-Fai Ng, Gaetane Nocturne, Anna Niarakis

**Author notes:** A list of consortia investigators and their affiliations appear in Supplementary Data S1 (PRECISESADS Clinical Consortium). Co-last authors (Gaetane Nocturne, Anna Niarakis). Corresponding author details: Anna Niarakis, Postal address: Center for Integrative Biology, 165 Rue Marianne Grunberg-Manago, 31400 Toulouse.

## Abstract

**Objectives:** Sjögren’s disease (SjD) remains a major unmet medical challenge, marked by biological complexity, patient heterogeneity, and a lack of curative treatments. To advance the understanding of its pathogenesis and support therapeutic discovery, we developed a comprehensive knowledgebase in the form of a Molecular Interaction Map (MIM).

**Methods:** Differential expression analysis was performed on peripheral blood samples from SjD patients and healthy controls across three datasets: GSE51092 (190 SjD vs 32 controls), UKPSSR (151 SjD vs 29 controls) and PRECISESADS (304 SjD vs 341 controls). Pathway enrichment analyses provided guidance for MIM construction, which was further refined through literature mining to integrate data-driven results with curated knowledge.

**Results:** A total of 1,625 differentially expressed genes (DEGs) were identified: 725 from PRECISESADS, 1,162 from GSE51092, and 239 from UKPSSR, with 25 DEGs shared across all three datasets. Among these, nine common DEGs were associated with interferon signalling, reinforcing experimental evidence pointing to its pivotal role in SjD. Enrichment analyses revealed 146 pathways, 43 of which were successfully incorporated into the MIM. The resulting SjD Map freely accessible at https://sjdmap.elixir-luxembourg.org/, comprises 829 molecular entities connected by 598 interactions, with 45% of the information depicted supported by transcriptomic data and 47% derived from literature. The map also includes overlays of experimental data and clinical trial information.

**Conclusion:** This first comprehensive Sjögren’s Disease Map, developed through a hybrid data-driven and literature-based approach, offers an integrative view of SjD pathogenesis. It supports visualisation of mechanistic pathways, omics-based data overlays, enables incorporation of user data and drug queries.

Graphical abstract

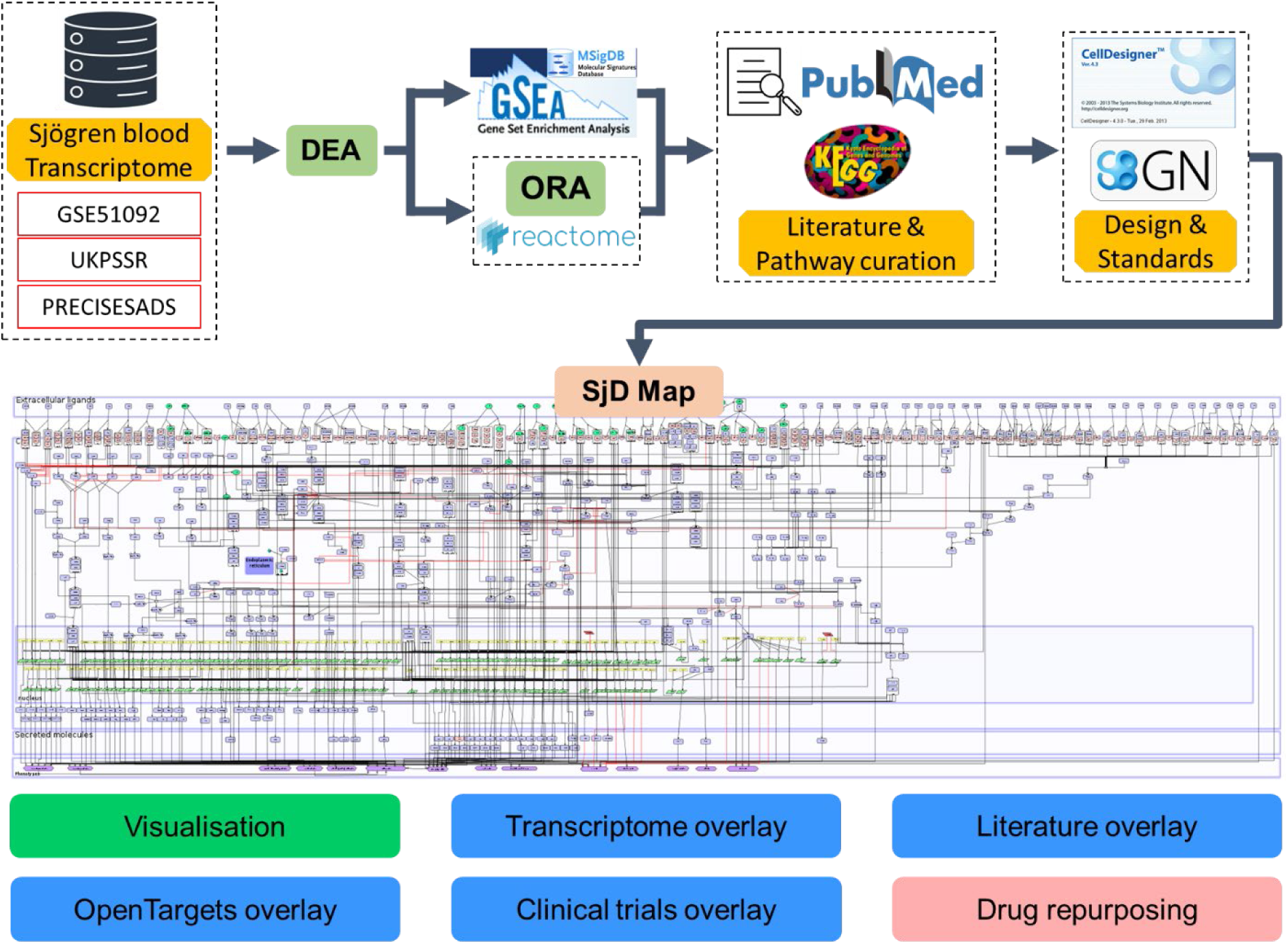

**Key messages:**

- This is the first molecular interaction map specific of Sjögren’s Disease
- It enables comprehensive visualisation of affected pathways through integrated transcriptomic and literature-based evidence, following system’s biology graphical notation schemes
- The map may support therapeutic discovery by linking molecular mechanisms to clinical data and drug targets

## Introduction

Sjögrens Disease (SjD) is a prototypic systemic autoimmune disorder that significantly impairs patients quality of life. This chronic condition affects 0.05 −0.1% of the population, with a striking sex disparity: women are 9 to 20 times more likely to develop the disease (1,2). Patients primarily experience a triad of symptoms: musculo-articular pain, dryness and fatigue. In addition, 30 to 50% of the patients develop systemic complications, with 5-10% of them developing lymphoma, the most severe complication (3). No treatment has been approved until now in SjD (4).

In recent years, systems biology and network medicine have emerged as powerful approaches to reduce biological complexity and extract meaningful insights from large-scale datasets. Traditional statistical methods such as Gene Set Enrichment Analysis (GSEA) or Overrepresentation Analysis (ORA) are widely used to infer biological significance from differentially expressed genes (DEGs), typically by comparing them to curated pathway databases (5). Resources such as KEGG (6), Wikipathways (7), Reactome (8), and pantherDB (9) have played a central role in enabling pathway-level interpretation of omics data, offering mechanistic detail and visual context.

Such bioinformatics approaches have already been applied to SjD aiming to decipher its underlying pathogenesis, identify biomarkers and suggest potential drug targets. However, most of these studies were limited in scope, primarily focusing on differential expression, pathway enrichment, and protein-protein interactions analysis, without addressing the broader mechanistic understanding of the disease (10–13).

To address the limitations of conventional approaches, systems biology has increasingly embraced integrative methods that combine heterogeneous data sources, including literature-based knowledge and high-throughput experimental data, within formalized, standardized frameworks. This holistic strategy has led to the development of several “disease map” projects (14), centred around Molecular Interaction Maps (MIMs) that represent disease mechanisms as a whole. Notable examples include the Covid-19 map (15), the Parkinson’s map (16), the Atlas of Cancer Signalling Network (17) or more recently the Rheumatoid Arthritis Atlas (18,19). These initiatives facilitate a deeper understanding of disease pathogenesis and offer a structured platform for therapeutic hypothesis generation.

Such maps serve as disease-specific knowledge repositories that are both human- and machine-readable, enabling comprehensive exploration and integration of molecular mechanisms. To ensure interoperability, reproducibility, and standardization, these resources are often developed using the Systems Biology Graphical Notation (SBGN) standard (20) which formalizes the representation and exchange of biological networks across computational tools and research groups.

In this study, we aimed to build the first fully standardized MIM for SjD, properly curated and annotated, available at https://sjdmap.elixir-luxembourg.org/ via a web hosting platform to allow for easy navigation and significant zoom in capabilities. The SjD map, hosted on MINERVA platform, ensures accessibility to all dedicated Sjogren-specific content for all users (21). It offers an integrative framework that incorporates mechanistic insights from peer-reviewed literature, transcriptomic datasets, and relevant clinical information. Its primary aim is to bridge the existing knowledge gap between disciplines and facilitate cross-domain exploration of SjD pathogenesis. In this work, we highlighted the different uses and applications of MIMs, enabling both the visualisation of pathways involved in specific manifestations of SjD (e.g., lymphoma), and the identification of relevant biomarkers, thereby paving the way for future targeted therapeutic strategies.

## Methods

### Datasets

We sought to use 3 different whole blood SjD specific datasets in our study. We used GSE51092, a publicly available microarray dataset on GEO, containing samples of 190 SjD patients and 32 controls (22). The series matrix file was downloaded for downstream analysis. The 2 other datasets were available upon request via the NECESSITY consortium. UKPSSR, a microarray dataset of 151 SjD patients and 29 controls was sourced from the United Kingdom Primary Sjögrens Syndrome Registry (23). PRECISESADS, an RNA-Seq study for 304 SjD patients and 341 controls. More technical details about data collection and processing are available in PRECISESADS main article (24).

### Statistical analyses

Differential expression analysis (DEA) was performed on R v4.3 using limma pipeline (25). STRING database was used to assess interaction relationships among intersecting genes and construct a protein-protein interaction (PPI) network (26). Gene set enrichment analysis was performed using the t-test statistic to rank DEGs on the Hallmark pathways reference dataset (H) (27). Overrepresentation Analysis on Reactome was performed by inputting significant total DEGs (FDR adjusted p-value <0.05 and logFC <-0.25 or >0.25) in the “Analyze gene list” of the pathway analyzer present on their web-based server with the options “Project to human” validated. Reactome overrepresented pathways were then filtered using adjusted p-value (FDR <0.05) and excluded disease maps. Significant pathways were compared between techniques and shared identified biological species were selected for integration to the map.

### Map construction process, standards and annotations

We used a framework for building disease maps based on a hybrid approach, combining literature, transcriptomic and pathways analyses inspired by the RA-map and RA-atlas (18,19). Significant pathways identified by both data-driven statistical approaches (GSEA and ORA) and literature curation were integrated in the map. The map was developed using the software CellDesigner (version 4.4.2) (28). Detailed information on the map construction process, literature curation, the use of Systems Biology standards for the pathway representation and the annotation encoding is available in Supplementary Data S1.

### Visualisation and Overlays

SjD map is available as an online interactive map on MINERVA (21). We provide eight downloadable data overlays that can be visualized directly on the map. These overlays offer insights into transcriptomic coverage, literature-based annotations, and clinical trial information related to SjD, as detailed in Supplementary Data S1.

### Ethics

As each dataset has already been published, written informed consent was obtained from all participants in accordance with the requirements of the institutional review board or ethics committee overseeing each respective study.

## Results

### Identification of signalling pathways implicated in Sjögren’s disease pathogenesis via a data-driven approach

1) DEG analysis in blood samples from SjD patients.

In the first step of our workflow, based on the analysis of the three different blood transcriptomic datasets, we identified 1,162 DEGs in GSE51092, 239 in UKPSSR, and 774 in PRECISESADS (Supplementary Data S2). The volcano plots in Figure 1 illustrate the dispersion of genes, revealing that although the range of log fold change (logFC) is similar across studies (approximately −2 to 4) the datasets differ in terms of statistical significance. GSE51092 and UKPSSR, both microarray-based datasets, exhibit comparable maximum adjusted p-values (around 10-10), whereas the RNA-seq dataset PRECISESADS shows a much higher significance, with adjusted p-values approaching 10-80. This greater statistical power can be attributed to the more advanced transcriptomic technology (RNA-seq) and the larger cohort size in the PRECISESADS study (see Methods).

**Figure 1:**
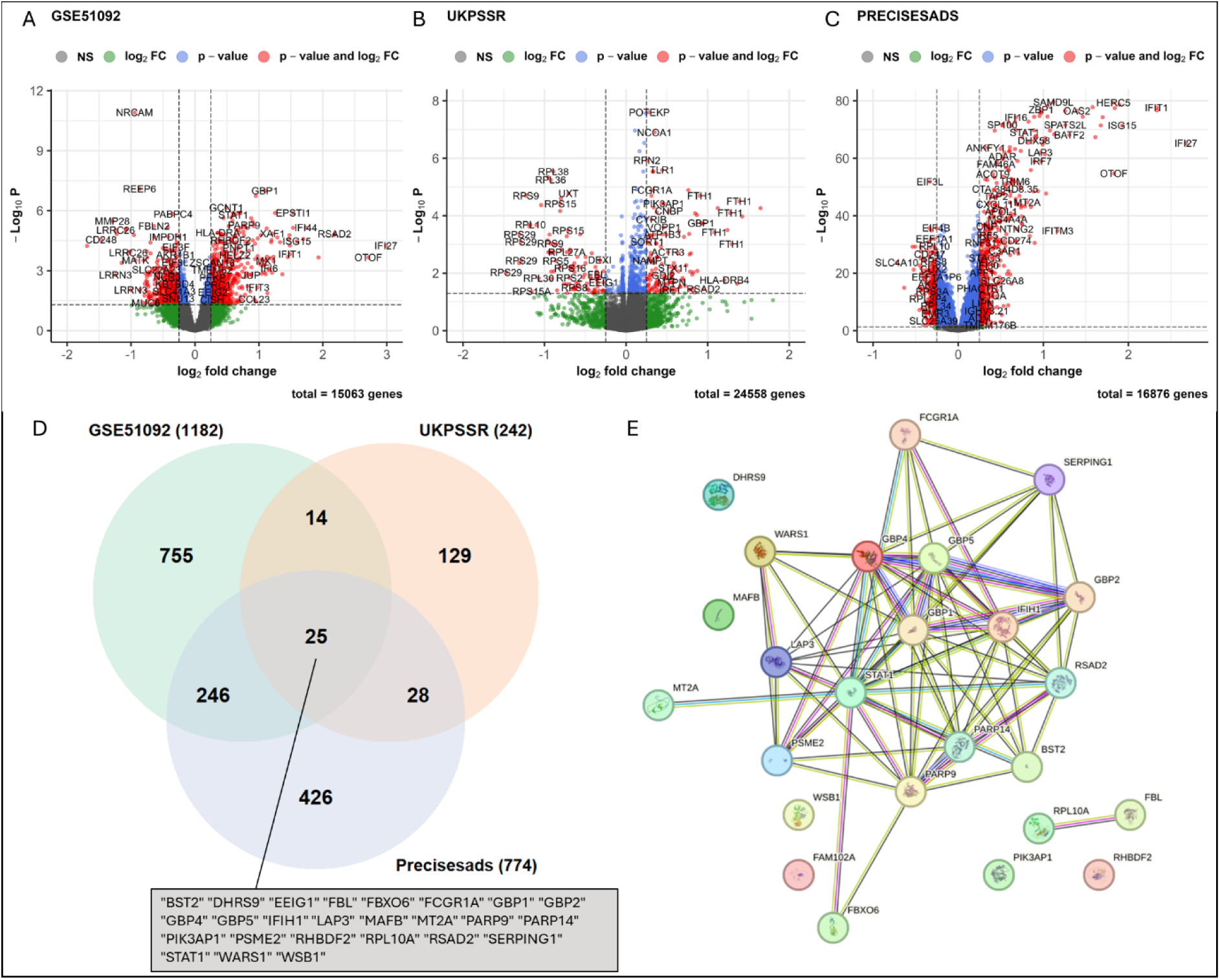
Gene expression changes in SjD. Volcano plots showing in (A) DEGs of SjD patients vs. controls in GSE51092 (microarray data), (B) DEGs of SjD patients vs. controls in UKPSSR (microarray data), (C) DEGs of SjD patients vs. controls in PRECISESADS (RNA-Seq data), (D) Venn diagram showing the common DEGs of the above mentioned three comparisons. (E) Network Analysis of the 25 intersected genes performed on STRINGDB

Regarding expression patterns, GSE51092 revealed 609 upregulated and 553 downregulated genes, UKPSSR showed 140 upregulated and 99 downregulated genes, while PRECISESADS exhibited 526 upregulated and 248 downregulated genes. A comparative analysis of the three datasets revealed 25 commonly upregulated genes across all studies (Figure 1D). Most of these genes, located in the upper-right quadrant of the volcano plots (Figure 1A, B and C), are interferon-related genes, including for instance STAT1 and interferon stimulated genes (ISGs) such as IRF7, IFI44 and IFIT1. A network analysis using STRINGdb showed significant interactions among 17 of the 25 intersecting genes, further emphasizing the involvement of interferon signalling in these datasets (Figure 1E).

2) Pathway enrichment analysis.

Following the DEA, pathway enrichment analysis was conducted using two different statistical techniques namely GSEA and ORA. The GSEA identified a total of 15, 29 and 22 enriched pathways for GSE51092, UKPSSR and PRECISESADS respectively. The Top 15 enriched pathways in GSEA are shown in Figure 2A, B and C, displaying a significant increase in pathways related to inflammation. The first two enriched pathways that are upregulated and common to all datasets are Interferon alpha and Interferon gamma pathways with around 0.6 for the GeneRatio, translated to 60% of common enriched genes between our study and the reference dataset. In the Venn Diagram in Figure 2D, we observed 12 enriched pathways common to all datasets with a positive normalized enrichment score. Apart from Mitotic Spindle, G2M checkpoint, E2F targets and Apoptosis, all eight pathways left are related to inflammation (Figure 2E).

**Figure 2:**
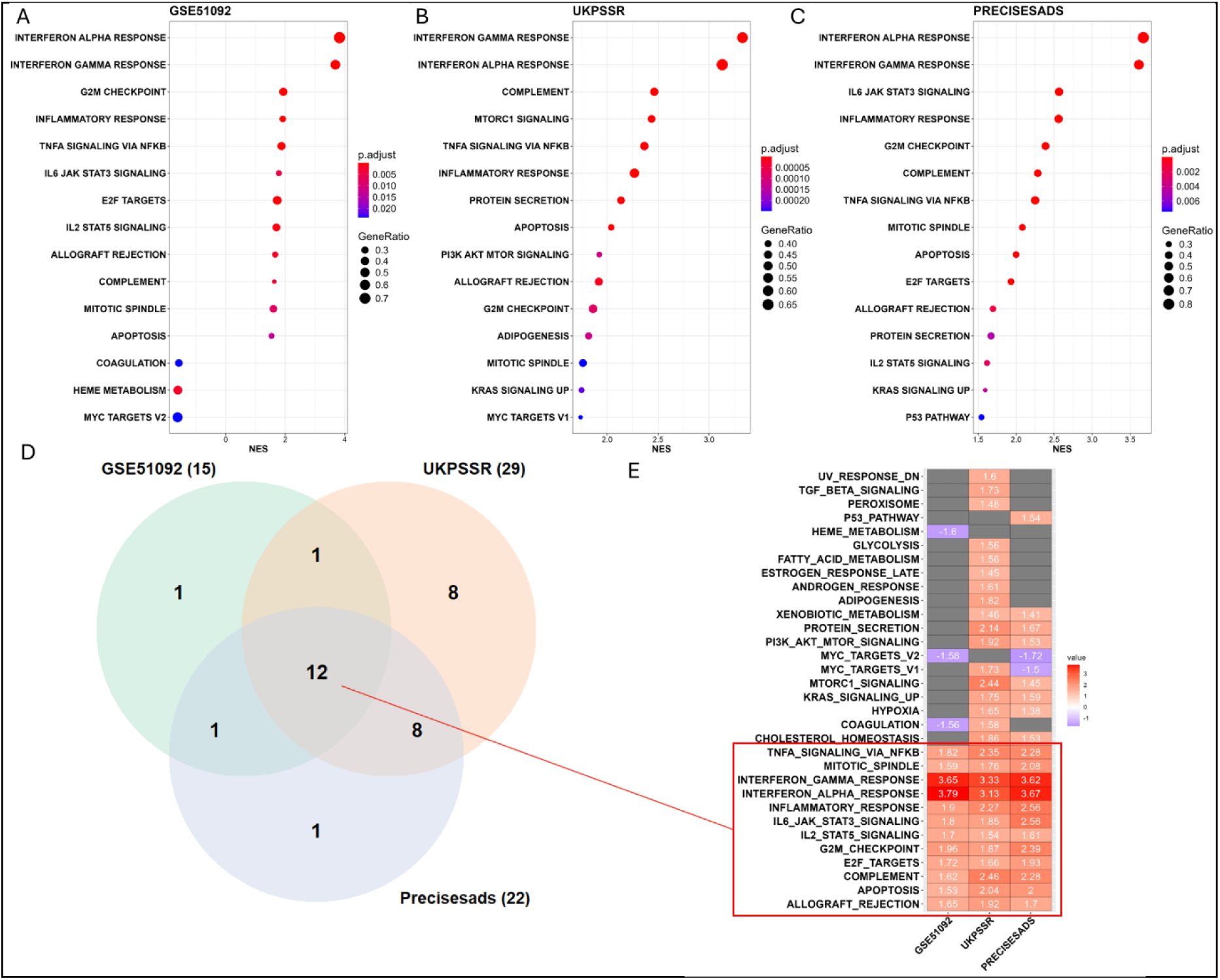
Enrichment Analyses of Sjögren Blood Transcriptome. Dotplots showing in (A) Enrichment GSEA of GSE51092 (B) Enrichment GSEA of UKPSSR (C) Enrichment GSEA of PRECISESADS (D) Venn diagram showing the common enriched pathways (E) Heatmap displaying normalized enrichment scores of the pathways enriched. In the red box are indicated the common enriched pathways to all 3 datasets.

The ORA using the Reactome knowledge base and the cumulative DEG from SjD blood transcriptomic data identified 137 enriched pathways. Of these, 43 pathways were retained based on mechanistic relevance, supported by expert validation and alignment with GSEA results (See Methods). The top 10 pathways of this list are presented in Table 1, while the remaining pathways, along with the Reactome report, are provided as Supplementary Data S3 and S4. The pathways highlight immune system-related processes, emphasizing the pivotal role of immune cells in the pathogenesis of SjD. In particular, the analysis indicates a significant involvement of the type I and type II interferon pathways with nearly half of the entities represented in each case, as well as signalling mechanisms related to macrophages and B cells. These 43 statistically and biologically significant mechanistic pathways were subsequently utilized to construct and enrich the SjD Map.

**Table 1:** TOP10 overrepresented and curated pathways on Reactome.

| Pathway identifier | Pathway name | #Entities found | #Entities total | Entities FDR |
| --- | --- | --- | --- | --- |
| R-HSA-877300 | Interferon gamma signalling | 87 | 177 | 1.35E-14 |
| R-HSA-909733 | Interferon alpha/beta signalling | 82 | 129 | 1.35E-14 |
| R-HSA-198933 | Immunoregulatory interactions between a Lymphoid and a non-Lymphoid cell | 83 | 249 | 1.99E-12 |
| R-HSA-2029481 | FCGR activation | 44 | 103 | 1.30E-09 |
| R-HSA-2029485 | Role of phospholipids in phagocytosis | 48 | 129 | 1.30E-08 |
| R-HSA-5690714 | CD22 mediated BCR regulation | 33 | 72 | 5.46E-08 |
| R-HSA-2730905 | Role of LAT2/NTAL/LAB on calcium mobilization | 41 | 107 | 8.98E-08 |
| R-HSA-2029482 | Regulation of actin dynamics for phagocytic cup formation | 52 | 158 | 1.09E-07 |
| R-HSA-2871809 | FCERI mediated Ca+2 mobilization | 44 | 129 | 5.38E-07 |
| R-HSA-2029480 | Fcgamma receptor (FCGR) dependent phagocytosis | 56 | 193 | 1.45E-06 |

### Assembly of the identified signalling pathways into the SjD Map

The SjD map depicts hallmark signalling pathways, gene regulation, molecular mechanisms and phenotypes involved in SjD pathogenesis. The MIM is constructed in CellDesigner allowing the building of a formalized and standardized map with detailed mechanistic insights into the biological reactions involved (See Methods). The SjD map is compartmentalized in a top to bottom way to represent the flow of information going from the extracellular space (ligands) to the secreted compartments and cellular phenotype section. Figure 3A compares the conventional representation of the interferon signalling pathway commonly found in research articles with its standardized depiction using SBGN-PD within a MIM. The use of SBGN-PD enhances the understanding of the biological processes that individual molecular species undergo. The signalling cascades start after ligand receptor connection, go through the plasma membrane, into the cytosol, to the nucleus (gene regulation) and end up in the secreted compartment (Figure 3B).

**Figure 3:**
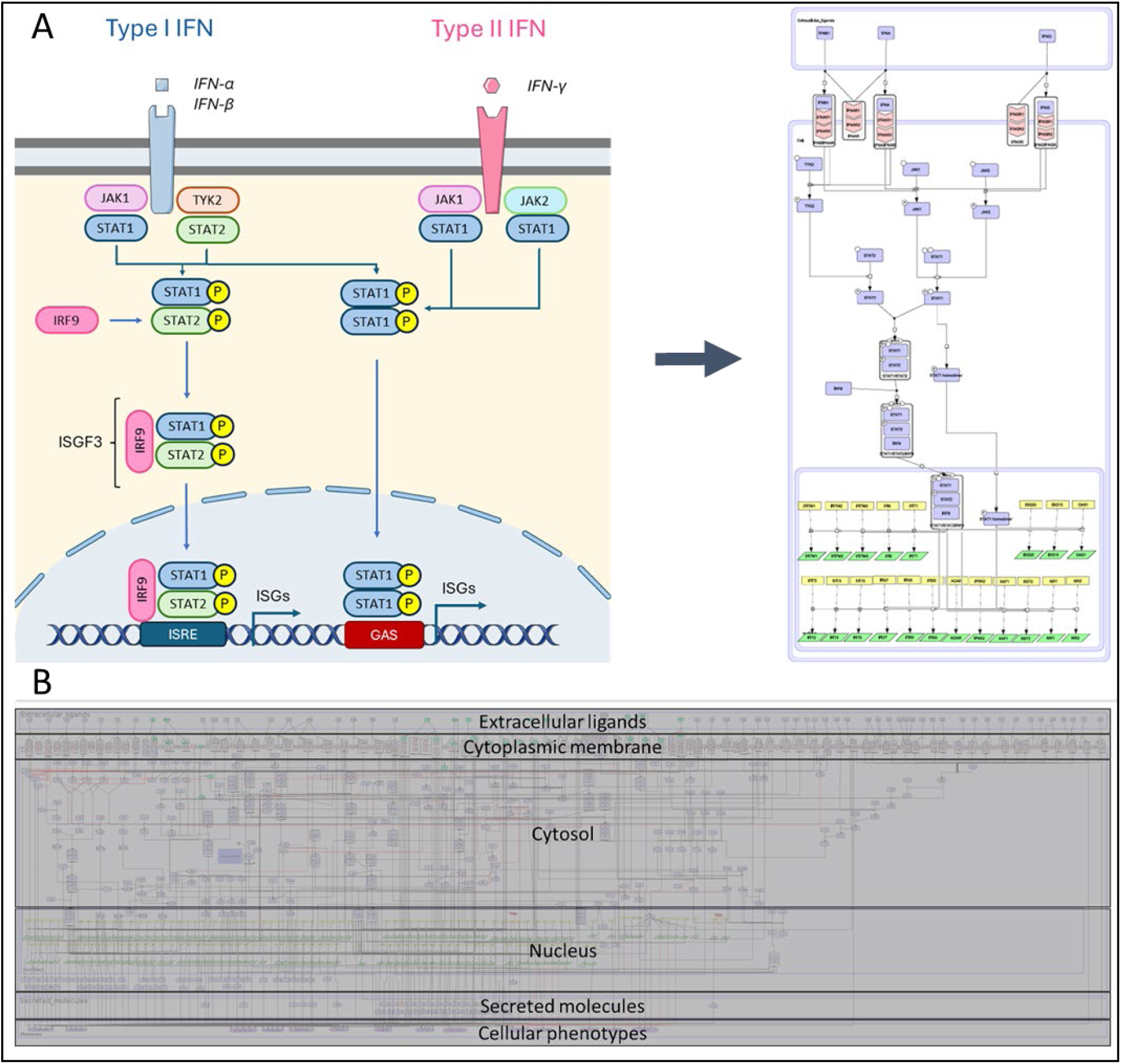
Map Building and Visualisation. (A) Representation of the Interferon signalling pathway in classical scientific articles (left) vs. its depiction in a standardized disease map format (right). This figure was drawn with images from Servier Medical Art (Servier). (B) Overview of the SjD Map. The layout of the map follows the organization of a cell: from the extracellular ligands to the distinct cellular phenotypes.

The SjD map is based on transcriptomic analysis of cohorts of SjD patients, exhaustive literature search, combined with manual pathway curation and data mining. During the pathway assembly, experts of SjD were consulted on data integration, and pathway representation, to ensure accurate interpretation of experimental evidence, and scientific consensus for the disease pathogenesis. The map benefited from several exhaustive reviews and constructive feedback from experts of the Necessity consortium. This thorough revision in every step of its construction led to scrutiny of the depicted mechanisms, both in terms of content and form. The SjD map comprises 829 molecular species interconnected by 598 interactions. The biological entities include 403 proteins, 123 complexes, 133 RNAs, 2 Antisense RNAs, 134 genes, 2 ions and 15 simple molecules such as lipopolysaccharides (LPS). The proteins encompass extracellular, membrane, and cytoplasmic proteins, including signalling intermediates, enzymes, and transcription factors. The 598 interactions between species represent various biological processes, including state transitions, catalysis, inhibition, transport, heterodimer formation, and physical stimulation.

The map contains hallmark cellular and molecular pathways involved in SjD pathogenesis, including the type I and type II interferon signalling (IFN-α and IFN-γ), and a broad array of interleukin-mediated cascades (IL-2, IL-6, IL-7, IL-12, IL-15, IL-21), primarily converging on the JAK-STAT pathway. Additionally, both canonical and non-canonical NF-κB pathways (activated by BAFF, APRIL, BCR, and CD40) are represented, along with Toll-like receptor (TLR) signalling and diverse chemokine axes (CXCL/CCL).

Finally, these signalling cascades converge to drive specific cellular outcomes, informed by both literature and pathway databases, and categorized into fourteen distinct phenotypes: MHC Class I Activation, MHC Class II Activation, T Cell Activation/Differentiation, B Cell Activation/Survival, Cell Proliferation/Survival, Inflammation, Chemotaxis/Infiltration, Angiogenesis, Lymphoid Organ Development, Apoptosis, Regulated Necrosis, Matrix Degradation, Fibrosis, and Phagocytosis. These phenotypes represent the end point of several converged pathways and grouped biomarkers, and can be seen as cell fate decisions.

### Validity of the SjD Map

The confidence in the relevance of a biological entity in SjD pathogenesis increases with the number of supporting publications or gene expression data linked to that entity. The SjD map incorporates 216 PubMed references, distributed across the map and accounting for 57% of its content (Figure 4A, E). Of the biological entities, 13% are associated with a single PubMed reference, while 32% are linked to two to five references. Furthermore, 45% of the map is covered by blood transcriptomic data from Sjögren s patients, used in the construction of the map (Figure 4B). However, some components in the map lack direct associations with published studies or enrichment from the blood dataset. In many cases, these are small, simple molecules such as Phosphatidylinositol-3-4,5-bisphosphate (PI(3,4,5)P3) or Lipopolysaccharide (LPS), which are less frequently studied in SjD pathology compared to their receptors. Other unannotated components function as intermediates, included to ensure the complete visualisation of signalling pathways, offering a more comprehensive representation of the disease mechanism, and enhancing the connectivity of the underlying process description diagram.

**Figure 4:**
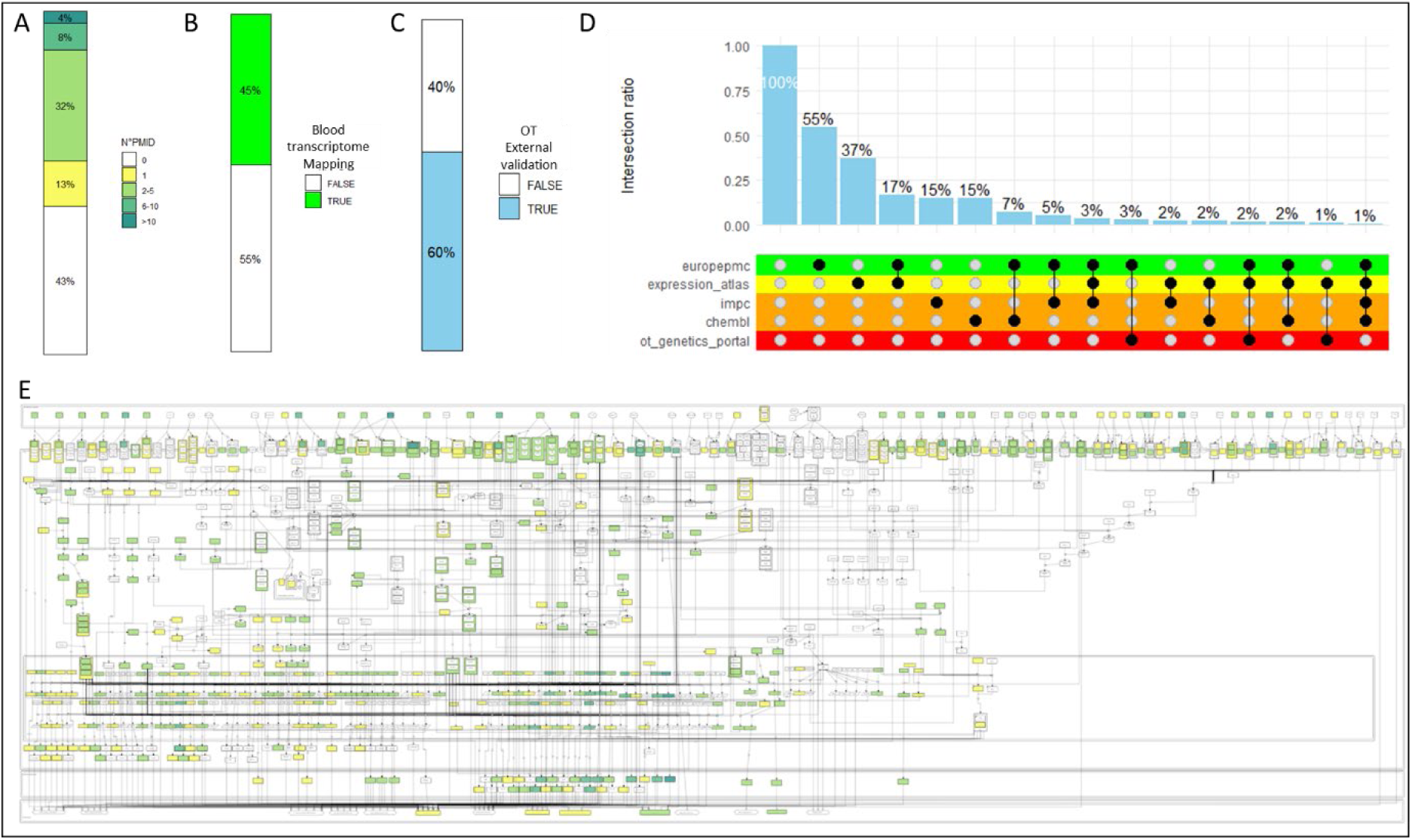
SjD Map annotation score and external validation. Barplots representing (A) the quantification of literature data. The number of publications associated with an entity by categories in white, yellow, light green and dark green respectively corresponding to 0, 1-5, 6-10, and more than 11 PMIDs articles; (B) the quantification of SjD blood transcriptome; and (C) Barplot representing the Coverage of Sjögren OpenTargets database information on the SjD Map. (D) Upset plot representing the main sources retrieved in OpenTargets as an external validation database and their intersections. Each category is color-coded consistently with the OpenTargets_Validation overlay available on the SjD map (E) Overlay of Literature data on the SjD map. Each component is coloured according to its annotation.

To assess the relevance of the SjD map, we also used the OpenTargets database, which integrates data from various sources such as literature, genetic variants, and gene expression related to SjD. We found that 60% of the map was covered by this external database (Figure 4C). Most of the matching content was derived from literature data (Europe PMC), accounting for 55%, followed by gene expression data (Expression Atlas) at 37%, with a 17% overlap between the two (Figure 4D).

The important coverage indicates that our SjD map successfully captures the most relevant molecular entities associated with the disease, with the added value of the mechanistic representation.

### Applications

The SjD map is structured to support integrative analysis of SjD experimental and clinical data through its standardized and mechanistic molecular representation.

A key feature of the map is its capacity to contextualize transcriptomic data by allowing differential expression overlays directly onto the molecular pathways, providing a systems-level view of pathway regulation.

As an example of application shown in Figure 5A, we projected the DEGs between SjD patients who developed lymphoma and those who did not (See Methods) onto the SjD map (DEGs are given in Supplementary Data S5). This overlay notably highlights, for example, the upregulation (in red) of BTK and APRIL (TNFSF13) two genes recently shown to be involved in lymphomagenesis in SjD patients (29).

**Figure 5:**
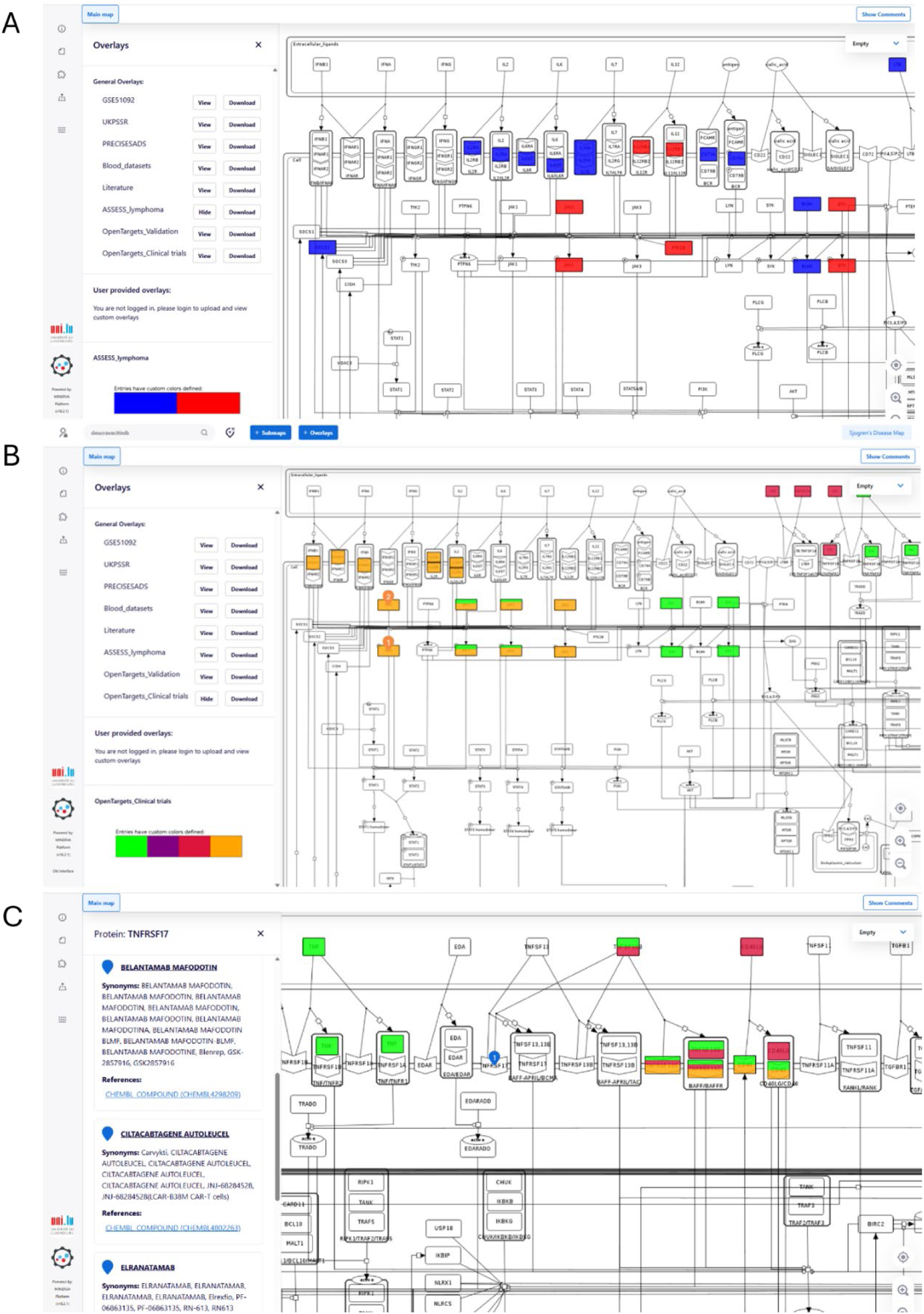
Functionality of the SjD Map. (A) ASSESS lymphoma Differential Expression analysis overlay; Overexpressed species in red, underexpressed species in blue. (B) Drug targets search and overlay for SjD with different colours for the state of the clinical trial: Completed in Green, Not yet recruiting/Active not recruiting/Unknown status in Purple, Withdrawn or terminated in Red, Recruiting in orange; (C) Results of the search for Drug compounds targeting TNFSF17 (BCMA).

In addition to transcriptomic projections, the SjD map allows users to explore drug-target interactions. For instance, users can visualize biological targets of drugs currently being tested in clinical trials for SjD via a combination of DrugBank (31) and a specific overlay. In Figure 5B, we show the result of the research for “Deucravacitinib” TYK2-selective JAK inhibitor. The map highlights TYK2 as the molecular target of this drug. The “Clinical trials overlay” feature provides further insights by showing the clinical trial status of specific drug targets. Deucravacitinib is shown in orange, indicating that the trial is still recruiting (ClinicalTrials.gov Identifier: NCT05946941). This Phase III trial, sponsored by Bristol Myers Squibb, was initiated in 2023 and is expected to report efficacy results by the end of 2026. In contrast, terminated or withdrawn trials are marked in red, such as those targeting Lymphotoxin A/B, suggesting that these pathways may have diminished clinical interest, helping prioritize more promising therapeutic avenues for this disease. Components displayed in green are of particular interest because their clinical trials have been completed, and results have already been posted.

Additionally, reverse searches are possible for a drug repurposing approach: by selecting a protein on the map, users can view all drugs that target it, which could inform future in vitro or in vivo experiments. For instance, to explore therapeutic strategies aimed at B cell modulation, beyond BAFF and BAFF receptor which have already been clinically tested, users can select TNFSF17 (also known as BCMA) on the map. This reveals several therapies targeting this receptor in other diseases, including Elranatamab (bispecific antibody), Belantamab mafodotin (antibody-drug conjugate), and Ciltacabtagene autoleucel (CAR-T cell therapy). Given the central role of B cells in SjD pathogenesis, such tools may support the identification of promising drug repurposing candidates for experimental validation. Finally, the SjD Map available on the MINERVA platform enables users to create custom overlays, facilitating the visualisation of impacted components or pathways in SjD according to their individual research objectives.

## Discussion

This study describes the first MIM dedicated to SjD. The SjD Map was developed using systems biology approaches and a hybrid framework, employing a concomitant integration of both transcriptomic data analysis and literature curation, building upon schemes described previously (18,19). The map is SjD-specific and can be used as a complementary tool for target identification, mechanistic insights, and drug repurposing, as showcased in the selected applications. Moreover, SjD Map allows the visualisation of pathways involved in clinical manifestations of particular interest in SjD, such as lymphomagenesis. As Sjögren’s disease is a heterogeneous disease, understanding different disease phenotypes is of outmost importance for stratifying patients. The SjD map can also be used as the basis of targeted drug repurposing efforts, as well as for the in-silico prediction of drug effects at both molecular and phenotypic levels (32–35).

The transcriptomic analysis of the three datasets of blood samples, primarily identified upregulated interferon-related genes in SjD patients. Subsequent enrichment analyses confirmed the interferon signatures and further highlighted inflammatory processes. Our findings align with previous studies, which have consistently identified strong inflammatory signatures, particularly an interferon type 1 and 2 response, across various omics levels in both salivary gland cells and immune cells, reinforcing the central role of interferon pathways in SjD pathophysiology (10–13). However, despite advances in data analysis, many previous studies fail to integrate mechanistic details, and prior knowledge. The SjD Map is an effort to combine omics data, clinical features, and mechanistic representations in an easy-to-use web-accessible platform.

Existing resources such as STRINGDB (26) and other PPI databases provide valuable interaction networks, yet they often fail to capture the mechanistic insights underlying biological processes or diseases. In contrast, most existing mechanistic pathway databases often lack either disease specificity, proper annotations, or referenced literature. The SjD Map addresses this gap by incorporating hallmark disease pathways through extensive manual curation, rigorous omics data analysis, and expert validation. As part of the disease map community (14), the building of the SjD map has followed standardized methodologies aiming to synthesize and visualize complex biological interactions in a disease-specific context.

Subjecting the map to omics data projection, we analysed potential DEGs driving lymphomagenesis and notably found BTK and APRIL (TNFSF13), previously demonstrated as such by Duret et al. (29). The MIM thus provides a valuable tool for visualizing the molecular heterogeneity of SjD. For instance, the identification of upregulated pathways in patients who develop lymphoma may pave the way for specific therapeutic strategies or the development of predictive biomarkers for this serious complication, both of which represent major clinical challenges. Given the broad heterogeneity of the disease, the potential applications are numerous. The MIM could serve as an in-silico pre-screening platform to explore pathophysiological hypotheses related to specific, and sometimes rare, disease complications.

Additionally, overlaying clinical trials on the map allowed us to visualize drugs tested in completed SjD clinical trials, including well-known inhibitors such as filgotinib (JAK3 inhibitor), tirabrutinib (BTK inhibitor), and lanraplenib (SYK inhibitor). While most of these trials yielded negative outcomes, the case of filgotinib illustrates the map’s potential to support refined therapeutic hypotheses: although initially unsuccessful, filgotinib was later reconsidered for use in a molecularly defined subgroup of patients exhibiting an interferon signature (36). Notably, the SjD map allows the integration of transcriptomic signatures with clinical trial metadata, enabling users to explore associations between molecular phenotypes and therapeutic responses. These overlays enhance the clinical relevance of the SjD Map, making it useful for both educational purposes and in-depth drug discovery efforts.

SjD involves multiple biological processes and affects various cell types, particularly salivary gland epithelial cells and both blood and infiltrating immune cells. In this study, we conducted a literature curation to enrich the map with both blood and gland-related pathways. However, our transcriptomic analysis focused exclusively on blood datasets, as they are more relevant for identifying biomarkers. Although integrating omic data from salivary gland tissues could enhance the maps biological relevance, obtaining clean and robust gland datasets remains challenging due to heterogeneity in term of methods and samples. Addressing this limitation was one of the primary motivations behind the development of ADEx (https://adex.genyo.es/), a web-based application designed to facilitate the integration and exploration of autoimmune disease datasets (37). Future studies should aim to incorporate this data to further refine the map.

Moreover, although the SjD Map is currently designed to represent a single-cell context, it can be dissociated to illustrate the interactions between multiple cell types, thereby improving its utility in modelling the complex cellular interplay characteristic of SjD. Inspired by the RA-Atlas (19), this flexibility allows for a more nuanced understanding of the diseases pathophysiology, ultimately contributing to more targeted therapeutic approaches. One possible approach would be to structure the map into two or three compartments, corresponding to blood and glandular environments, with an intermediary compartment representing infiltrating immune cells.

Additionally, while the map was manually curated and enriched, a process that is both rigorous and time-intensive, future iterations could benefit from artificial intelligence algorithms capable of extracting biological insights and network relationships from literature, like INDRA (38) or BioChatter (39). The integration of large language models may further streamline this process and enhance both the efficiency and depth of map development.

Finally, the MIM currently provides a static framework for studying the immunopathology of SjD. Our goal is to go further by developing a dynamic model capable of performing in silico simulations to predict molecular behaviour in response to defined perturbations, such as pathway activation or modulation by therapeutic agents. This would allow for the assessment of downstream effects on molecular phenotypes. The MIM represents the essential foundation for such dynamic modelling, which could open new avenues for understanding and managing SjD.

### Conclusion

This study introduces the Sjögren’s Disease Map, integrating literature and transcriptomic data to identify key pathways, offering clinical relevance, and providing a foundation for future research and targeted therapeutic development.

## Acknowledgements

We would like to thank the NECESSITY consortium for providing access to the transcriptomic data, and the Luxembourg ELIXIR Node for hosting and ensuring the technical maintenance of the online version of the SjD map. We are also grateful to Dr. Sylvain Soliman, researcher at the Lifeware Group, Inria (Saclay–Île-de-France, Palaiseau, France), for his valuable support during the early development of the SjD map.

## Author Contributions

Conceptualisation: AN; Methodology: SESS, AN, GN, XM; Data Acquisition: SESS, JEG, MB, DC, MAR, MB, SN, WFN, XM, GN, AN; Data curation: SESS, AN, GN, XM; Data analysis and visualisation: SESS; Map development: SESS; Writing—original draft preparation: SESS; Writing—review and editing: SESS, JEG, MB, DC, MAR, MB, SN, WFN, XM, GN, AN; Supervision and project administration: AN, GN, XM; Funding: AN All authors have read and agreed to the published version of the manuscript.

## Data availability statement

The SjD Map is freely accessible at https://sjdmap.elixir-luxembourg.org/. All PMIDs of scientific articles used for the map construction are available in the annotation section of the map. All data and code used to generate results, including XML files of the maps, and confidence score calculations, overlays, and gene expression analysis are available on the Zenodo 10.5281/zenodo.15373343 and GitLab repository: https://gitlab.com/genhotel/SjD_Map.

## References

1. Beydon M, Seror R, Le Guern V, Chretien P, Mariette X, Nocturne G. Impact of patient ancestry on heterogeneity of Sjögren’s disease. RMD Open. mars 2023;9(1):e002955.

2. Thurtle E, Grosjean A, Steenackers M, Strege K, Barcelos G, Goswami P. Epidemiology of Sjögren’s: A Systematic Literature Review. Rheumatol Ther. févr 2024;11(1):1–17.

3. Mariette X, Criswell LA. Primary Sjögren’s Syndrome. N Engl J Med. mars 2018;378(10):931–9.

4. Saraux A, Pers JO, Devauchelle-Pensec V. Treatment of primary Sjögren syndrome. Nat Rev Rheumatol. août 2016;12(8):456–71.

5. Huang DW, Sherman BT, Lempicki RA. Bioinformatics enrichment tools: paths toward the comprehensive functional analysis of large gene lists. Nucleic Acids Res. 1 janv 2009;37(1):1-13.

6. Ogata H, Goto S, Sato K, Fujibuchi W, Bono H, Kanehisa M. KEGG: Kyoto Encyclopedia of Genes and Genomes. Nucleic Acids Res. janv 1999;27(1):29–34.

7. Agrawal A, Balcı H, Hanspers K, Coort SL, Martens M, Slenter DN, et al. WikiPathways 2024: next generation pathway database. Nucleic Acids Res. janv 2024;52(D1):D679-89.

8. Fabregat A, Sidiropoulos K, Garapati P, Gillespie M, Hausmann K, Haw R, et al. The Reactome pathway Knowledgebase. Nucleic Acids Res. janv 2016;44(D1):D481–487.

9. Mi H, Muruganujan A, Huang X, Ebert D, Mills C, Guo X, et al. Protocol Update for large-scale genome and gene function analysis with the PANTHER classification system (v.14.0). Nat Protoc. mars 2019;14(3):703-21.

10. Felten R, Ye T, Schleiss C, Schwikowski B, Sibilia J, Monneaux F, et al. Identification of new candidate drugs for primary Sjögren’s syndrome using a drug repurposing transcriptomic approach. Rheumatol Oxf Engl. nov 2023;62(11):3715–23.

11. Trutschel D, Bost P, Mariette X, Bondet V, Llibre A, Posseme C, et al. Variability of Primary Sjögren’s Syndrome Is Driven by Interferon-α and Interferon-α Blood Levels Are Associated With the Class II HLA-DQ Locus. Arthritis Rheumatol Hoboken NJ. déc 2022;74(12):1991–2002.

12. Soret P, Le Dantec C, Desvaux E, Foulquier N, Chassagnol B, Hubert S, et al. A new molecular classification to drive precision treatment strategies in primary Sjögren’s syndrome. Nat Commun. juin 2021;12(1):3523.

13. Horvath S, Nazmul-Hossain ANM, Pollard RPE, Kroese FGM, Vissink A, Kallenberg CGM, et al. Systems analysis of primary Sjögren’s syndrome pathogenesis in salivary glands identifies shared pathways in human and a mouse model. Arthritis Res Ther. nov 2012;14(6):R238.

14. Ostaszewski M, Gebel S, Kuperstein I, Mazein A, Zinovyev A, Dogrusoz U, et al. Community-driven roadmap for integrated disease maps. Brief Bioinform. mars 2019;20(2):659–70.

15. Ostaszewski M, Niarakis A, Mazein A, Kuperstein I, Phair R, Orta-Resendiz A, et al. COVID19 Disease Map, a computational knowledge repository of virus-host interaction mechanisms. Mol Syst Biol. oct 2021;17(10):e10387.

16. Fujita KA, Ostaszewski M, Matsuoka Y, Ghosh S, Glaab E, Trefois C, et al. Integrating pathways of Parkinson’s disease in a molecular interaction map. Mol Neurobiol. févr 2014;49(1):88–102.

17. Kuperstein I, Bonnet E, Nguyen HA, Cohen D, Viara E, Grieco L, et al. Atlas of Cancer Signalling Network: a systems biology resource for integrative analysis of cancer data with Google Maps. Oncogenesis. juill 2015;4(7):e160–e160.

18. Singh V, Kalliolias GD, Ostaszewski M, Veyssiere M, Pilalis E, Gawron P, et al. RA-map: building a state-of-the-art interactive knowledge base for rheumatoid arthritis. Database. janv 2020;2020:baaa017.

19. Zerrouk N, Aghakhani S, Singh V, Augé F, Niarakis A. A Mechanistic Cellular Atlas of the Rheumatic Joint. Front Syst Biol [Internet]. juill 2022 [cité 18 sept 2024];2. Disponible sur: https://www.frontiersin.org/journals/systems-biology/articles/10.3389/fsysb.2022.925791/full

20. Le Novère N, Hucka M, Mi H, Moodie S, Schreiber F, Sorokin A, et al. The Systems Biology Graphical Notation. Nat Biotechnol. août 2009;27(8):735–41.

21. Gawron P, Ostaszewski M, Satagopam V, Gebel S, Mazein A, Kuzma M, et al. MINERVA—a platform for visualization and curation of molecular interaction networks. Npj Syst Biol Appl. sept 2016;2(1):1–6.

22. Lessard CJ, Li H, Adrianto I, Ice JA, Rasmussen A, Grundahl KM, et al. Variants at multiple loci implicated in both innate and adaptive immune responses are associated with Sjögren’s syndrome. Nat Genet. nov 2013;45(11):1284–92.

23. James K, Al-Ali S, Tarn J, Cockell SJ, Gillespie CS, Hindmarsh V, et al. A Transcriptional Signature of Fatigue Derived from Patients with Primary Sjögren’s Syndrome. PLOS ONE. 2015;10(12):e0143970.

24. Barturen G, Babaei S, Català-Moll F, Martínez-Bueno M, Makowska Z, Martorell-Marugán J, et al. Integrative Analysis Reveals a Molecular Stratification of Systemic Autoimmune Diseases. Arthritis Rheumatol Hoboken NJ. juin 2021;73(6):1073–85.

25. Ritchie ME, Phipson B, Wu D, Hu Y, Law CW, Shi W, et al. limma powers differential expression analyses for RNA-sequencing and microarray studies. Nucleic Acids Res. avr 2015;43(7):e47.

26. Szklarczyk D, Kirsch R, Koutrouli M, Nastou K, Mehryary F, Hachilif R, et al. The STRING database in 2023: protein–protein association networks and functional enrichment analyses for any sequenced genome of interest. Nucleic Acids Res. janv 2023;51(D1):D638–46.

27. Subramanian A, Tamayo P, Mootha VK, Mukherjee S, Ebert BL, Gillette MA, et al. Gene set enrichment analysis: a knowledge-based approach for interpreting genome-wide expression profiles. Proc Natl Acad Sci U S A. oct 2005;102(43):15545–50.

28. Funahashi A, Morohashi M, Kitano H, Tanimura N. CellDesigner: a process diagram editor for gene-regulatory and biochemical networks. BIOSILICO. nov 2003;1(5):159-62.

29. Duret PM, Schleiss C, Kawka L, Meyer N, Ye T, Saraux A, et al. Association Between Bruton’s Tyrosine Kinase Gene Overexpression and Risk of Lymphoma in Primary Sjögren’s Syndrome. Arthritis Rheumatol Hoboken NJ. oct 2023;75(10):1798–811.

30. Ochoa D, Hercules A, Carmona M, Suveges D, Baker J, Malangone C, et al. The next-generation Open Targets Platform: reimagined, redesigned, rebuilt. Nucleic Acids Res. janv 2023;51(D1):D1353–9.

31. Wishart DS, Knox C, Guo AC, Shrivastava S, Hassanali M, Stothard P, et al. DrugBank: a comprehensive resource for in silico drug discovery and exploration. Nucleic Acids Res. janv 2006;34(Database issue):D668-672.

32. Zerrouk N, Augé F, Niarakis A. Building a modular and multi-cellular virtual twin of the synovial joint in Rheumatoid Arthritis. Npj Digit Med. 24 déc 2024;7(1):1-15.

33. Singh V, Naldi A, Soliman S, Niarakis A. A large-scale Boolean model of the rheumatoid arthritis fibroblast-like synoviocytes predicts drug synergies in the arthritic joint. Npj Syst Biol Appl. 15 juill 2023;9(1):1-13.

34. Aghakhani S, Soliman S, Niarakis A. Metabolic reprogramming in Rheumatoid Arthritis Synovial Fibroblasts: A hybrid modeling approach. PLOS Comput Biol. 12 déc 2022;18(12):e1010408.

35. Aghakhani S, Silva-Saffar SE, Soliman S, Niarakis A. Hybrid computational modeling highlights reverse warburg effect in breast cancer-associated fibroblasts. Comput Struct Biotechnol J. 20 août 2023;21:4196-206.

36. Price E, Bombardieri M, Kivitz A, Matzkies F, Gurtovaya O, Pechonkina A, et al. Safety and efficacy of filgotinib, lanraplenib and tirabrutinib in Sjögren’s syndrome: a randomized, phase 2, double-blind, placebo-controlled study. Rheumatol Oxf Engl. 28 nov 2022;61(12):4797–808.

37. Martorell-Marugán J, López-Domínguez R, García-Moreno A, Toro-Domínguez D, Villatoro-García JA, Barturen G, et al. A comprehensive database for integrated analysis of omics data in autoimmune diseases. BMC Bioinformatics. 24 juin 2021;22(1):343.

38. Bachman JA, Gyori BM, Sorger PK. Automated assembly of molecular mechanisms at scale from text mining and curated databases. Mol Syst Biol. 9 mai 2023;19(5):e11325.

39. Lobentanzer S, Feng S, Bruderer N, Maier A, Wang C, Baumbach J, et al. A platform for the biomedical application of large language models. Nat Biotechnol. févr 2025;43(2):166–9.

