## Supplementary Data S1 for "The SjD Map: An interactive pathway tour into Sjögren’s disease signalling mechanisms"

#### Authors

#The PRECISESADS Clinical Consortium is composed of the following members:

Lorenzo Beretta1 , Barbara Vigone1 , Jacques-Olivier Pers2 , Alain Saraux2 , Valérie Devauchelle-Pensec2 , Divi Cornec2 , Sandrine Jousse-Joulin2 , Bernard Lauwerys3 , Julie Ducreux3 , Anne-Lise Maudoux3 , Carlos Vasconcelos4 , Ana Tavares4 , Esmeralda Neves4 , Raquel Faria4 , Mariana Brandão4 , Ana Campar4 , António Marinho4 , Fátima Farinha4 , Isabel Almeida4 , Miguel Angel Gonzalez-Gay Montecón5 , Ricardo Blanco Alonso5 , Alfonso Corrales Martinez5 , Ricard Cervera6 , Ignasi Rodríguez-Pintó6 , Gerard Espinosa6 , Rik Lories7 , Ellen De Langhe7 , Nicolas Huzelmann8 , Doreen Belz8 , Torsten Witte9 , Niklas Baerlecken9 , Georg Stummvoll10, Michael Zauner10, Michaela Lehner10, Eduardo Collantes11, Rafaela Ortega-Castro11, Ma Angeles Aguirre-Zamorano11, Alejandro Escudero-Contreras11, Ma Carmen Castro-Villegas11, Norberto Ortego12, María Concepción Fernández Roldán12, Enrique Raya13, Immaculada Jiménez Moleón13, Enrique de Ramon14, Isabel Díaz Quintero14, Pier Luigi Meroni15, Maria Gerosa15, Tommaso Schioppo15, Carolina Artusi15, Carlo Chizzolini16, Aleksandra Zuber16, Donatienne Wynar16, Laszló Kovács17, Attila Balog17, Magdolna Deák17, Márta Bocskai17, Sonja Dulic17, Gabriella Kádár17, Falk Hiepe18, Velia Gerl18, Silvia Thiel18, Manuel Rodriguez Maresca19, Antonio López-Berrio19, Rocío Aguilar-Quesada19, Héctor Navarro-Linares19, and Marta E. Alarcon-Riquelme20.

1 Referral Center for Systemic Autoimmune Diseases, Fondazione IRCCS Ca' Granda Ospedale Maggiore Policlinico di Milano, Italy; 2 Centre Hospitalier Universitaire de Brest, Hospital de la Cavale Blanche, Brest, France; 3 Pôle de pathologies rhumatismales systémiques et inflammatoires, Institut de Recherche Expérimentale et Clinique, Université catholique de Louvain, Brussels, Belgium; 4 Centro Hospitalar do Porto, Portugal; 5 Servicio Cantabro de Salud, Hospital Universitario Marqués de Valdecilla, Santander, Spain; 6 Hospital Clinic I Provicia, Institut d'Investigacions Biomèdiques August Pi i Sunyer, Barcelona, Spain; 7 Katholieke Universiteit Leuven, Belgium; 8 Klinikum der Universitaet zu Koeln, Cologne, Germany; 9 Medizinische Hochschule Hannover, Germany; 10Medical University Vienna, Vienna, Austria; 11Servicio Andaluz de Salud, Hospital Universitario Reina Sofía Córdoba, Spain; 12Servicio Andaluz de Salud, Complejo hospitalario Universitario de Granada (Hospital Universitario San Cecilio), Spain; 13Servicio Andaluz de Salud, Complejo hospitalario Universitario de Granada (Hospital Virgen de las Nieves), Spain; 14Servicio Andaluz de Salud, Hospital Regional Universitario de Málaga, Spain; 15Università degli studi di Milano, Milan, Italy; 16Hospitaux Universitaires de Genève, Switzerland; 17University of Szeged, Szeged, Hungary; 18Charite, Berlin, Germany; 19Andalusian Public Health System Biobank, Granada, Spain; 20Genyo, Center for Genomics and Oncological Research, Pfizer/University of Granada/Andalusian Regional Government, Granada, Spain.

The study was approved by the following ethic committees: Comitato Etico Area 2 (Fondazione IRCCS Ca' Granda Ospedale Maggiore Policlinico di Milano and University of

Milan); approval no. 425bis Nov 19, 2014, and no. 671_2018 Sep 19, 2018; Klinikum der Universitaet zu Koeln, Cologne, Germany. Geschaftsstelle Ethikkommission; Pôle de pathologies rhumatismales systémiques et inflammatoires, Institut de Recherche Expérimentale et Clinique, Université catholique de Louvain, Brussels, Belgium. Comité d`Èthique Hospitalo-Facultaire; University of Szeged, Szeged, Hungary. Csongrad Megyei Kormanyhivatal; Hospital Clinic I Provicia, Institut d'Investigacions Biomèdiques August Pi i Sunyer, Barcelona, Spain. Comité Ética de Investigación Clínica del Hospital Clínic de Barcelona. Hospital Clinic del Barcelona; Servicio Andaluz de Salud, Hospital Universitario Reina Sofía Córdoba, Spain. Comité de Ética e la Investigación de Centro de Granada (CEI – Granada); Centro Hospitalar do Porto, Portugal. Comissao de ética para a Saude – CES do CHP; Centre Hospitalier Universitaire de Brest, Hospital de la Cavale Blanche, Avenue Tanguy Prigent 29609, Brest, France. Comite de Protection des Personnes Ouest VI; Hospitaux Universitaires de Genève, Switzerland. DEAS – Commission Cantonale d'ethique de la recherche Hopitaux universitaires de Geneve; Andalusian Public Health System Biobank, Granada, Spain; Katholieke Universiteit Leuven, Belgium. Commissie Medische Ethiek UZ KU Leuven /Onderzoek; Charite, Berlin, Germany. Ethikkommission; Medizinische Hochschule Hannover, Germany. Ethikkommission. PRECISESADS Study was funded by the Innovative Medicines Initiative of the European Union with grant number 115565 partly supported by the EFPIA Companies (Alarcon-Riquelme).

### Methods

**Map construction process, standards and annotation**

When designing the Map, we applied the Systems Biology Graphical Notation Process Description (SBGN-PD) scheme. The layout of the map was performed according to a top-down flux of information from the cell. Reactome and KEGG pathway database were used to provide mechanistic details to link pathway-related species depicted in research articles and reviews. In addition, HUGO Gene Nomenclature Committee identifiers (HGNC) were used for pathway components. The CellDesigner maps are available in Systems Biology Markup Language (SBML) format. Information regarding species, reactions, and compartments of the map were referenced using MIRIAM (Minimal Information Requested In the Annotation of Models), a standard for annotating and curating computational models and maps. MIRIAM annotations are added in the CellDesigner dedicated section with the preceding "bqbiol: is describedby" term, which is used to define species or relationship between them according to the information source (e.g., PubMed references (PMIDs), DOI, KEGG identifier). When annotations could not be added into the MIRIAM section for technical issues, the pathway corresponding webpage link was added in the notes section of CellDesigner. Annotated maps provide confidence regarding the map specificity and source of information. See figure below.

**Curation Criteria: Literature enrichment**

First, a systematic Pubmed search was performed to extract biological species, reactions and pathway information about SjD. The search used the following keywords and criteria: (Sjögren[tiab] AND Human NOT mouse) AND (pathways OR cytokines OR Signal Transduction); Article type: Research support, Review, Systematic review; [en] articles; From 2010 until 2024. Were excluded and filtered manually from the search: Clinical studies; Drug/ethnopharmacology studies; General rheumatic/autoimmune diseases articles; Genetic association when not Caucasian; Intercellular studies (flow cytometry); Case reports; Microbiota studies; Dry eye/sicca without control studies. Besides this literature retrieval, a more precise search was performed using the "Sjögren" + "names of the biological species" to enhance literature enrichment. Articles focused on systemic autoimmune diseases were also added to enhance pathway/map connectivity.

**Expert advice and feedback**

Due to the amount of information gathered by the study and the complexity of SjD, regular review of the pathways and factors, and thorough feedback was given by SjD experts (XM, GN) to ensure inclusion of hallmark disease pathways. Their advice allowed to filter, refine and construct the network based on the available data, showcasing the importance of prior knowledge integration in disease network inference.

**Overlays**

PRECISESADS, UKPSSR and GSE51092 representing overexpressed genes in red and underexpressed in blue.

Blood_transcriptome overlay that consists of the combination of all PRECISESADS, UKPSSR and GSE51092, in green.

ASSESS, an overlay highlighting species with DEGs in SjD patients who developed lymphoma (overexpressed genes in red and underexpressed in blue) was generated using data from the ASSESS cohort, a French longitudinal study of SjD patients. The associated transcriptomic data are available in the publicly available microarray dataset GSE140161/ASSESS via GEO (29). Patients were classified based on their lymphoma status prior to differential expression analysis using the limma model, as detailed in the main text. Given the limited number of differentially expressed genes identified following stringent multiple testing correction, we opted to retain genes with unadjusted P values $< 0.05$. This approach was justified by our focus on inferring pathway-level relevance rather than identifying individual differentially expressed genes.

A Literature overlay provides the number of publications associated with an entity by categories in white, yellow, light green and dark green respectively corresponding to 0, 1-5, 6-10, and more than 11 PMIDs articles.

Lastly, the map includes two OpenTargets overlays, that were created using the platform's API: OpenTargets_Drugs used for identifying drug targets used in Sjögren's clinical trials with a color code associated to identify the stage of the clinical trial (green = completed; orange = Recruiting; purple = Not yet recruiting, Active not recruiting or unknown status; red = Withdrawn or terminated). OpenTargets_Validation used for enrichment validation with a colour code associated to identify Sjögren's (EFO_0000699) external sources of information (green = europePMC; red = genetic variant associations; yellow = expression Atlas; orange = other databases) (30).
