## Supplementary Data S4 for "The SjD Map: An interactive pathway tour into Sjögren’s disease signalling mechanisms"

### Pathway Analysis Report

This report contains the pathway analysis results for the submitted sample ''. Analysis was performed against Reactome version 86 on 06/12/2023. The web link to these results is:

https://reactome.org/PathwayBrowser/#/ANALYSIS=MjAyMzEyMDYwOTU0MzdfNDc3Nw%3D%3D

Please keep in mind that analysis results are temporarily stored on our server. The storage period depends on usage of the service but is at least 7 days. As a result, please note that this URL is only valid for a limited time period and it might have expired.

### Table of Contents


1. Introduction
2. Properties
3. Genome-wide overview
4. Most significant pathways
5. Pathways details
6. Identifiers found
7. Identifiers not found

### 1. Introduction

Reactome is a curated database of pathways and reactions in human biology. Reactions can be considered as pathway 'steps'. Reactome defines a 'reaction' as any event in biology that changes the state of a biological molecule. Binding, activation, translocation, degradation and classical biochemical events involving a catalyst are all reactions. Information in the database is authored by expert biologists, entered and maintained by Reactome's team of curators and editorial staff. Reactome content frequently cross-references other resources e.g. NCBI, Ensembl, UniProt, KEGG (Gene and Compound), ChEBI, PubMed and GO. Orthologous reactions inferred from annotation for Homo sapiens are available for 14 non-human species including mouse, rat, chicken, puffer fish, worm, fly and yeast. Pathways are represented by simple diagrams following an SBGN-like format.

Reactome's annotated data describe reactions possible if all annotated proteins and small molecules were present and active simultaneously in a cell. By overlaying an experimental dataset on these annotations, a user can perform a pathway over-representation analysis. By overlaying quantitative expression data or time series, a user can visualize the extent of change in affected pathways and its progression. A binomial test is used to calculate the probability shown for each result, and the p-values are corrected for the multiple testing (Benjamini–Hochberg procedure) that arises from evaluating the submitted list of identifiers against every pathway.

To learn more about our Pathway Analysis, please have a look at our relevant publications:


Fabregat A, Sidiropoulos K, Garapati P, Gillespie M, Hausmann K, Haw R, … D'Eustachio P (2016). The reactome pathway knowledgebase. Nucleic Acids Research, 44(D1), D481–D487. https://doi.org/10.1093/nar/gkv1351.

Fabregat A, Sidiropoulos K, Viteri G, Forner O, Marin-Garcia P, Arnau V, … Hermjakob H (2017). Reactome pathway analysis: a high-performance in-memory approach. BMC Bioinformatics, 18.

#### 2. Properties

- This is an **overrepresentation** analysis: A statistical (hypergeometric distribution) test that determines whether certain Reactome pathways are over-represented (enriched) in the submitted data. It answers the question ‘Does my list contain more proteins for pathway X than would be expected by chance?’ This test produces a probability score, which is corrected for false discovery rate using the Benjamani-Hochberg method.
- 1084 out of 1660 identifiers in the sample were found in Reactome, where 1805 pathways were hit by at least one of them.
- All non-human identifiers have been converted to their human equivalent.
- This report is filtered to show only results for species 'Homo sapiens' and resource 'all resources'.
- The unique ID for this analysis (token) is MjAyMzEyMDYwOTU0MzdfNDc3Nw%3D%3D. This ID is valid for at least 7 days in Reactome's server. Use it to access Reactome services with your data.

### 3. Genome-wide overview

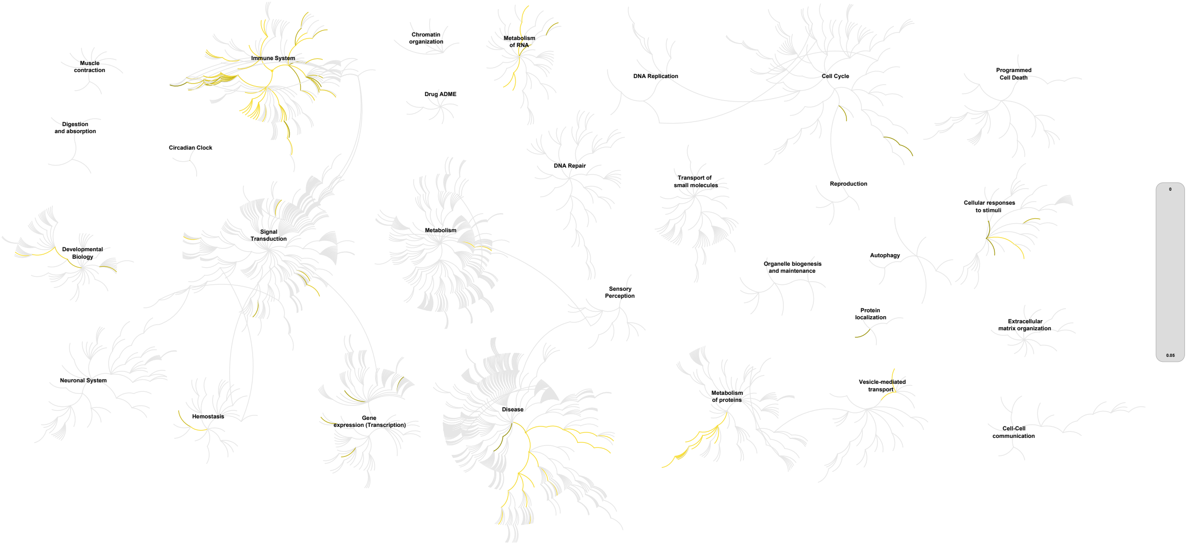


reactome

This figure shows a genome-wide overview of the results of your pathway analysis. Reactome pathways are arranged in a hierarchy. The center of each of the circular "bursts" is the root of one top-level pathway, for example "DNA Repair". Each step away from the center represents the next level lower in the pathway hierarchy. The color code denotes over-representation of that pathway in your input dataset. Light grey signifies pathways which are not significantly over-represented.

### 4. Most significant pathways

The following table shows the 25 most relevant pathways sorted by p-value.

| Pathway name | Entities | | | | Reactions | |
|---|---|---|---|---|---|---|
| | found | ratio | p-value | FDR* | found | ratio |
| Eukaryotic Translation Elongation | 60 / 102 | 0.007 | 1.11e-16 | 1.35e-14 | 9 / 9 | 6.21e-04 |
| Peptide chain elongation | 57 / 97 | 0.006 | 1.11e-16 | 1.35e-14 | 5 / 5 | 3.45e-04 |
| Nonsense Mediated Decay (NMD) independent of the Exon Junction Complex (EJC) | 56 / 101 | 0.007 | 1.11e-16 | 1.35e-14 | 1 / 1 | 6.91e-05 |
| Formation of a pool of free 40S subunits | 57 / 106 | 0.007 | 1.11e-16 | 1.35e-14 | 2 / 2 | 1.38e-04 |
| Nonsense-Mediated Decay (NMD) | 61 / 124 | 0.008 | 1.11e-16 | 1.35e-14 | 6 / 6 | 4.14e-04 |
| Nonsense Mediated Decay (NMD) enhanced by the Exon Junction Complex (EJC) | 61 / 124 | 0.008 | 1.11e-16 | 1.35e-14 | 5 / 5 | 3.45e-04 |
| L13a-mediated translational silencing of Ceruloplasmin expression | 59 / 120 | 0.008 | 1.11e-16 | 1.35e-14 | 3 / 3 | 2.07e-04 |
| GTP hydrolysis and joining of the 60S ribosomal subunit | 58 / 120 | 0.008 | 1.11e-16 | 1.35e-14 | 3 / 3 | 2.07e-04 |
| SRP-dependent cotranslational protein targeting to membrane | 57 / 119 | 0.008 | 1.11e-16 | 1.35e-14 | 5 / 5 | 3.45e-04 |
| Interferon gamma signaling | 87 / 177 | 0.012 | 1.11e-16 | 1.35e-14 | 22 / 23 | 0.002 |
| Interferon alpha/beta signaling | 82 / 129 | 0.008 | 1.11e-16 | 1.35e-14 | 23 / 25 | 0.002 |
| Interferon Signaling | 153 / 398 | 0.026 | 1.11e-16 | 1.35e-14 | 109 / 119 | 0.008 |
| Cytokine Signaling in Immune system | 265 / 1,101 | 0.072 | 1.11e-16 | 1.35e-14 | 524 / 785 | 0.054 |
| Immune System | 525 / 2,661 | 0.173 | 1.11e-16 | 1.35e-14 | 1,091 / 1,704 | 0.118 |
| Eukaryotic Translation Termination | 55 / 106 | 0.007 | 1.11e-16 | 1.35e-14 | 3 / 5 | 3.45e-04 |
| Response of EIF2AK4 (GCN2) to amino acid deficiency | 57 / 115 | 0.007 | 1.11e-16 | 1.35e-14 | 6 / 16 | 0.001 |
| Selenocysteine synthesis | 55 / 112 | 0.007 | 2.22e-16 | 2.55e-14 | 2 / 7 | 4.83e-04 |
| Viral mRNA Translation | 55 / 114 | 0.007 | 4.44e-16 | 4.31e-14 | 2 / 2 | 1.38e-04 |
| Eukaryotic Translation Initiation | 59 / 130 | 0.008 | 4.44e-16 | 4.31e-14 | 14 / 21 | 0.001 |
| Cap-dependent Translation Initiation | 59 / 130 | 0.008 | 4.44e-16 | 4.31e-14 | 11 / 18 | 0.001 |
| Immunoregulatory interactions between a Lymphoid and a non-Lymphoid cell | 83 / 249 | 0.016 | 2.14e-14 | 1.99e-12 | 33 / 44 | 0.003 |
| Regulation of expression of SLITs and ROBOs | 64 / 183 | 0.012 | 2.81e-12 | 2.48e-10 | 5 / 20 | 0.001 |
| Cellular response to starvation | 62 / 176 | 0.011 | 4.58e-12 | 3.89e-10 | 18 / 28 | 0.002 |

| Pathway name | Entities | | | | Reactions | |
|---|---|---|---|---|---|---|
| | found | ratio | p-value | FDR* | found | ratio |
| **SARS-CoV-1 modulates host translation machinery** | 27 / 41 | 0.003 | 1.56e-11 | 1.26e-09 | 2 / 3 | 2.07e-04 |
| **FCGR activation** | 44 / 103 | 0.007 | 1.66e-11 | 1.30e-09 | 6 / 6 | 4.14e-04 |

* False Discovery Rate

### 5. Pathways details

For every pathway of the most significant pathways, we present its diagram, as well as a short summary, its bibliography and the list of inputs found in it.

#### 1. Eukaryotic Translation Elongation (R-HSA-156842)

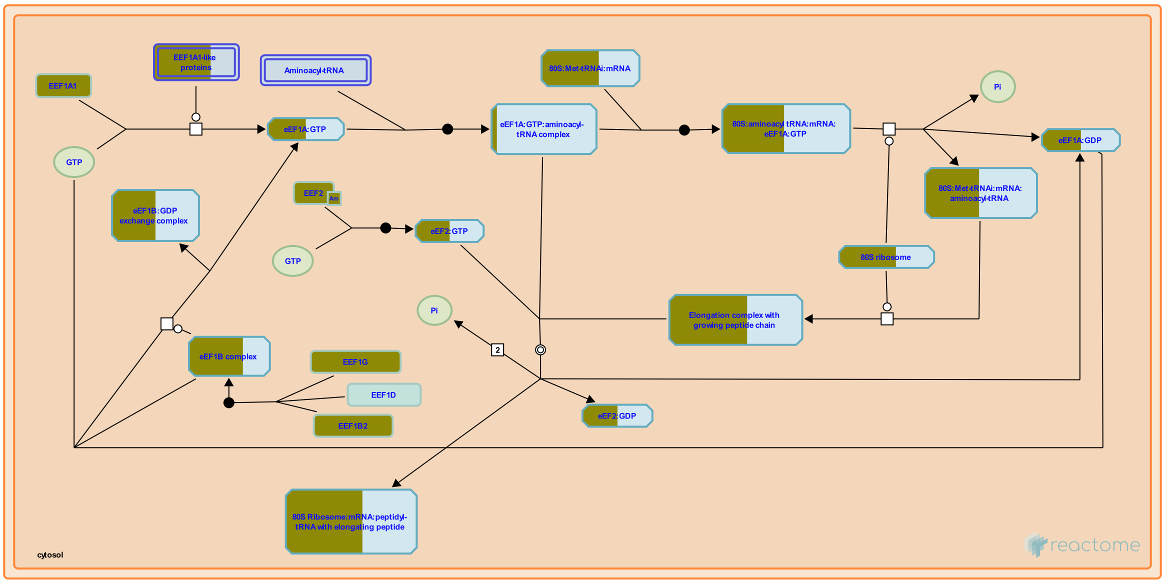


**Cellular compartments:** cytosol.

The translation elongation cycle adds one amino acid at a time to a growing polypeptide according to the sequence of codons found in the mRNA. The next available codon on the mRNA is exposed in the aminoacyl-tRNA (aa-tRNA) binding site (A site) on the 30S subunit.

A: Ternary complexes of aa -tRNA:eEF1A:GTP enter the ribosome and enable the anticodon of the tRNA to make a codon/anticodon interaction with the A-site codon of the mRNA. B: Upon cognate recognition, the eEF1A:GTP is brought into the GTPase activating center of the ribosome, GTP is hydrolyzed and eEF1A:GDP leaves the ribosome. C: The peptidyl transferase center of ribosome catalyses the formation of a peptide bond between the incoming amino acid and the peptide found in the peptidyl-tRNA binding site (P site). D: In the pre-translocation state of the ribosome, the eEF2:GTP enters the ribosome, physically translocating the peptidyl-tRNA out of the A site to P site and leaves the ribosome eEF2:GDP. This action of eEF2:GTP accounts for the precise movement of the mRNA by 3 nucleotides.Consequently, deacylated tRNA is shifted to the E site. A ribosome associated ATPase activity is proposed to stimulate the release of deacylated tRNA from the E site subsequent to translocation (Elskaya et al., 1991). In this post-translocation state, the ribosome is now ready to receive a new ternary complex.

This process is illustrated below with: an amino acyl-tRNA with an amino acid, a peptidyl-tRNA with a growing peptide, a deacylated tRNA with an -OH, and a ribosome with A,P and E sites to accommodate these three forms of tRNA.

##### References

Kapp LD & Lorsch JR (2004). The molecular mechanics of eukaryotic translation. Annu Rev Biochem, 73, 657-704.

#### Edit history

| Date | Action | Author |
|---|---|---|
| 2004-12-14 | Created | Balar B, Ulloque R, Ortiz PA, Pittman YR, Kinzy TG et al. |
| 2005-03-13 | Authored | Gopinathrao G |
| 2023-08-26 | Modified | Wright A |

#### 57 submitted entities found in this pathway, mapping to 71 Reactome entities

| Input | UniProt Id | Input | UniProt Id | Input | UniProt Id |
|---|---|---|---|---|---|
| AGRN | P32969 | EEF1A1 | P68104 | EEF1A1P5 | Q5VTE0 |
| EEF1B2 | P24534 | EEF1G | P26641 | EEF2 | P13639 |
| FAU | P62861 | RPL10 | P27635, Q96L21 | RPL10A | P61313, P62906 |
| RPL13 | P26373, P40429 | RPL13A | P40429, P61313 | RPL15 | P61313 |
| RPL18 | Q07020 | RPL21 | P46778 | RPL27A | P46776, P61353 |
| RPL29 | P47914 | RPL3 | P39023, Q92901 | RPL30 | P62888 |
| RPL31 | P62899 | RPL32 | P62888, P62910 | RPL34 | P49207, P62899 |
| RPL35 | P42766 | RPL35A | P18077 | RPL36 | Q9Y3U8 |
| RPL38 | P63173 | RPL3L | Q92901 | RPL4 | P36578 |
| RPL5 | P46777 | RPL7 | P18124 | RPL7A | P18124, P62424 |
| RPL9P8 | P32969 | RPLP0 | P05388 | RPLP2 | P05387 |
| RPS10 | P46783 | RPS11 | P62280 | RPS12 | P25398 |
| RPS13 | P62277 | RPS14 | P62263 | RPS15 | P62841 |
| RPS15A | P62244 | RPS16 | P62249 | RPS17 | P08708 |
| RPS18 | P62269 | RPS2 | P15880, P46782 | RPS23 | P62266 |
| RPS27 | P42677, Q71UM5 | RPS27A | P42677, P62979, P62987 | RPS29 | P62273 |
| RPS3 | P23396 | RPS3A | P61247 | RPS4X | P62701, Q8TD47 |
| RPS5 | P46782 | RPS6 | P62753 | RPS8 | P62241 |
| RPS9 | P46781 | RPSA | P08865 | SOS2 | P42766 |

#### 2. Peptide chain elongation (R-HSA-156902)

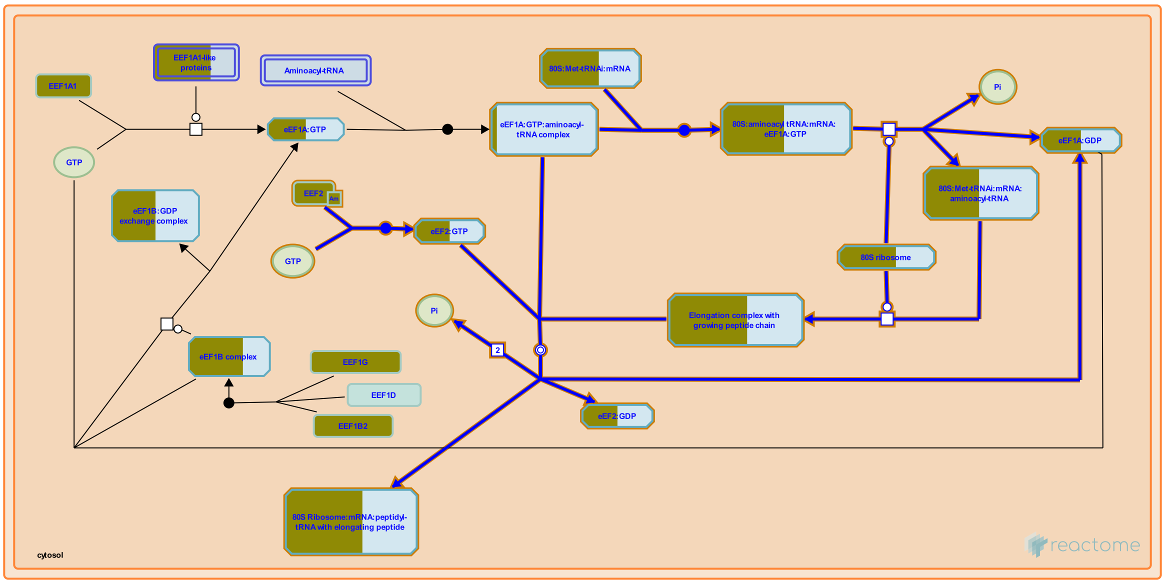


**Cellular compartments:** cytosol.

The mechanism of a peptide bond requires the movement of three protons. First the deprotonation of the ammonium ion generates a reactive amine, allowing a nucleophilic attack on the carbonyl group. This is followed by the loss of a proton from the reaction intermediate, only to be taken up by the oxygen on the leaving group (from the end of the amino acid chain bound to the tRNA in the P-site). The peptide bond formation results in the net loss of one water molecule, leaving a deacylated-tRNA in the P-site, and a nascent polypeptide chain one amino acid larger in the A-site.

For the purpose of illustration, the figures used in the section show one amino acid being added to a peptidyl-tRNA with a growing peptide chain.

##### References


Lorsch JR & Green R (2002). The path to perdition is paved with protons. Cell, 110, 665-8.


##### Edit history

| Date | Action | Author |
|---|---|---|
| 2005-03-13 | Authored | Gopinathrao G |
| 2005-03-17 | Created | Balar B, Ulloque R, Kinzy TG |
| 2023-08-31 | Modified | Wright A |

##### 54 submitted entities found in this pathway, mapping to 68 Reactome entities

| Input | UniProt Id | Input | UniProt Id | Input | UniProt Id |
|---|---|---|---|---|---|
| AGRN | P32969 | EEF1A1 | P68104 | EEF2 | P13639 |
| FAU | P62861 | RPL10 | P27635, Q96L21 | RPL10A | P61313, P62906 |
| RPL13 | P26373, P40429 | RPL13A | P40429, P61313 | RPL15 | P61313 |
| RPL18 | Q07020 | RPL21 | P46778 | RPL27A | P46776, P61353 |
| RPL29 | P47914 | RPL3 | P39023, Q92901 | RPL30 | P62888 |
| RPL31 | P62899 | RPL32 | P62888, P62910 | RPL34 | P49207, P62899 |
| RPL35 | P42766 | RPL35A | P18077 | RPL36 | Q9Y3U8 |

| Input | UniProt Id | Input | UniProt Id | Input | UniProt Id |
|---|---|---|---|---|---|
| RPL38 | P63173 | RPL3L | Q92901 | RPL4 | P36578 |
| RPL5 | P46777 | RPL7 | P18124 | RPL7A | P18124, P62424 |
| RPL9P8 | P32969 | RPLP0 | P05388 | RPLP2 | P05387 |
| RPS10 | P46783 | RPS11 | P62280 | RPS12 | P25398 |
| RPS13 | P62277 | RPS14 | P62263 | RPS15 | P62841 |
| RPS15A | P62244 | RPS16 | P62249 | RPS17 | P08708 |
| RPS18 | P62269 | RPS2 | P15880, P46782 | RPS23 | P62266 |
| RPS27 | P42677, Q71UM5 | RPS27A | P42677, P62979, P62987 | RPS29 | P62273 |
| RPS3 | P23396 | RPS3A | P61247 | RPS4X | P62701, Q8TD47 |
| RPS5 | P46782 | RPS6 | P62753 | RPS8 | P62241 |
| RPS9 | P46781 | RPSA | P08865 | SOS2 | P42766 |

#### 3. Nonsense Mediated Decay (NMD) independent of the Exon Junction Complex (EJC) (R-HSA-975956)

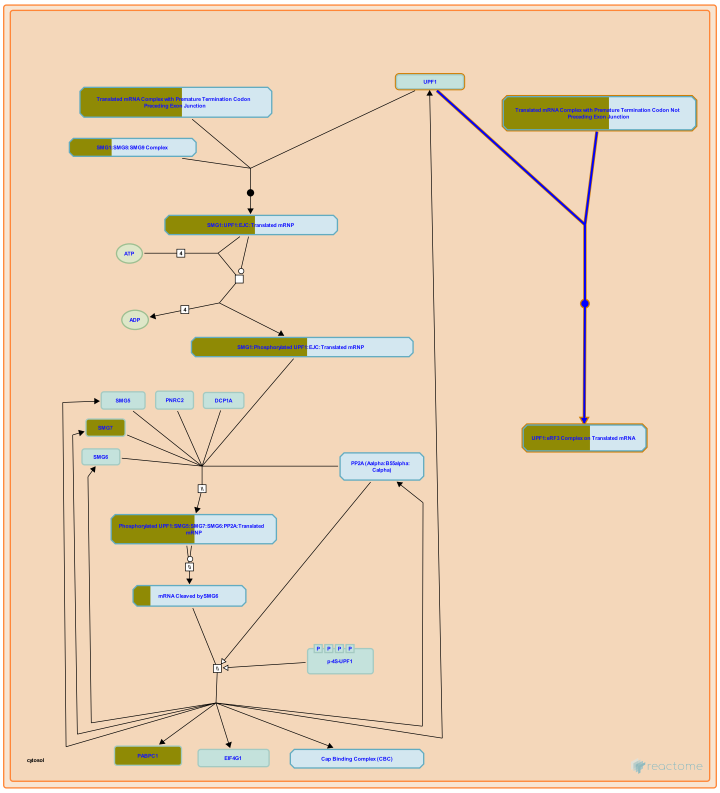


**Cellular compartments:** cytosol.

Nonsense-mediated decay has been observed with mRNAs that do not have an exon junction complex (EJC) downstream of the termination codon (reviewed in Isken and Maquat 2007, Chang et al. 2007, Behm-Ansmant et al. 2007, Rebbapragada and Lykke-Andersen 2009, Nicholson et al. 2010). In these cases the trigger is unknown but a correlation with the length of the 3' UTR has sometimes been seen. The current model posits a competition between PABP and UPF1 for access to eRF3 at the terminating ribosome (Ivanov et al. 2008, Singh et al. 2008, reviewed in Bhuvanagiri et al. 2010). Abnormally long 3' UTRs may prevent PABP from efficiently interacting with eRF3 and allow UPF1 to bind eRF3 instead. Long UTRs with hairpin loops may bring PABP closer to eRF3 and help evade NMD (Eberle et al. 2008).

The pathway of degradation taken during EJC-independent NMD has not been elucidated. It is thought that phosphorylation of UPF1 by SMG1 and recruitment of SMG6 or SMG5 and SMG7 are involved, as with EJC-enhanced NMD, but this has not yet been shown.

##### References


Yepiskoposyan H, Kleinschmidt N, Zamudio Orozco R, Metze S, Muhlemann O & Nicholson P (2010). Nonsense-mediated mRNA decay in human cells: mechanistic insights, functions beyond quality control and the double-life of NMD factors. Cell Mol Life Sci, 67, 677-700.

Stalder L, Eberle AB, Mathys H, Orozco RZ & Muhlemann O (2008). Posttranscriptional gene regulation by spatial rearrangement of the 3' untranslated region. PLoS Biol, 6, e92.

Saulière J, Wittkopp N, Behm-Ansmant I, Izaurralde E, Rehwinkel J & Kashima I (2007). mRNA quality control: an ancient machinery recognizes and degrades mRNAs with nonsense codons. FEBS Lett, 581, 2845-53.

Rebbapragada I, Singh G & Lykke-Andersen J (2008). A competition between stimulators and antagonists of Upf complex recruitment governs human nonsense-mediated mRNA decay. PLoS Biol, 6, e111.

Bhuvanagiri M, Kulozik AE, Hentze MW & Schlitter AM (2010). NMD: RNA biology meets human genetic medicine. Biochem J, 430, 365-77.

#### Edit history

| Date | Action | Author |
|---|---|---|
| 2010-10-08 | Edited | May B |
| 2010-10-08 | Authored | May B |
| 2010-10-11 | Created | May B |
| 2011-05-19 | Reviewed | Neu-Yilik G |
| 2023-08-31 | Modified | Wright A |

#### 53 submitted entities found in this pathway, mapping to 67 Reactome entities

| Input | UniProt Id | Input | UniProt Id | Input | UniProt Id |
|---|---|---|---|---|---|
| AGRN | P32969 | FAU | P62861 | PABPC1 | P11940 |
| RPL10 | P27635, Q96L21 | RPL10A | P61313, P62906 | RPL13 | P26373, P40429 |
| RPL13A | P40429, P61313 | RPL15 | P61313 | RPL18 | Q07020 |
| RPL21 | P46778 | RPL27A | P46776, P61353 | RPL29 | P47914 |
| RPL3 | P39023, Q92901 | RPL30 | P62888 | RPL31 | P62899 |
| RPL32 | P62888, P62910 | RPL34 | P49207, P62899 | RPL35 | P42766 |
| RPL35A | P18077 | RPL36 | Q9Y3U8 | RPL38 | P63173 |
| RPL3L | Q92901 | RPL4 | P36578 | RPL5 | P46777 |
| RPL7 | P18124 | RPL7A | P18124, P62424 | RPL9P8 | P32969 |
| RPLP0 | P05388 | RPLP2 | P05387 | RPS10 | P46783 |
| RPS11 | P62280 | RPS12 | P25398 | RPS13 | P62277 |
| RPS14 | P62263 | RPS15 | P62841 | RPS15A | P62244 |
| RPS16 | P62249 | RPS17 | P08708 | RPS18 | P62269 |
| RPS2 | P15880, P46782 | RPS23 | P62266 | RPS27 | P42677, Q71UM5 |
| RPS27A | P42677, P62979, P62987 | RPS29 | P62273 | RPS3 | P23396 |
| RPS3A | P61247 | RPS4X | P62701, Q8TD47 | RPS5 | P46782 |
| RPS6 | P62753 | RPS8 | P62241 | RPS9 | P46781 |
| RPSA | P08865 | SOS2 | P42766 | | |

#### 4. Formation of a pool of free 40S subunits (R-HSA-72689)

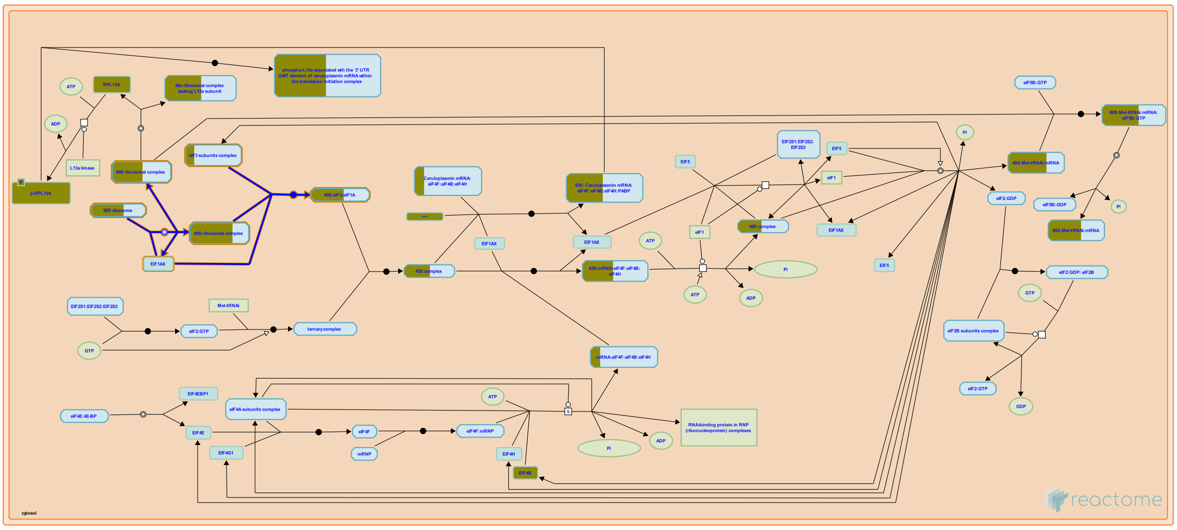

**Cellular compartments:** cytosol.

The 80S ribosome dissociates into free 40S (small) and 60S (large) ribosomal subunits. Each ribosomal subunit is constituted by several individual ribosomal proteins and rRNA.

##### References

##### Edit history

| Date | Action | Author |
|---|---|---|
| 2023-08-31 | Modified | Wright A |

##### 54 submitted entities found in this pathway, mapping to 68 Reactome entities

| Input | UniProt Id | Input | UniProt Id | Input | UniProt Id |
|---|---|---|---|---|---|
| AGRN | P32969 | EIF3F | O00303 | EIF3L | Q9Y262 |
| FAU | P62861 | RPL10 | P27635, Q96L21 | RPL10A | P61313, P62906 |
| RPL13 | P26373, P40429 | RPL13A | P40429, P61313 | RPL15 | P61313 |
| RPL18 | Q07020 | RPL21 | P46778 | RPL27A | P46776, P61353 |
| RPL29 | P47914 | RPL3 | P39023, Q92901 | RPL30 | P62888 |
| RPL31 | P62899 | RPL32 | P62888, P62910 | RPL34 | P49207, P62899 |
| RPL35 | P42766 | RPL35A | P18077 | RPL36 | Q9Y3U8 |
| RPL38 | P63173 | RPL3L | Q92901 | RPL4 | P36578 |
| RPL5 | P46777 | RPL7 | P18124 | RPL7A | P18124, P62424 |
| RPL9P8 | P32969 | RPLP0 | P05388 | RPLP2 | P05387 |
| RPS10 | P46783 | RPS11 | P62280 | RPS12 | P25398 |
| RPS13 | P62277 | RPS14 | P62263 | RPS15 | P62841 |
| RPS15A | P62244 | RPS16 | P62249 | RPS17 | P08708 |
| RPS18 | P62269 | RPS2 | P15880, P46782 | RPS23 | P62266 |
| RPS27 | P42677, Q71UM5 | RPS27A | P42677, P62979, P62987 | RPS29 | P62273 |
| RPS3 | P23396 | RPS3A | P61247 | RPS4X | P62701, Q8TD47 |
| RPS5 | P46782 | RPS6 | P62753 | RPS8 | P62241 |
| RPS9 | P46781 | RPSA | P08865 | SOS2 | P42766 |

#### 5. Nonsense-Mediated Decay (NMD) (R-HSA-927802)

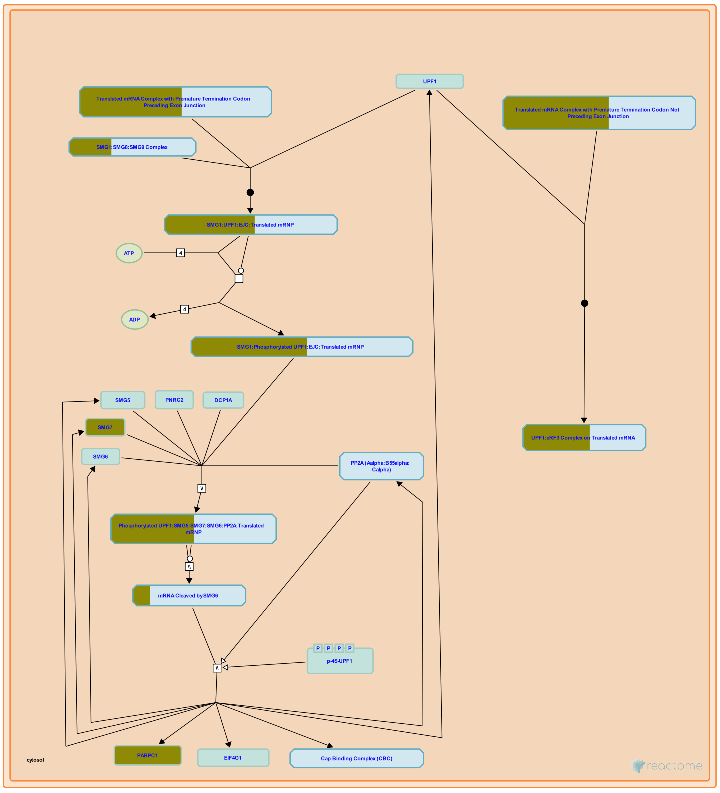


**Cellular compartments:** cytosol.

The Nonsense-Mediated Decay (NMD) pathway activates the destruction of mRNAs containing premature termination codons (PTCs) (reviewed in Isken and Maquat 2007, Chang et al. 2007, Behm-Ansmant et al. 2007, Neu-Yilik and Kulozik 2008, Rebbapragada and Lykke-Andersen 2009, Bhuvanagiri et al. 2010, Nicholson et al. 2010, Durand and Lykke-Andersen 2011). In mammalian cells a termination codon can be recognized as premature if it precedes an exon-exon junction by at least 50-55 nucleotides or if it is followed by an abnormal 3' untranslated region (UTR). While length of the UTR may play a part, the qualifications for being "abnormal" have not been fully elucidated. Also, some termination codons preceding exon junctions are not degraded by NMD so the criteria for triggering NMD are not yet fully known (reviewed in Rebbapragada and Lykke-Andersen 2009). While about 30% of disease-associated mutations in humans activate NMD, about 10% of normal human transcripts are also degraded by NMD (reviewed in Stalder and Muhlemann 2008, Neu-Yilik and Kulozik 2008, Bhuvanagiri et al. 2010, Nicholson et al. 2010). Thus NMD is a normal physiological process controlling mRNA stability in unmutated cells.

Exon junction complexes (EJCs) are deposited on an mRNA during splicing in the nucleus and are displaced by ribosomes during the first round of translation. When a ribosome terminates translation the A site encounters the termination codon and the eRF1 factor enters the empty A site and recruits eRF3. Normally, eRF1 cleaves the translated polypeptide from the tRNA in the P site and eRF3 interacts with Polyadenylate-binding protein (PABP) bound to the polyadenylated tail of the mRNA.

During activation of NMD eRF3 interacts with UPF1 which is contained in a complex with SMG1, SMG8, and SMG9. NMD can arbitrarily be divided into EJC-enhanced and EJC-independent pathways. In EJC-enhanced NMD, an exon junction is located downstream of the PTC and the EJC remains on the mRNA after termination of the pioneer round of translation. The core EJC is associated with UPF2 and UPF3, which interact with UPF1 and stimulate NMD. Once bound near the PTC, UPF1 is phosphorylated by SMG1. The phosphorylation is the rate-limiting step in NMD and causes UPF1 to recruit either SMG6, which is an endoribonuclease, or SMG5 and SMG7, which recruit ribonucleases. SMG6 and SMG5:SMG7 recruit phosphatase PP2A to dephosphorylate UPF1 and allow further rounds of degradation. How EJC-independent NMD is activated remains enigmatic but may involve competition between PABP and UPF1 for eRF3.

#### References


Yepiskoposyan H, Kleinschmidt N, Zamudio Orozco R, Metze S, Muhlemann O & Nicholson P (2010). Nonsense-mediated mRNA decay in human cells: mechanistic insights, functions beyond quality control and the double-life of NMD factors. Cell Mol Life Sci, 67, 677-700.

Saulière J, Wittkopp N, Behm-Ansmant I, Izaurralde E, Rehwinkel J & Kashima I (2007). mRNA quality control: an ancient machinery recognizes and degrades mRNAs with nonsense codons. FEBS Lett, 581, 2845-53.

Durand S & Lykke-Andersen J (2011). SnapShot: Nonsense-Mediated mRNA Decay. Cell, 145, 324-324.e2.

Bhuvanagiri M, Kulozik AE, Hentze MW & Schlitter AM (2010). NMD: RNA biology meets human genetic medicine. Biochem J, 430, 365-77.

Neu-Yilik G & Kulozik AE (2008). NMD: multitasking between mRNA surveillance and modulation of gene expression. Adv Genet, 62, 185-243.


#### Edit history

| Date | Action | Author |
|---|---|---|
| 2010-08-06 | Edited | May B |
| 2010-08-06 | Authored | May B |
| 2010-08-10 | Created | May B |
| 2011-05-19 | Reviewed | Neu-Yilik G |
| 2023-08-26 | Modified | Wright A |

#### 58 submitted entities found in this pathway, mapping to 72 Reactome entities

| Input | UniProt Id | Input | UniProt Id | Input | UniProt Id |
|---|---|---|---|---|---|
| AGRN | P32969 | CASC3 | O15234 | FAU | P62861 |
| MAGOH | P61326 | PABPC1 | P11940 | RPL10 | P27635, Q96L21 |
| RPL10A | P61313, P62906 | RPL13 | P26373, P40429 | RPL13A | P40429, P61313 |
| RPL15 | P61313 | RPL18 | Q07020 | RPL21 | P46778 |
| RPL27A | P46776, P61353 | RPL29 | P47914 | RPL3 | P39023, Q92901 |
| RPL30 | P62888 | RPL31 | P62899 | RPL32 | P62888, P62910 |
| RPL34 | P49207, P62899 | RPL35 | P42766 | RPL35A | P18077 |
| RPL36 | Q9Y3U8 | RPL38 | P63173 | RPL3L | Q92901 |
| RPL4 | P36578 | RPL5 | P46777 | RPL7 | P18124 |

| Input | UniProt Id | Input | UniProt Id | Input | UniProt Id |
|---|---|---|---|---|---|
| RPL7A | P18124, P62424 | RPL9P8 | P32969 | RPLP0 | P05388 |
| RPLP2 | P05387 | RPS10 | P46783 | RPS11 | P62280 |
| RPS12 | P25398 | RPS13 | P62277 | RPS14 | P62263 |
| RPS15 | P62841 | RPS15A | P62244 | RPS16 | P62249 |
| RPS17 | P08708 | RPS18 | P62269 | RPS2 | P15880, P46782 |
| RPS23 | P62266 | RPS27 | P42677, Q71UM5 | RPS27A | P42677, P62979, P62987 |
| RPS29 | P62273 | RPS3 | P23396 | RPS3A | P61247 |
| RPS4X | P62701, Q8TD47 | RPS5 | P46782 | RPS6 | P62753 |
| RPS8 | P62241 | RPS9 | P46781 | RPSA | P08865 |
| SMG7 | Q92540 | SMG9 | Q9H0W8 | SOS2 | P42766 |
| UPF3B | Q9BZI7 | | | | |

#### 6. Nonsense Mediated Decay (NMD) enhanced by the Exon Junction Complex (EJC) (R-HSA-975957)

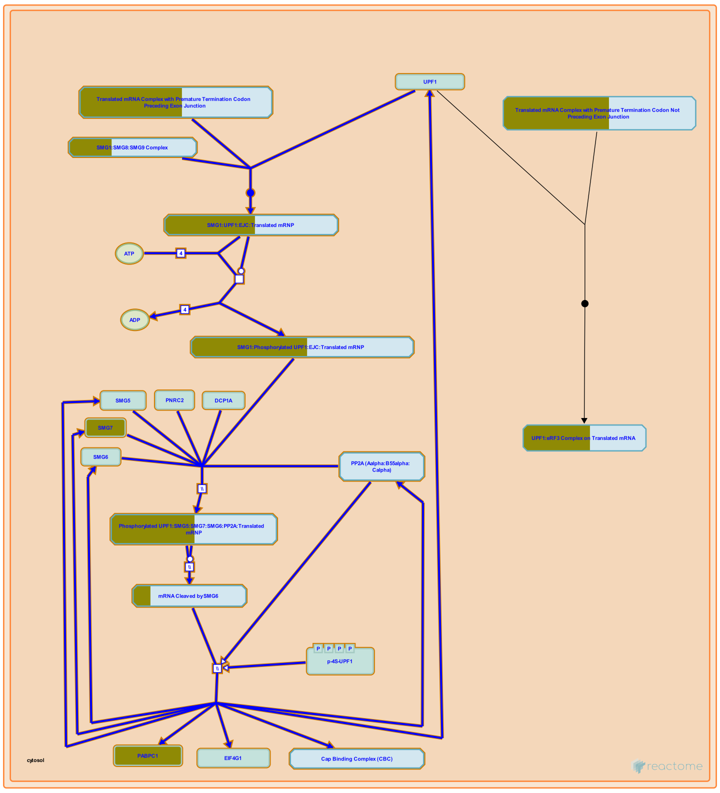


**Cellular compartments:** cytosol.

During normal translation termination eRF3 associates with the ribosome and then interacts with PABP bound to the polyadenylate tail of the mRNA to release the ribosome and allow a new round of translation to commence. Nonsense-mediated decay (NMD) is triggered if eRF3 at the ribosome interacts with UPF1, which may compete with PABP (reviewed in Isken and Maquat 2007, Chang et al. 2007, Behm-Ansmant et al. 2007, Rebbapragada and Lykke-Andersen 2009, Bhuvanagiri et al. 2010, Nicholson et al. 2010, Durand and Lykke-Andersen 2011). An exon junction located 50-55 nt downstream of a termination codon is observed to enhance NMD.

Exon-junction complexes (EJCs) are deposited on the mRNA during splicing in the nucleus, remain on mRNAs after transport to the cytosol, and are dislodged by the ribosome as it progresses along the mRNA during the pioneer round of translation (Gehring et al. 2009). EJCs contain the core factors eIF4A-III, Magoh-Y14, and CASC3 as well as the peripheral factors RNPS1, UPF2, and UPF3. UPF2 and UPF3  recruit UPF1 to eRF3 at the terminating ribosome. Thus an EJC downstream of a termination codon will not have been dislodged during translation and will recruit UPF1, triggering NMD.

UPF1 is believed to form a complex containing SMG1, SMG8, and SMG9. In the key regulatory step of NMD SMG1 phosphorylates UPF1. The phosphorylated UPF1 then recruits either SMG6 or SMG5 and SMG7. SMG6 is itself an endoribonuclease that cleaves the mRNA. SMG5 and SMG7 do not have endoribonuclease activty, but are thought to recruit ribonucleases. Nonsense-mediated decay has been observed to involve deadenlyation, decapping, and both 5' to 3' and 3' to 5' exonuclease activities, but the exact degradative pathways taken by a given mRNA are not yet known.

UPF1 also plays roles in Staufen-mediated decay, histone mRNA decay, telomere maintenance, genome integrity, and may play a role in normal termination of translation.

#### References


Yepiskoposyan H, Kleinschmidt N, Zamudio Orozco R, Metze S, Muhlemann O & Nicholson P (2010). Nonsense-mediated mRNA decay in human cells: mechanistic insights, functions beyond quality control and the double-life of NMD factors. Cell Mol Life Sci, 67, 677-700.

Saulière J, Wittkopp N, Behm-Ansmant I, Izaurralde E, Rehwinkel J & Kashima I (2007). mRNA quality control: an ancient machinery recognizes and degrades mRNAs with nonsense codons. FEBS Lett, 581, 2845-53.

Durand S & Lykke-Andersen J (2011). SnapShot: Nonsense-Mediated mRNA Decay. Cell, 145, 324-324.e2.

Gehring NH, Lamprinaki S, Kulozik AE & Hentze MW (2009). Disassembly of exon junction complexes by PYM. Cell, 137, 536-48.

Bhuvanagiri M, Kulozik AE, Hentze MW & Schlitter AM (2010). NMD: RNA biology meets human genetic medicine. Biochem J, 430, 365-77.


#### Edit history

| Date | Action | Author |
|---|---|---|
| 2010-10-08 | Edited | May B |
| 2010-10-08 | Authored | May B |
| 2010-10-11 | Created | May B |
| 2011-05-19 | Reviewed | Neu-Yilik G |
| 2023-08-31 | Modified | Wright A |

#### 58 submitted entities found in this pathway, mapping to 72 Reactome entities

| Input | UniProt Id | Input | UniProt Id | Input | UniProt Id |
|---|---|---|---|---|---|
| AGRN | P32969 | CASC3 | O15234 | FAU | P62861 |
| MAGOH | P61326 | PABPC1 | P11940 | RPL10 | P27635, Q96L21 |
| RPL10A | P61313, P62906 | RPL13 | P26373, P40429 | RPL13A | P40429, P61313 |
| RPL15 | P61313 | RPL18 | Q07020 | RPL21 | P46778 |
| RPL27A | P46776, P61353 | RPL29 | P47914 | RPL3 | P39023, Q92901 |
| RPL30 | P62888 | RPL31 | P62899 | RPL32 | P62888, P62910 |
| RPL34 | P49207, P62899 | RPL35 | P42766 | RPL35A | P18077 |
| RPL36 | Q9Y3U8 | RPL38 | P63173 | RPL3L | Q92901 |
| RPL4 | P36578 | RPL5 | P46777 | RPL7 | P18124 |
| RPL7A | P18124, P62424 | RPL9P8 | P32969 | RPLP0 | P05388 |
| RPLP2 | P05387 | RPS10 | P46783 | RPS11 | P62280 |
| RPS12 | P25398 | RPS13 | P62277 | RPS14 | P62263 |
| RPS15 | P62841 | RPS15A | P62244 | RPS16 | P62249 |
| RPS17 | P08708 | RPS18 | P62269 | RPS2 | P15880, P46782 |
| RPS23 | P62266 | RPS27 | P42677, Q71UM5 | RPS27A | P42677, P62979, P62987 |
| RPS29 | P62273 | RPS3 | P23396 | RPS3A | P61247 |
| RPS4X | P62701, Q8TD47 | RPS5 | P46782 | RPS6 | P62753 |

| Input | UniProt Id | Input | UniProt Id | Input | UniProt Id |
|---|---|---|---|---|---|
| RPS8 | P62241 | RPS9 | P46781 | RPSA | P08865 |
| SMG7 | Q92540 | SMG9 | Q9H0W8 | SOS2 | P42766 |
| UPF3B | Q9BZI7 | | | | |

#### 7. L13a-mediated translational silencing of Ceruloplasmin expression (R-HSA-156827)

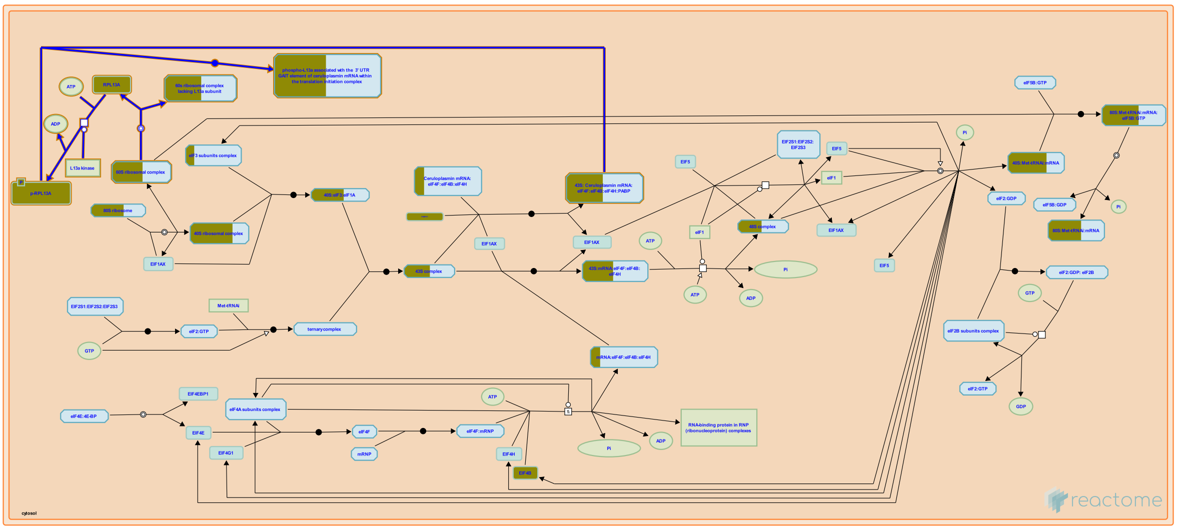

**Cellular compartments:** cytosol.

While circularization of mRNA during translation initiation is thought to contribute to an increase in the efficiency of translation, it also appears to provide a mechanism for translational silencing. This might be achieved by bringing inhibitory 3' UTR-binding proteins into a position in which they interfere either with the function of the translation initiation complex or with the assembly of the ribosome (Mazumder et al 2001). Translational silencing of Ceruloplasmin (Cp) occurs 16 hrs after its induction by INF-gamma (Mazumder et al., 1997). Although the mechanism by which silencing occurs has not yet been determined, this process is mediated by the L13a subunit of the 60s ribosome and thought to require circularization of the Cp mRNA (Sampath et al., 2003; Mazumder et al., 2001; Mazumder et al., 2003). Between 14 and 16 hrs after INF gamma induction, the L13a subunit of the 60s ribosome is phosphorylated and released from the 60s subunit. Phosphorylated L13a then associates with the GAIT element in the 3' UTR of the Cp mRNA inhibiting its translation.

##### References


Seshadri V, Sampath P, DiCorleto PE, Maitra RK, Fox PL & Mazumder B (2003). Regulated release of L13a from the 60S ribosomal subunit as a mechanism of transcript-specific translational control . Cell, 115, 187-98.


##### Edit history

| Date | Action | Author |
|---|---|---|
| 2004-12-13 | Authored | Matthews L |
| 2004-12-20 | Created | Gebauer F |
| 2013-11-25 | Edited | Matthews L |
| 2023-08-31 | Modified | Wright A |

##### 56 submitted entities found in this pathway, mapping to 70 Reactome entities

| Input | UniProt Id | Input | UniProt Id | Input | UniProt Id |
|---|---|---|---|---|---|
| AGRN | P32969 | EIF3F | O00303 | EIF3L | Q9Y262 |

| Input | UniProt Id | Input | UniProt Id | Input | UniProt Id |
|---|---|---|---|---|---|
| EIF4B | P23588 | FAU | P62861 | PABPC1 | P11940 |
| RPL10 | P27635, Q96L21 | RPL10A | P61313, P62906 | RPL13 | P26373, P40429 |
| RPL13A | P40429, P61313 | RPL15 | P61313 | RPL18 | Q07020 |
| RPL21 | P46778 | RPL27A | P46776, P61353 | RPL29 | P47914 |
| RPL3 | P39023, Q92901 | RPL30 | P62888 | RPL31 | P62899 |
| RPL32 | P62888, P62910 | RPL34 | P49207, P62899 | RPL35 | P42766 |
| RPL35A | P18077 | RPL36 | Q9Y3U8 | RPL38 | P63173 |
| RPL3L | Q92901 | RPL4 | P36578 | RPL5 | P46777 |
| RPL7 | P18124 | RPL7A | P18124, P62424 | RPL9P8 | P32969 |
| RPLP0 | P05388 | RPLP2 | P05387 | RPS10 | P46783 |
| RPS11 | P62280 | RPS12 | P25398 | RPS13 | P62277 |
| RPS14 | P62263 | RPS15 | P62841 | RPS15A | P62244 |
| RPS16 | P62249 | RPS17 | P08708 | RPS18 | P62269 |
| RPS2 | P15880, P46782 | RPS23 | P62266 | RPS27 | P42677, Q71UM5 |
| RPS27A | P42677, P62979, P62987 | RPS29 | P62273 | RPS3 | P23396 |
| RPS3A | P61247 | RPS4X | P62701, Q8TD47 | RPS5 | P46782 |
| RPS6 | P62753 | RPS8 | P62241 | RPS9 | P46781 |
| RPSA | P08865 | SOS2 | P42766 | | |

#### 8. GTP hydrolysis and joining of the 60S ribosomal subunit (R-HSA-72706)

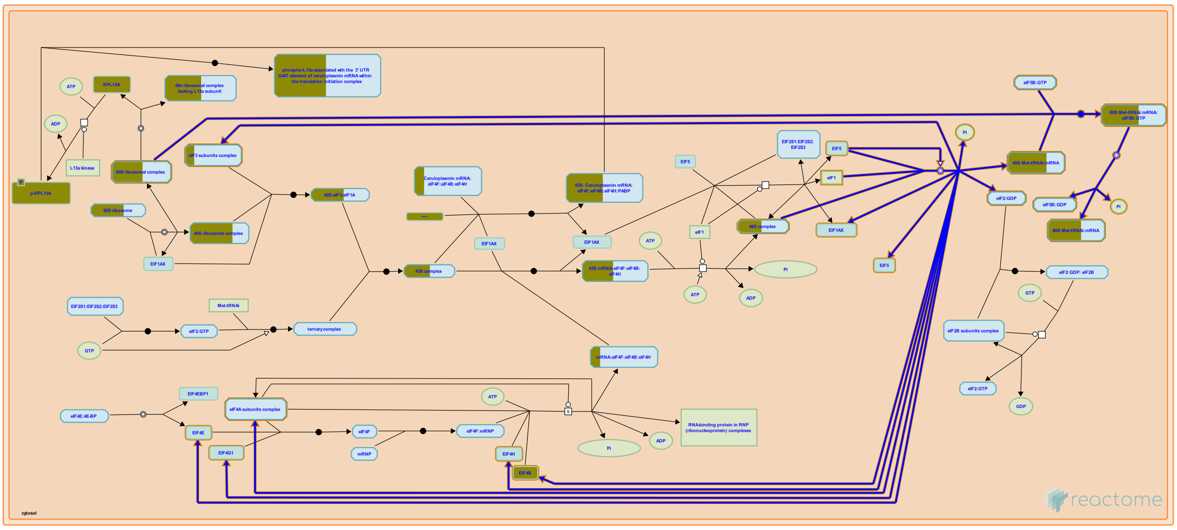

Hydrolysis of eIF2-GTP occurs after the Met-tRNAi has recognized the AUG. This reaction is catalyzed by eIF5 (or eIF5B) and is thought to cause dissociation of all other initiation factors and allow joining of the large 60S ribosomal subunit. The 60S subunit joins - a reaction catalyzed by eIF5 or eIF5B - resulting in a translation-competent 80S ribosome. Following 60S subunit joining, eIF5B hydrolyzes its GTP and is released from the 80S ribosome, which is now ready to start elongating the polypeptide chain.

##### References


Anderson WF, Merrick WC & Kemper WM (1975). Purification and characterization of homogeneous initiation factor M2A from rabbit reticulocytes. J Biol Chem, 250, 5556-62.

Erni B, Schreier MH, Trachsel H & Staehelin T (1978). Initiation of mammalian protein synthesis. II. The assembly of the initiation complex with purified initiation factors. J Mol Biol, 116, 755-67.

Benne R & Hershey JW (1978). The mechanism of action of protein synthesis initiation factors from rabbit reticulocytes. J Biol Chem, 253, 3078-87.

Choi SK, Dever TE, Lomakin IB, Lee JH, Hellen CU & Pestova TV (2000). The joining of ribosomal subunits in eukaryotes requires eIF5B. Nature, 403, 332-5.

Maitra U & Chakrabarti A (1991). Function of eukaryotic initiation factor 5 in the formation of an 80 S ribosomal polypeptide chain initiation complex. J Biol Chem, 266, 14039-45.


##### Edit history

| Date | Action | Author |
|---|---|---|
| 2002-12-16 | Created | Merrick WC |
| 2023-08-31 | Modified | Wright A |

##### 55 submitted entities found in this pathway, mapping to 69 Reactome entities

| Input | UniProt Id | Input | UniProt Id | Input | UniProt Id |
|---|---|---|---|---|---|
| AGRN | P32969 | EIF3F | O00303 | EIF3L | Q9Y262 |
| EIF4B | P23588 | FAU | P62861 | RPL10 | P27635, Q96L21 |

| Input | UniProt Id | Input | UniProt Id | Input | UniProt Id |
|---|---|---|---|---|---|
| RPL10A | P61313, P62906 | RPL13 | P26373, P40429 | RPL13A | P40429, P61313 |
| RPL15 | P61313 | RPL18 | Q07020 | RPL21 | P46778 |
| RPL27A | P46776, P61353 | RPL29 | P47914 | RPL3 | P39023, Q92901 |
| RPL30 | P62888 | RPL31 | P62899 | RPL32 | P62888, P62910 |
| RPL34 | P49207, P62899 | RPL35 | P42766 | RPL35A | P18077 |
| RPL36 | Q9Y3U8 | RPL38 | P63173 | RPL3L | Q92901 |
| RPL4 | P36578 | RPL5 | P46777 | RPL7 | P18124 |
| RPL7A | P18124, P62424 | RPL9P8 | P32969 | RPLP0 | P05388 |
| RPLP2 | P05387 | RPS10 | P46783 | RPS11 | P62280 |
| RPS12 | P25398 | RPS13 | P62277 | RPS14 | P62263 |
| RPS15 | P62841 | RPS15A | P62244 | RPS16 | P62249 |
| RPS17 | P08708 | RPS18 | P62269 | RPS2 | P15880, P46782 |
| RPS23 | P62266 | RPS27 | P42677, Q71UM5 | RPS27A | P42677, P62979, P62987 |
| RPS29 | P62273 | RPS3 | P23396 | RPS3A | P61247 |
| RPS4X | P62701, Q8TD47 | RPS5 | P46782 | RPS6 | P62753 |
| RPS8 | P62241 | RPS9 | P46781 | RPSA | P08865 |
| SOS2 | P42766 | | | | |

#### 9. SRP-dependent cotranslational protein targeting to membrane (R-HSA-1799339)

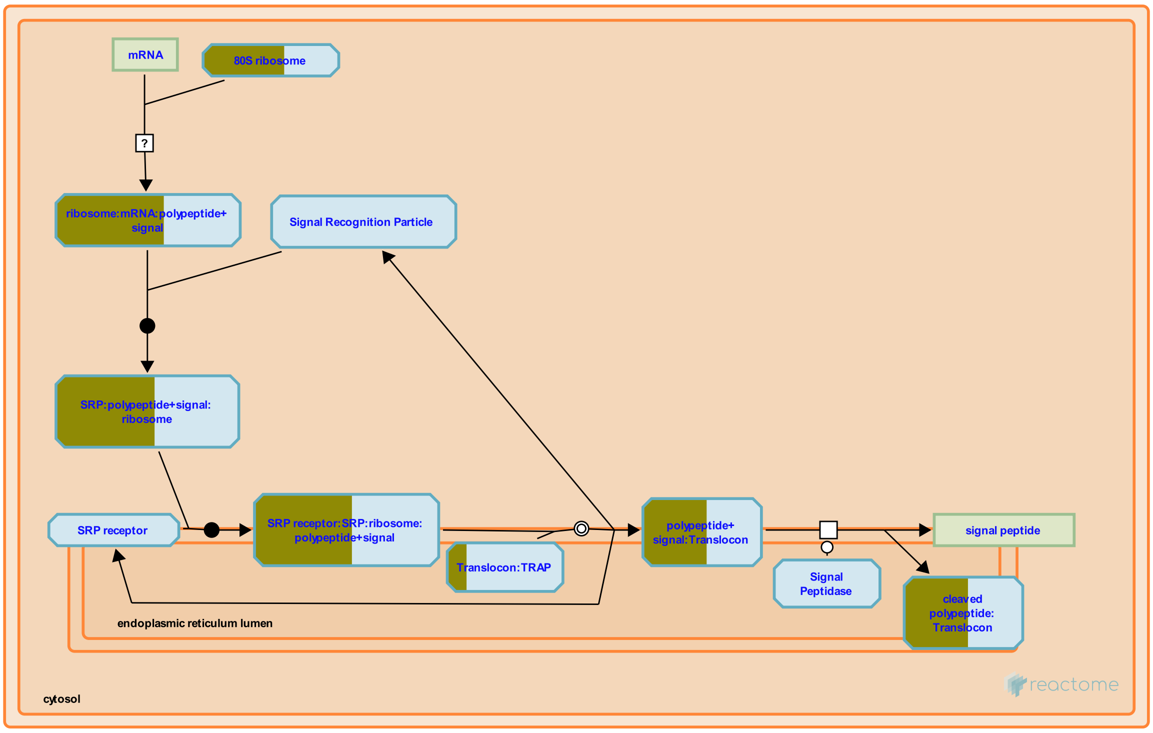


**Cellular compartments:** endoplasmic reticulum membrane, endoplasmic reticulum lumen, cytosol.

The process for translation of a protein destined for the endoplasmic reticulum (ER) branches from the canonical cytoslic translation process at the point when a nascent polypeptide containing a hydrophobic signal sequence is exposed on the surface of the cytosolic ribosome:mRNA:peptide complex. The signal sequence mediates the interaction of this complex with a cytosolic signal recognition particle (SRP) to form a complex which in turn docks with an SRP receptor complex on the ER membrane. There the ribosome complex is transferred from the SRP complex to a translocon complex embedded in the ER membrane and reoriented so that the nascent polypeptide protrudes through a pore in the translocon into the ER lumen. Translation, which had been halted by SRP binding, now resumes, the signal peptide is cleaved from the polypeptide, and elongation proceeds, with the growing polypeptide oriented into the ER lumen.

##### References

##### Edit history

| Date | Action | Author |
|---|---|---|
| 2008-11-20 | Authored | May B, Gopinathrao G |
| 2008-12-02 | Reviewed | Matthews L, Gillespie ME, D'Eustachio P |
| 2011-10-22 | Revised | D'Eustachio P |
| 2011-10-23 | Edited | D'Eustachio P |
| 2011-10-23 | Created | D'Eustachio P |
| 2023-08-31 | Modified | Wright A |

#### 54 submitted entities found in this pathway, mapping to 68 Reactome entities

| Input | UniProt Id | Input | UniProt Id | Input | UniProt Id |
|---|---|---|---|---|---|
| AGRN | P32969 | FAU | P62861 | RPL10 | P27635, Q96L21 |
| RPL10A | P61313, P62906 | RPL13 | P26373, P40429 | RPL13A | P40429, P61313 |
| RPL15 | P61313 | RPL18 | Q07020 | RPL21 | P46778 |
| RPL27A | P46776, P61353 | RPL29 | P47914 | RPL3 | P39023, Q92901 |
| RPL30 | P62888 | RPL31 | P62899 | RPL32 | P62888, P62910 |
| RPL34 | P49207, P62899 | RPL35 | P42766 | RPL35A | P18077 |
| RPL36 | Q9Y3U8 | RPL38 | P63173 | RPL3L | Q92901 |
| RPL4 | P36578 | RPL5 | P46777 | RPL7 | P18124 |
| RPL7A | P18124, P62424 | RPL9P8 | P32969 | RPLP0 | P05388 |
| RPLP2 | P05387 | RPN2 | P04844 | RPS10 | P46783 |
| RPS11 | P62280 | RPS12 | P25398 | RPS13 | P62277 |
| RPS14 | P62263 | RPS15 | P62841 | RPS15A | P62244 |
| RPS16 | P62249 | RPS17 | P08708 | RPS18 | P62269 |
| RPS2 | P15880, P46782 | RPS23 | P62266 | RPS27 | P42677, Q71UM5 |
| RPS27A | P42677, P62979, P62987 | RPS29 | P62273 | RPS3 | P23396 |
| RPS3A | P61247 | RPS4X | P62701, Q8TD47 | RPS5 | P46782 |
| RPS6 | P62753 | RPS8 | P62241 | RPS9 | P46781 |
| RPSA | P08865 | SEC61G | P60059 | SOS2 | P42766 |

#### 10. Interferon gamma signaling (R-HSA-877300)

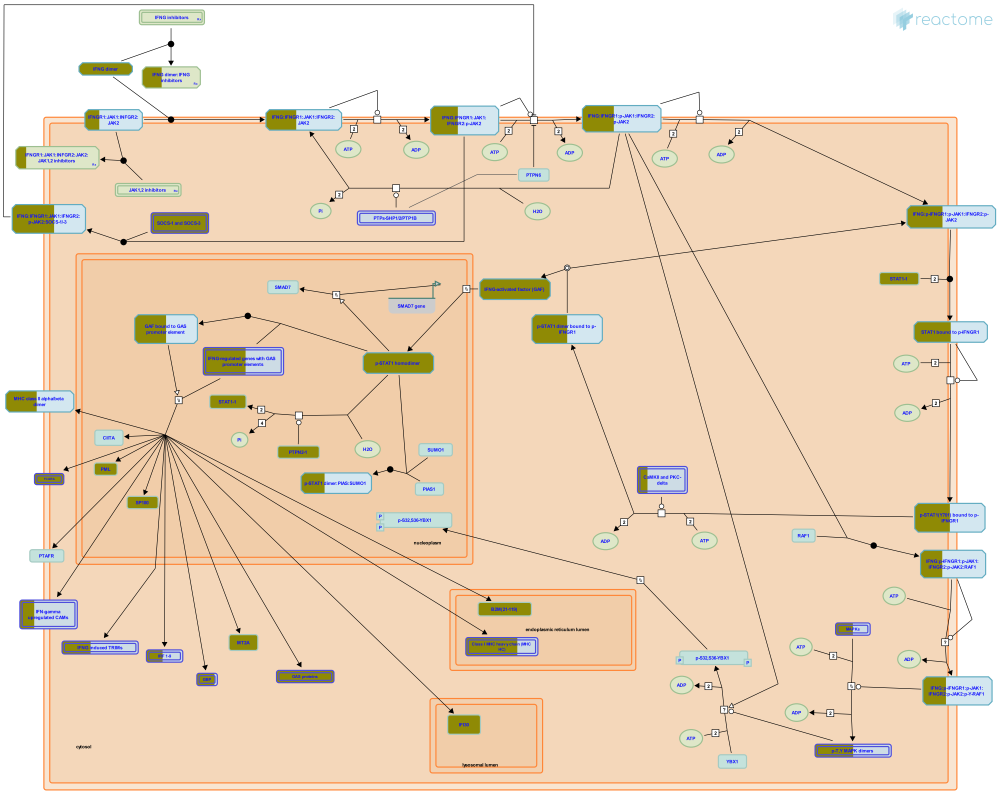

Interferon-gamma (IFN-gamma) belongs to the type II interferon family and is secreted by activated immune cells-primarily T and NK cells, but also B-cells and APC. INFG exerts its effect on cells by interacting with the specific IFN-gamma receptor (IFNGR). IFNGR consists of two chains, namely IFNGR1 (also known as the IFNGR alpha chain) and IFNGR2 (also known as the IFNGR beta chain). IFNGR1 is the ligand binding receptor and is required but not sufficient for signal transduction, whereas IFNGR2 do not bind IFNG independently but mainly plays a role in IFNG signaling and is generally the limiting factor in IFNG responsiveness. Both IFNGR chains lack intrinsic kinase/phosphatase activity and thus rely on other signaling proteins like Janus-activated kinase 1 (JAK1), JAK2 and Signal transducer and activator of transcription 1 (STAT-1) for signal transduction. IFNGR complex in its resting state is a preformed tetramer and upon IFNG association undergoes a conformational change. This conformational change induces the phosphorylation and activation of JAK1, JAK2, and STAT1 which in turn induces genes containing the gamma-interferon activation sequence (GAS) in the promoter.

##### References


Schroder K, Ravasi T, Hume DA & Hertzog PJ (2004). Interferon-gamma: an overview of signals, mechanisms and functions. J Leukoc Biol, 75, 163-89.

Aguet M, Bach EA & Schreiber RD (1997). The IFN gamma receptor: a paradigm for cytokine receptor signaling. Annu Rev Immunol, 15, 563-91.

Gough DJ, Levy DE, Clarke CJ & Johnstone RW (2008). IFNgamma signaling-does it mean JAK-STAT?. Cytokine Growth Factor Rev, 19, 383-94.

Izotova LS, Garotta G, Muthukumaran G, Kotenko SV, Cook JR & Pestka S (1997). The interferon gamma (IFN-gamma) receptor: a paradigm for the multichain cytokine receptor. Cytokine Growth Factor Rev, 8, 189-206.

#### Edit history

| Date | Action | Author |
|---|---|---|
| 2010-06-08 | Edited | Garapati P V |
| 2010-06-08 | Authored | Garapati P V |
| 2010-06-11 | Created | Garapati P V |
| 2010-08-17 | Reviewed | Abdul-Sater AA, Schindler C |
| 2023-08-31 | Modified | Wright A |

#### 49 submitted entities found in this pathway, mapping to 89 Reactome entities

| Input | UniProt Id | Input | UniProt Id | Input | UniProt Id |
|---|---|---|---|---|---|
| B2M | P61769 | FCGR1A | P12314 | FCGR1B | Q92637 |
| GBP1 | P32455 | GBP2 | P32456 | GBP3 | Q9H0R5 |
| GBP4 | Q96PP9 | GBP5 | Q96PP8 | GBP6 | Q6ZN66 |
| HLA-A | P04439 | HLA-B | P01889 | HLA-C | P10321 |
| HLA-DQA1 | P01909, P20036 | HLA-DQA2 | P01906 | HLA-DRA | P01903 |
| HLA-DRB4 | P13762 | HLA-G | P17693 | HLA-H | P01893 |
| IFI30 | P13284 | IFNG | P01579 | IRF1 | P10914 |
| IRF5 | Q13568 | IRF7 | Q92985 | IRF9 | Q00978 |
| JAK2 | O60674 | MAPK3 | P27361 | MT1A | P02795 |
| MT2A | P02795 | NCAM1 | P13591 | OAS1 | P00973 |
| OAS2 | P29728 | OAS3 | Q9Y6K5 | OASL | Q15646 |
| PML | P29590 | PRKCD | Q05655 | PTPN2 | P17706-2 |
| SOCS1 | O15524 | SOCS3 | O14543 | SP100 | P23497 |
| SP110 | P23497 | STAT1 | P42224-1 | TRIM14 | Q14142 |
| TRIM21 | P19474 | TRIM22 | Q8IYM9 | TRIM25 | Q14258 |
| TRIM34 | Q9BYJ4 | TRIM38 | O00635 | TRIM5 | Q9C035 |
| TRIM6 | Q9C030 | | | | |

| Input | Ensembl Id | Input | Ensembl Id | Input | Ensembl Id |
|---|---|---|---|---|---|
| B2M | ENSG00000166710 | FCGR1A | ENSG00000150337 | FCGR1B | ENSG00000198019 |
| GBP1 | ENSG00000117228 | GBP2 | ENSG00000162645 | GBP3 | ENSG00000117226 |
| GBP4 | ENSG00000162654 | GBP5 | ENSG00000154451 | GBP6 | ENSG00000183347 |
| HLA-A | ENSG00000206503 | HLA-B | ENSG00000234745 | HLA-C | ENSG00000204525 |
| HLA-DQA1 | ENSG00000196735 | HLA-DQA2 | ENSG00000237541 | HLA-DRA | ENSG00000204287 |
| HLA-DRB4 | ENSG00000227357 | HLA-G | ENSG00000204632 | HLA-H | ENSG00000206341 |
| IFI30 | ENSG00000216490 | IRF1 | ENSG00000125347 | IRF5 | ENSG00000128604 |
| IRF7 | ENSG00000185507 | IRF9 | ENSG00000213928 | MT2A | ENSG00000125148 |
| NCAM1 | ENSG00000149294 | OAS1 | ENSG00000089127 | OAS2 | ENSG00000111335 |
| OAS3 | ENSG00000111331 | OASL | ENSG00000135114 | PML | ENSG00000140464 |
| SP100 | ENSG00000067066 | TRIM14 | ENSG00000106785 | TRIM21 | ENSG00000132109 |
| TRIM22 | ENSG00000132274 | TRIM25 | ENSG00000121060 | TRIM34 | ENSG00000258659 |
| TRIM38 | ENSG00000112343 | TRIM5 | ENSG00000132256 | TRIM6 | ENSG00000121236 |

### 11. Interferon alpha/beta signaling (R-HSA-909733)

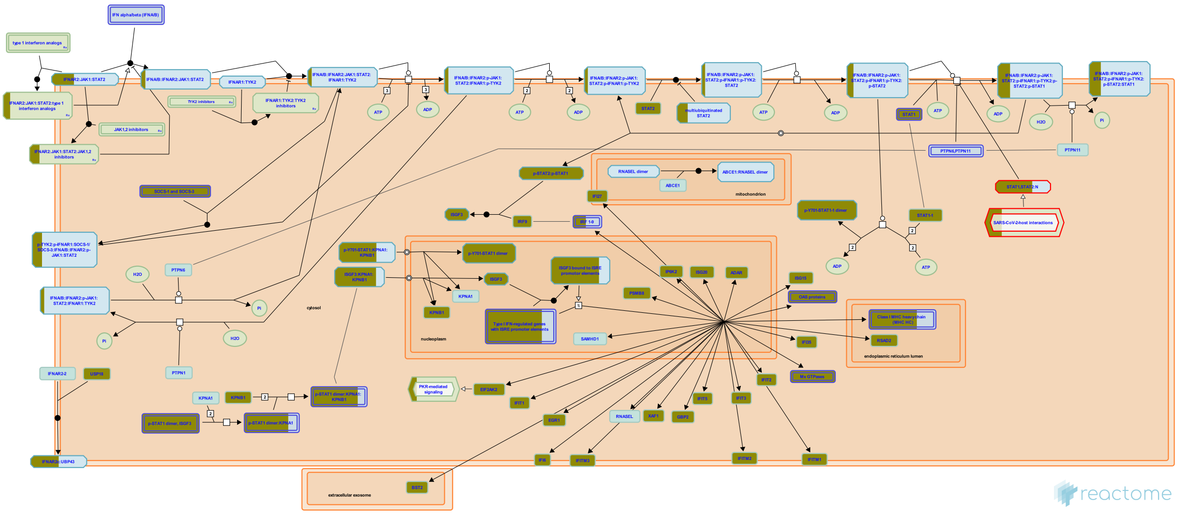

Type I interferons (IFNs) are composed of various genes including IFN alpha (IFNA), beta (IFNB), omega, epsilon, and kappa. In humans the IFNA genes are composed of more than 13 subfamily genes, whereas there is only one IFNB gene. The large family of IFNA/B proteins all bind to a single receptor which is composed of two distinct chains: IFNAR1 and IFNAR2. The IFNA/B stimulation of the IFNA receptor complex leads to the formation of two transcriptional activator complexes: IFNA-activated-factor (AAF), which is a homodimer of STAT1 and IFN-stimulated gene factor 3 (ISGF3), which comprises STAT1, STAT2 and a member of the IRF family, IRF9/P48. AAF mediates activation of the IRF-1 gene by binding to GAS (IFNG-activated site), whereas ISGF3 activates several IFN-inducible genes including IRF3 and IRF7.

#### References


Gupta S, Greenlund AC, Krolewski JJ, Yan H, Schreiber RD, Schindler CW, ... Krishnan K (1996). Phosphorylated interferon-alpha receptor 1 subunit (IFNaR1) acts as a docking site for the latent form of the 113 kDa STAT2 protein. EMBO J, 15, 1064-74.

Pellegrini S, Piehler J, Schreiber G & Uzé G (2007). The receptor of the type I interferon family. Curr Top Microbiol Immunol, 316, 71-95.

Gauzzi MC, Pellegrini S, Velazquez L, McKendry R, Fellous M & Mogensen KE (1996). Interferon-alpha-dependent activation of Tyk2 requires phosphorylation of positive regulatory tyrosines by another kinase. J Biol Chem, 271, 20494-500.

Stark GR, Darnell JE Jr, Qureshi S, Li X & Leung S (1996). Formation of STAT1-STAT2 heterodimers and their role in the activation of IRF-1 gene transcription by interferon-alpha. J Biol Chem, 271, 5790-4.


#### Edit history

| Date | Action | Author |
|---|---|---|
| 2010-07-07 | Edited | Garapati P V |
| 2010-07-07 | Authored | Garapati P V |
| 2010-07-07 | Created | Garapati P V |
| 2010-08-17 | Reviewed | Abdul-Sater AA, Schindler C |
| 2023-08-31 | Modified | Wright A |

#### 43 submitted entities found in this pathway, mapping to 83 Reactome entities

| Input | UniProt Id | Input | UniProt Id | Input | UniProt Id |
|---|---|---|---|---|---|
| ADAR | P55265 | BST2 | Q10589 | EGR1 | P18146 |
| EIF2AK2 | P19525 | GBP2 | P32456 | HLA-A | P04439 |
| HLA-B | P01889 | HLA-C | P10321 | HLA-G | P17693 |
| HLA-H | P01893 | IFI27 | P40305 | IFI35 | P80217 |
| IFI6 | P09912 | IFIT1 | P09914 | IFIT2 | P09913 |
| IFIT3 | O14879 | IFIT5 | Q13325 | IFITM1 | P13164 |
| IFITM2 | Q01629 | IFITM3 | P13164, Q01628 | IP6K2 | Q9UHH9 |
| IRF1 | P10914 | IRF5 | Q13568 | IRF7 | Q92985 |
| IRF9 | Q00978 | ISG15 | P05161 | ISG20 | Q96AZ6 |
| KPNB1 | Q14974 | MX1 | P20591 | MX2 | P20592 |
| OAS1 | P00973 | OAS2 | P29728 | OAS3 | Q9Y6K5 |
| OASL | Q15646 | PSMB8 | P28062 | RPS27A | P62979, P62987 |
| RSAD2 | Q8WXG1 | SOCS1 | O15524 | SOCS3 | O14543 |
| STAT1 | P42224, P42224-1, P42224-2 | STAT2 | P52630 | USP18 | Q9UMW8 |
| XAF1 | Q6GPH4 | | | | |

| Input | Ensembl Id | Input | Ensembl Id | Input | Ensembl Id |
|---|---|---|---|---|---|
| ADAR | ENSG00000160710 | BST2 | ENSG00000130303 | EGR1 | ENSG00000120738 |
| EIF2AK2 | ENSG00000055332 | GBP2 | ENSG00000162645 | HLA-A | ENSG00000206503 |
| HLA-B | ENSG00000234745 | HLA-C | ENSG00000204525 | HLA-G | ENSG00000204632 |
| HLA-H | ENSG00000206341 | IFI27 | ENSG00000165949 | IFI35 | ENSG00000068079 |
| IFI6 | ENSG00000126709 | IFIT1 | ENSG00000185745 | IFIT2 | ENSG00000119922 |
| IFIT3 | ENSG00000119917 | IFIT5 | ENSG00000152778 | IFITM1 | ENSG00000185885 |
| IFITM2 | ENSG00000185201 | IFITM3 | ENSG00000142089 | IP6K2 | ENSG00000068745 |
| IRF1 | ENSG00000125347 | IRF5 | ENSG00000128604 | IRF7 | ENSG00000185507 |
| IRF9 | ENSG00000213928 | ISG15 | ENSG00000187608 | ISG20 | ENSG00000172183 |
| MX1 | ENSG00000157601 | MX2 | ENSG00000183486 | OAS1 | ENSG00000089127 |
| OAS2 | ENSG00000111335 | OAS3 | ENSG00000111331 | OASL | ENSG00000135114 |
| PSMB8 | ENSG00000204264 | RSAD2 | ENSG00000134321 | XAF1 | ENSG00000132530 |

#### 12. Interferon Signaling (R-HSA-913531)

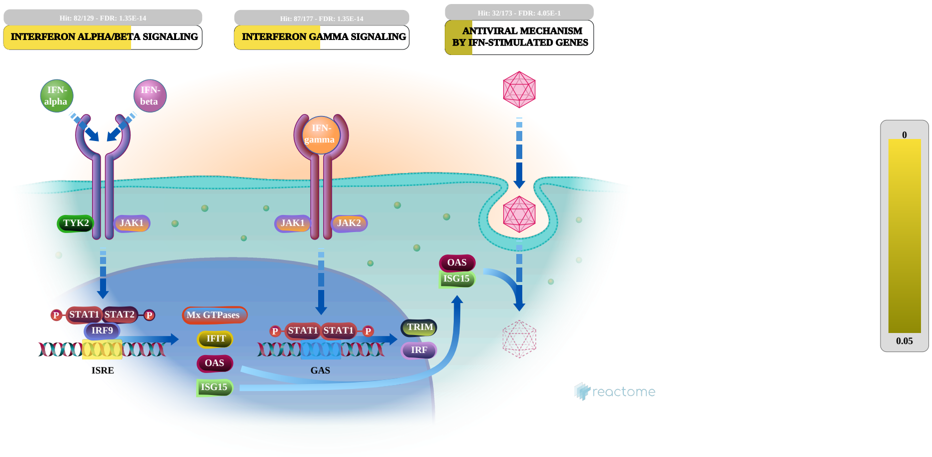


Interferons (IFNs) are cytokines that play a central role in initiating immune responses, especially antiviral and antitumor effects. There are three types of IFNs:Type I (IFN-alpha, -beta and others, such as omega, epsilon, and kappa), Type II (IFN-gamma) and Type III (IFN-lamda). In this module we are mainly focusing on type I IFNs alpha and beta and type II IFN-gamma. Both type I and type II IFNs exert their actions through cognate receptor complexes, IFNAR and IFNGR respectively, present on cell surface membranes. Type I IFNs are broadly expressed heterodimeric receptors composed of the IFNAR1 and IFNAR2 subunits, while the type II IFN receptor consists of IFNGR1 and IFNGR2. Type III interferon lambda has three members: lamda1 (IL-29), lambda2 (IL-28A), and lambda3 (IL-28B) respectively. IFN-lambda signaling is initiated through unique heterodimeric receptor composed of IFN-LR1/IF-28Ralpha and IL10R2 chains.

Type I IFNs typically recruit JAK1 and TYK2 proteins to transduce their signals to STAT1 and 2; in combination with IRF9 (IFN-regulatory factor 9), these proteins form the heterotrimeric complex ISGF3. In nucleus ISGF3 binds to IFN-stimulated response elements (ISRE) to promote gene induction.

Type II IFNs in turn rely upon the activation of JAKs 1 and 2 and STAT1. Once activated, STAT1 dimerizes to form the transcriptional regulator GAF (IFNG activated factor) and this binds to the IFNG activated sequence (GAS) elements and initiate the transcription of IFNG-responsive genes.

Like type I IFNs, IFN-lambda recruits TYK2 and JAK1 kinases and then promote the phosphorylation of STAT1/2, and induce the ISRE3 complex formation.

##### References

Schroder K, Ravasi T, Hume DA & Hertzog PJ (2004). Interferon-gamma: an overview of signals, mechanisms and functions. J Leukoc Biol, 75, 163-89.

Platanias LC (2005). Mechanisms of type-I- and type-II-interferon-mediated signalling. Nat Rev Immunol, 5, 375-86.

Gough DJ, Levy DE, Clarke CJ & Johnstone RW (2008). IFNgamma signaling-does it mean JAK-STAT?. Cytokine Growth Factor Rev, 19, 383-94.

Ferreira PC, Bonjardim CA & Kroon EG (2009). Interferons: signaling, antiviral and viral evasion. Immunol Lett, 122, 1-11.

Platanias LC & Uddin S (2004). Mechanisms of type-I interferon signal transduction. J Biochem Mol Biol, 37, 635-41.

#### Edit history

| Date | Action | Author |
|---|---|---|
| 2010-07-07 | Edited | Garapati P V |
| 2010-07-07 | Authored | Garapati P V |
| 2010-07-16 | Created | Garapati P V |
| 2010-08-17 | Reviewed | Abdul-Sater AA, Schindler C |
| 2023-08-26 | Modified | Wright A |

#### 90 submitted entities found in this pathway, mapping to 158 Reactome entities

| Input | UniProt Id | Input | UniProt Id | Input | UniProt Id |
|---|---|---|---|---|---|
| ADAR | P55265 | B2M | P61769 | BST2 | Q10589 |
| DDX58 | O95786 | EGR1 | P18146 | EIF2AK2 | P19525 |
| FAAP20 | Q6NZ36 | FANCA | O15360 | FANCL | Q9NW38 |
| FCGR1A | P12314 | FCGR1B | Q92637 | GBP1 | P32455 |
| GBP2 | P32456 | GBP3 | Q9H0R5 | GBP4 | Q96PP9 |
| GBP5 | Q96PP8 | GBP6 | Q6ZN66 | HERC5 | Q9UII4 |
| HLA-A | P04439 | HLA-B | P01889 | HLA-C | P10321 |
| HLA-DQA1 | P01909, P20036 | HLA-DQA2 | P01906 | HLA-DRA | P01903 |
| HLA-DRB4 | P13762 | HLA-G | P17693 | HLA-H | P01893 |
| HSPA1A | P0DMV8 | IFI27 | P40305 | IFI30 | P13284 |
| IFI35 | P80217 | IFI6 | P09912 | IFIT1 | P09914 |
| IFIT2 | P09913 | IFIT3 | O14879 | IFIT5 | Q13325 |
| IFITM1 | P13164 | IFITM2 | Q01629 | IFITM3 | P13164, Q01628 |
| IFNG | P01579 | IP6K2 | Q9UHH9 | IRF1 | P10914 |
| IRF5 | Q13568 | IRF7 | Q92985 | IRF9 | Q00978 |
| ISG15 | P05161 | ISG20 | Q96AZ6 | JAK2 | O60674 |
| KPNB1 | Q14974 | MAP2K6 | P52564 | MAPK3 | P27361 |
| MT1A | P02795 | MT2A | P02795 | MX1 | P20591 |
| MX2 | P20592 | NCAM1 | P13591 | NUP188 | Q5SRE5 |
| OAS1 | P00973 | OAS2 | P29728 | OAS3 | Q9Y6K5 |
| OASL | Q15646 | PIN1 | Q13526 | PML | P29590 |
| PRKCD | Q05655 | PSMB8 | P28062 | PTPN2 | P17706, P17706-2 |
| RIGI | O95786 | RPS27A | P62979, P62987 | RSAD2 | Q8WXG1 |
| SOCS1 | O15524 | SOCS3 | O14543 | SP100 | P23497 |
| SP110 | P23497 | STAT1 | P42224, P42224-1, P42224-2 | STAT2 | P52630 |
| TARBP2 | Q15633 | TRIM14 | Q14142 | TRIM21 | P19474 |
| TRIM22 | Q8IYM9 | TRIM25 | Q14258 | TRIM34 | Q9BYJ4 |
| TRIM38 | O00635 | TRIM5 | Q9C035 | TRIM6 | Q9C030 |
| TUBA4A | P68366 | UBA7 | P41226 | UBE2L6 | O14933 |
| USP18 | Q3LFD5, Q9UMW8 | USP41 | Q3LFD5 | XAF1 | Q6GPH4 |

| Input | Ensembl Id | Input | Ensembl Id | Input | Ensembl Id |
|---|---|---|---|---|---|
| ADAR | ENSG00000160710 | B2M | ENSG00000166710 | BST2 | ENSG00000130303 |
| EGR1 | ENSG00000120738 | EIF2AK2 | ENSG00000055332 | FCGR1A | ENSG00000150337 |
| FCGR1B | ENSG00000198019 | GBP1 | ENSG00000117228 | GBP2 | ENSG00000162645 |
| GBP3 | ENSG00000117226 | GBP4 | ENSG00000162654 | GBP5 | ENSG00000154451 |
| GBP6 | ENSG00000183347 | HLA-A | ENSG00000206503 | HLA-B | ENSG00000234745 |
| HLA-C | ENSG00000204525 | HLA-DQA1 | ENSG00000196735 | HLA-DQA2 | ENSG00000237541 |
| HLA-DRA | ENSG00000204287 | HLA-DRB4 | ENSG00000227357 | HLA-G | ENSG00000204632 |
| HLA-H | ENSG00000206341 | IFI27 | ENSG00000165949 | IFI30 | ENSG00000216490 |
| IFI35 | ENSG00000068079 | IFI6 | ENSG00000126709 | IFIT1 | ENSG00000185745 |
| IFIT2 | ENSG00000119922 | IFIT3 | ENSG00000119917 | IFIT5 | ENSG00000152778 |
| IFITM1 | ENSG00000185885 | IFITM2 | ENSG00000185201 | IFITM3 | ENSG00000142089 |
| IP6K2 | ENSG00000068745 | IRF1 | ENSG00000125347 | IRF5 | ENSG00000128604 |
| IRF7 | ENSG00000185507 | IRF9 | ENSG00000213928 | ISG15 | ENSG00000187608 |
| ISG20 | ENSG00000172183 | MT2A | ENSG00000125148 | MX1 | ENSG00000157601 |
| MX2 | ENSG00000183486 | NCAM1 | ENSG00000149294 | OAS1 | ENSG00000089127 |
| OAS2 | ENSG00000111335 | OAS3 | ENSG00000111331 | OASL | ENSG00000135114 |
| PML | ENSG00000140464 | PSMB8 | ENSG00000204264 | RSAD2 | ENSG00000134321 |
| SP100 | ENSG00000067066 | TRIM14 | ENSG00000106785 | TRIM21 | ENSG00000132109 |
| TRIM22 | ENSG00000132274 | TRIM25 | ENSG00000121060 | TRIM34 | ENSG00000258659 |
| TRIM38 | ENSG00000112343 | TRIM5 | ENSG00000132256 | TRIM6 | ENSG00000121236 |
| XAF1 | ENSG00000132530 | | | | |

#### 13. Cytokine Signaling in Immune system (R-HSA-1280215)

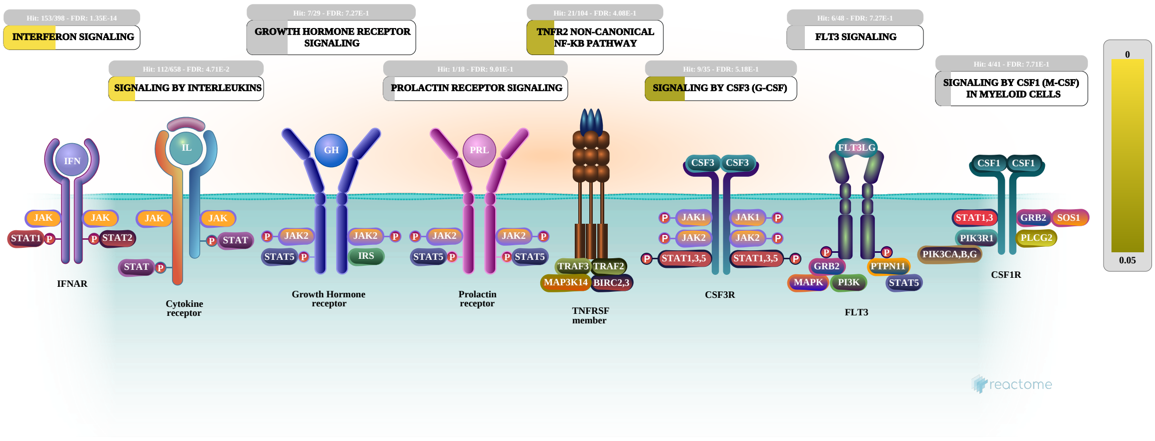


Cytokines are small proteins that regulate and mediate immunity, inflammation, and hematopoiesis. They are secreted in response to immune stimuli, and usually act briefly, locally, at very low concentrations. Cytokines bind to specific membrane receptors, which then signal the cell via second messengers, to regulate cellular activity.

##### References


Feldmann M & Oppenheim J (2002). *Cytokines and the immune system, Cytokine Reference* .

IMMPORT:Bioinformatics for the future of immunology. Retrieved from https://www.immport.org/immportWeb/queryref/geneListSummary.do

Santamaria P (2003). Cytokines and chemokines in autoimmune disease: an overview. Adv Exp Med Biol, 520, 1-7.

COPE. Retrieved from http://www.copewithcytokines.org/cope.cgi


##### Edit history

| Date | Action | Author |
|---|---|---|
| 2011-05-12 | Created | Garapati P V |
| 2011-05-22 | Edited | Ray KP, Jupe S, Garapati P V |
| 2011-05-22 | Authored | Ray KP, Jupe S, Garapati P V |
| 2011-05-29 | Reviewed | Abdul-Sater AA, Schindler C, Pinteaux E |
| 2023-08-26 | Modified | Wright A |

##### 163 submitted entities found in this pathway, mapping to 272 Reactome entities

| Input | UniProt Id | Input | UniProt Id | Input | UniProt Id |
|---|---|---|---|---|---|
| ADAR | P55265 | ALOX15 | P16050 | ARF1 | P84077 |
| B2M | P61769 | BIRC2 | Q13489, Q13490 | BST2 | Q10589 |
| CASP1 | P29466 | CCL2 | P13500 | CCL20 | P78556 |

| Input | UniProt Id | Input | UniProt Id | Input | UniProt Id |
|---|---|---|---|---|---|
| CCR1 | P32246 | CD27 | P26842 | CD40LG | P29965 |
| CDKN1A | P38936 | CISH | Q9NSE2 | CXCL10 | P02778 |
| DDX58 | O95786 | DUSP6 | Q16828 | EDAR | Q9UNE0 |
| EGR1 | P18146 | EIF2AK2 | P19525 | FAAP20 | Q6NZ36 |
| FANCA | O15360 | FANCL | Q9NW38 | FBXW11 | Q9UKB1 |
| FCGR1A | P12314 | FCGR1B | Q92637 | FLT3LG | P49771 |
| GBP1 | P32455 | GBP2 | P32456 | GBP3 | Q9H0R5 |
| GBP4 | Q96PP9 | GBP5 | Q96PP8 | GBP6 | Q6ZN66 |
| H3C4 | P68431, Q71DI3 | HERC5 | Q9UII4 | HLA-A | P04439 |
| HLA-B | P01889 | HLA-C | P10321 | HLA-DQA1 | P01909, P20036 |
| HLA-DQA2 | P01906 | HLA-DRA | P01903 | HLA-DRB4 | P13762 |
| HLA-G | P17693 | HLA-H | P01893 | HNRNPA2B1 | P22626 |
| HSPA1A | P0DMV8 | IFI27 | P40305 | IFI30 | P13284 |
| IFI35 | P80217 | IFI6 | P09912 | IFIT1 | P09914 |
| IFIT2 | P09913 | IFIT3 | O14879 | IFIT5 | Q13325 |
| IFITM1 | P13164 | IFITM2 | Q01629 | IFITM3 | P13164, Q01628 |
| IFNG | P01579 | IGHE | P01854 | IGHG1 | P01857, P01861 |
| IGHG4 | P01861 | IKBIP | Q70UQ0 | IL11RA | P08887, Q14626 |
| IL15 | P40933 | IL1RN | P18510 | IL27 | Q8NEV9 |
| IL2RB | P14784 | IL37 | Q9NZH6 | IL4R | P24394, Q01113 |
| IL5RA | P15509, Q01344 | IL6 | P05231 | IL6R | P08887, P08887-2 |
| IL7R | P16871 | IP6K2 | Q9UHH9 | IRF1 | P10914 |
| IRF5 | Q13568 | IRF7 | Q92985 | IRF9 | Q00978 |
| ISG15 | P05161 | ISG20 | Q96AZ6 | ITGB1 | P05556 |
| JAK2 | O60674 | KPNB1 | Q14974 | LGALS9 | O00182 |
| LIF | P15018 | LMNB1 | P20700 | LTB | Q06643 |
| MAP2K3 | P46734 | MAP2K6 | P52564 | MAPK3 | P27361 |
| MEF2C | Q06413 | MT1A | P02795 | MT2A | P02795 |
| MUC1 | P15941 | MX1 | P20591 | MX2 | P20592 |
| MYD88 | Q99836 | NCAM1 | P13591 | NLRC5 | Q86WI3 |
| NLRX1 | Q86UT6 | NOD2 | Q9HC29 | NUP188 | Q5SRE5 |
| OAS1 | P00973 | OAS2 | P29728 | OAS3 | Q9Y6K5 |
| OASL | Q15646 | PIK3CA | P42336 | PIN1 | Q13526 |
| PML | P29590 | PPIA | P62937 | PPP2R5D | Q14738 |
| PRKCD | Q05655 | PSMB8 | P28062 | PSMB9 | P28065 |
| PSME1 | Q06323 | PSME2 | Q9UL46 | PTK2B | Q14289 |
| PTPN2 | P17706, P17706-2 | REL | Q01201 | RIGI | O95786 |
| RORA | P35398 | RORC | P51449 | RPLP0 | P05388 |
| RPN2 | Q99460 | RPS27A | P62979, P62987 | RSAD2 | Q8WXG1 |
| S100B | P04271 | SIGIRR | Q6IA17 | SOCS1 | O15524 |
| SOCS2 | O14508 | SOCS3 | O14543 | SOS2 | Q07890 |
| SP100 | P23497 | SP110 | P23497 | SQSTM1 | Q13501 |
| STAT1 | P42224, P42224-1, P42224-2 | STAT2 | P52630 | SYK | P43405 |
| TAB3 | Q8N5C8 | TARBP2 | Q15633 | TIFA | Q96CG3 |
| TNFRSF17 | Q02223 | TNFRSF25 | Q93038 | TNFSF13 | O75888 |
| TNFSF13B | Q9Y275 | TNFSF14 | O43557 | TRAF3 | Q13114 |
| TRIM14 | Q14142 | TRIM21 | P19474 | TRIM22 | Q8IYM9 |
| TRIM25 | Q14258 | TRIM34 | Q9BYJ4 | TRIM38 | O00635 |
| TRIM5 | Q9C035 | TRIM6 | Q9C030 | TUBA4A | P68366 |

| Input | UniProt Id | Input | UniProt Id | Input | UniProt Id |
|---|---|---|---|---|---|
| UBA7 | P41226 | UBE2L6 | O14933 | USP18 | Q3LFD5, Q9UMW8 |
| USP41 | Q3LFD5 | VIM | P08670 | VRK3 | Q8IV63 |
| XAF1 | Q6GPH4 | | | | |

| Input | Ensembl Id | Input | Ensembl Id | Input | Ensembl Id |
|---|---|---|---|---|---|
| ADAR | ENSG00000160710 | ALOX15 | ENSG00000161905 | ARF1 | ENSG00000143761 |
| B2M | ENSG00000166710 | BST2 | ENSG00000130303 | CCL2 | ENSG00000108691 |
| CCL20 | ENSG00000115009 | CCR1 | ENSG00000163823 | CDKN1A | ENSG00000124762 |
| CISH | ENSG00000114737 | CXCL10 | ENSG00000169245 | EGR1 | ENSG00000120738 |
| EIF2AK2 | ENSG00000055332 | FCGR1A | ENSG00000150337 | FCGR1B | ENSG00000198019 |
| GBP1 | ENSG00000117228 | GBP2 | ENSG00000162645 | GBP3 | ENSG00000117226 |
| GBP4 | ENSG00000162654 | GBP5 | ENSG00000154451 | GBP6 | ENSG00000183347 |
| HLA-A | ENSG00000206503 | HLA-B | ENSG00000234745 | HLA-C | ENSG00000204525 |
| HLA-DQA1 | ENSG00000196735 | HLA-DQA2 | ENSG00000237541 | HLA-DRA | ENSG00000204287 |
| HLA-DRB4 | ENSG00000227357 | HLA-G | ENSG00000204632 | HLA-H | ENSG00000206341 |
| HNRNPA2B1 | ENSG00000122566 | IFI27 | ENSG00000165949 | IFI30 | ENSG00000216490 |
| IFI35 | ENSG00000068079 | IFI6 | ENSG00000126709 | IFIT1 | ENSG00000185745 |
| IFIT2 | ENSG00000119922 | IFIT3 | ENSG00000119917 | IFIT5 | ENSG00000152778 |
| IFITM1 | ENSG00000185885 | IFITM2 | ENSG00000185201 | IFITM3 | ENSG00000142089 |
| IFNG | ENSG00000111537 | IGHE | ENSG00000211891 | IGHG1 | ENSG00000211896 |
| IGHG4 | ENSG00000211892 | IKBIP | ENSG00000166130 | IL1RN | ENSG00000136689 |
| IL4R | ENSG00000077238 | IL6 | ENSG00000136244 | IL6R | ENSG00000160712 |
| IP6K2 | ENSG00000068745 | IRF1 | ENSG00000125347 | IRF5 | ENSG00000128604 |
| IRF7 | ENSG00000185507 | IRF9 | ENSG00000213928 | ISG15 | ENSG00000187608 |
| ISG20 | ENSG00000172183 | ITGB1 | ENSG00000150093 | LIF | ENSG00000128342 |
| LMNB1 | ENSG00000113368 | MT2A | ENSG00000125148 | MUC1 | ENSG00000185499 |
| MX1 | ENSG00000157601 | MX2 | ENSG00000183486 | NCAM1 | ENSG00000149294 |
| OAS1 | ENSG00000089127 | OAS2 | ENSG00000111335 | OAS3 | ENSG00000111331 |
| OASL | ENSG00000135114 | PML | ENSG00000140464 | PPIA | ENSG00000196262 |
| PSMB8 | ENSG00000204264 | PSME2 | ENSG00000100911 | PTPN2 | ENSG00000175354 |
| RORA | ENSG00000069667 | RORC | ENSG00000143365 | RPLP0 | ENSG00000089157 |
| RSAD2 | ENSG00000134321 | SOCS1 | ENSG00000185338 | SOCS2 | ENSG00000120833 |
| SOCS3 | ENSG00000184557, ENST00000330871 | SP100 | ENSG00000067066 | STAT1 | ENSG00000115415 |
| TRIM14 | ENSG00000106785 | TRIM21 | ENSG00000132109 | TRIM22 | ENSG00000132274 |
| TRIM25 | ENSG00000121060 | TRIM34 | ENSG00000258659 | TRIM38 | ENSG00000112343 |
| TRIM5 | ENSG00000132256 | TRIM6 | ENSG00000121236 | VIM | ENSG00000026025 |
| XAF1 | ENSG00000132530 | | | | |

#### 14. Immune System (R-HSA-168256)

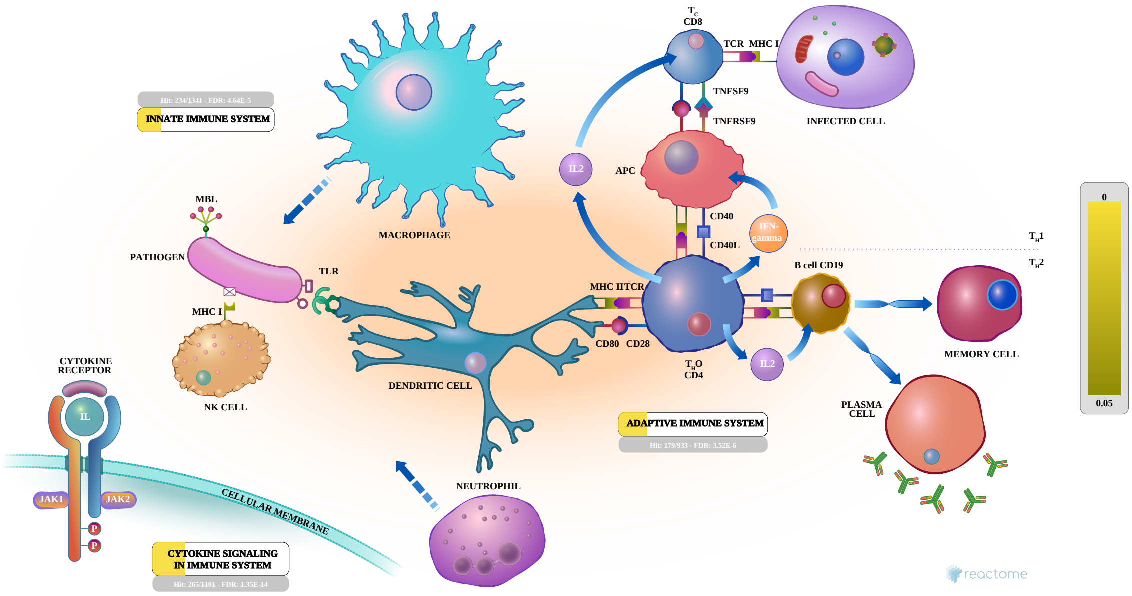


Humans are exposed to millions of potential pathogens daily, through contact, ingestion, and inhalation. Our ability to avoid infection depends on the adaptive immune system and during the first critical hours and days of exposure to a new pathogen, our innate immune system.

##### References

##### Edit history

| Date | Action | Author |
|---|---|---|
| 2005-11-12 | Created | Gillespie ME |
| 2006-03-30 | Authored | Luo F, Ouwehand WH, Gillespie ME, de Bono B |
| 2006-04-19 | Reviewed | Zwaginga JJ, D'Eustachio P, Gay NJ, Gale M Jr |
| 2023-08-26 | Modified | Wright A |

##### 406 submitted entities found in this pathway, mapping to 537 Reactome entities

| Input | UniProt Id | Input | UniProt Id | Input | UniProt Id |
|---|---|---|---|---|---|
| ACTR3 | P61158 | ADA2 | Q9NZK5 | ADAR | P55265 |
| ADGRE3 | Q9BY15 | ADGRG1 | Q86Y34 | AIM2 | O14862 |
| ALDOA | P04075 | ALDOC | P09972 | ALOX15 | P16050 |
| ANPEP | P15144 | AP1B1 | Q10567 | ARF1 | P84077 |
| ARPC1B | O15143 | ATP6V0E2 | Q8NHE4 | B2M | P28068, P61769 |
| BAIAP2 | Q9UQB8 | BIRC2 | Q13489, Q13490 | BRI3 | O95415 |
| BST1 | Q10588 | BST2 | Q10589 | BTF3 | O00478, P78410 |
| BTK | Q06187 | BTN2A2 | Q8WVV5, Q9UIR0 | BTN3A1 | O00481 |
| BTNL8 | Q6UX41 | C1QA | P02745 | C1QB | P02746 |
| C1QC | P02747 | C2 | P06681 | C3AR1 | Q16581 |
| C5 | P01031 | C5AR2 | Q9P296 | CASP1 | P29466 |
| CASP4 | P49662 | CAT | P04040 | CCL2 | P13500 |
| CCL20 | P78556 | CCR1 | P32246 | CCR6 | P51684 |

| Input | UniProt Id | Input | UniProt Id | Input | UniProt Id |
|---|---|---|---|---|---|
| CD160 | O95971 | CD1C | P29017 | CD22 | P20273 |
| CD247 | P20963-1 | CD27 | P26842 | CD274 | Q9NZQ7 |
| CD300E | Q496F6 | CD34 | P28906-1 | CD40LG | P29965 |
| CD68 | P34810 | CD8B | P10966 | CD96 | P40200 |
| CDKN1A | P38936 | CEACAM1 | P13688 | CFB | P00751 |
| CFP | P27918 | CISH | Q9NSE2 | CKAP4 | Q07065 |
| CLEC10A | Q6UW15, Q8IUN9 | CLEC12A | Q5QGZ9 | CLEC2B | Q92478 |
| CLEC4D | Q8WXI8 | CLEC4E | Q9UJ71, Q9ULY5 | CLEC7A | Q9BXN2 |
| CLTC | Q00610, Q00610-1 | CTSF | Q9UBX1 | CTSL | O60911, P07711 |
| CXCL10 | P02778 | CXCR2 | P25024, P25025 | CYBB | P04839 |
| CYFIP2 | Q96F07 | DAPP1 | Q9UN19 | DBNL | Q9UJU6 |
| DBP | Q03518 | DDX58 | O95786 | DHX58 | Q96C10 |
| DNASE1L3 | P49184 | DSC1 | Q08554 | DTX3L | Q8TDB6 |
| DUSP6 | Q16828 | DYNLT1 | P63172 | EDAR | Q9UNE0 |
| EEF1A1 | P68104 | EEF2 | P13639 | EGR1 | P18146 |
| EIF2AK2 | P19525 | FAAP20 | Q6NZ36 | FANCA | O15360 |
| FANCL | Q9NW38 | FBXL8 | Q96CD0 | FBXO30 | Q8TB52 |
| FBXO6 | Q9NRD1 | FBXO7 | Q9Y3I1 | FBXO9 | Q9UK97 |
| FBXW11 | Q9UKB1 | FCAR | P24071 | FCER1A | P12319 |
| FCGR1A | P12314 | FCGR1B | Q92637 | FCGR2A | P12318 |
| FCGR3A | P08637 | FCGR3B | O75015 | FCN1 | O00602 |
| FGL2 | Q14314 | FLT3LG | P49771 | FTH1 | P02794 |
| GBP1 | P32455 | GBP2 | P32456 | GBP3 | Q9H0R5 |
| GBP4 | Q96PP9 | GBP5 | Q96PP8 | GBP6 | Q6ZN66 |
| GDI2 | P50395 | GM2A | P17900 | GNLY | P22749 |
| GOLGA7 | Q7Z5G4 | GPR84 | Q9NQS5 | GRN | P28799 |
| GYG1 | P46976 | GZMM | P51124 | H3C4 | P68431, Q71DI3 |
| HERC4 | Q15034, Q5GLZ8 | HERC5 | Q9UII4 | HERC6 | Q8IVU3 |
| HK3 | P52790 | HLA-A | P04439 | HLA-B | P01889 |
| HLA-C | P10321 | HLA-DQA1 | P01909, P20036 | HLA-DQA2 | P01906 |
| HLA-DRA | P01903 | HLA-DRB4 | P13762 | HLA-G | P17693 |
| HLA-H | P01893 | HNRNPA2B1 | P22626 | HP | P00738 |
| HSPA1A | P0DMV8 | HSPA6 | P17066 | ICOSLG | O75144 |
| IFI16 | Q16666 | IFI27 | P40305 | IFI30 | P13284 |
| IFI35 | P80217 | IFI6 | P09912 | IFIH1 | Q9BYX4 |
| IFIT1 | P09914 | IFIT2 | P09913 | IFIT3 | O14879 |
| IFIT5 | Q13325 | IFITM1 | P13164 | IFITM2 | Q01629 |
| IFITM3 | P13164, Q01628 | IFNG | P01579 | IGHE | P01854 |
| IGHG1 | P01857, P01861 | IGHG3 | P01860 | IGHG4 | P01861 |
| IGHV1-69 | P01742 | IGHV3-11 | P01762 | IGHV3-23 | P01764 |
| IGHV3-30 | P01768 | IGHV3-33 | P01772 | IGHV4-34 | P06331 |
| IGHV4-39 | P01824 | IGHV4-59 | P01825 | IGKC | P01834 |
| IGKV1-12 | A0A0C4DH73 | IGKV1-17 | P01599 | IGKV1-33 | P01594 |
| IGKV1-5 | P01602 | IGKV1D-39 | P04432 | IGKV3-11 | P04433 |
| IGKV3-15 | P01624 | IGKV3-20 | P01619 | IGLC1 | P0CG04 |
| IGLC2 | P0DOY2 | IGLC3 | P0DOY3 | IGLC7 | A0M8Q6 |
| IGLV1-40 | P01703, Q5NV69 | IGLV1-44 | P01699, Q5NV81 | IGLV1-47 | P01700 |
| IGLV1-51 | P01701 | IGLV2-14 | P01704 | IGLV2-23 | P01705, Q5NV89 |
| IGLV3-1 | P01715 | IGLV3-19 | P01714 | IGLV3-21 | P80748 |

| Input | UniProt Id | Input | UniProt Id | Input | UniProt Id |
|---|---|---|---|---|---|
| IGLV3-25 | P01717, Q5NV90 | IGLV6-57 | P01721 | IKBIP | Q70UQ0 |
| IL11RA | P08887, Q14626 | IL15 | P40933 | IL1RN | P18510 |
| IL27 | Q8NEV9 | IL2RB | P14784 | IL37 | Q9NZH6 |
| IL4R | P24394, Q01113 | IL5RA | P15509, Q01344 | IL6 | P05231 |
| IL6R | P08887, P08887-2 | IL7R | P16871 | IMPDH1 | P20839 |
| IP6K2 | Q9UHH9 | IQGAP1 | P46940 | IRF1 | P10914 |
| IRF5 | Q13568 | IRF7 | Q92985 | IRF9 | Q00978 |
| ISG15 | P05161 | ISG20 | Q96AZ6 | ITGA4 | P13612 |
| ITGB1 | P05556 | JAK2 | O60674 | JUP | P14923 |
| KIR2DL1 | P43626, P43629 | KIR3DL1 | P43629 | KLHL3 | Q9UH77 |
| KLRB1 | Q12918 | KLRF1 | Q9NZS2 | KLRG1 | Q96E93 |
| KPNB1 | Q14974 | KRT1 | P04264 | LAG3 | P18627 |
| LAMP3 | P34810 | LAMTOR1 | Q6IAA8 | LAP3 | Q13867 |
| LGALS9 | O00182 | LGMN | Q99538 | LIF | P15018 |
| LILRA4 | P59901 | LILRA5 | A6NI73, Q8N149 | LILRA6 | Q6PI73 |
| LILRB1 | Q8NHL6 | LILRB2 | Q8N423 | LILRB4 | Q8N149, Q8NHJ6 |
| LMNB1 | P20700 | LTB | Q06643 | LY96 | Q9Y6Y9 |
| MAL | P01732, P58753 | MAP2K3 | P46734 | MAP2K6 | P52564 |
| MAP3K1 | Q13233 | MAPK3 | P27361 | MBP | P13727 |
| MEF2C | Q06413 | MEFV | O15553 | MICB | Q29980 |
| MMP8 | P22894 | MNDA | P41218 | MT1A | P02795, P53803 |
| MT2A | P02795 | MUC1 | P15941 | MUC6 | Q6W4X9 |
| MX1 | P20591 | MX2 | P20592 | MYD88 | Q99836 |
| NAPA | Q9Y2A7 | NCAM1 | P13591 | NCF1 | P14598 |
| NCF2 | P19878 | NCR3 | O14931 | NHLRC3 | Q5JS37 |
| NLRC5 | Q86WI3 | NLRX1 | Q86UT6 | NMUR1 | O94822 |
| NOD2 | Q9HC29 | NPC2 | P61916 | NPDC1 | Q9NQX5 |
| NUP188 | Q5SRE5 | OAS1 | P00973 | OAS2 | P29728 |
| OAS3 | Q9Y6K5 | OASL | Q15646 | P2RX1 | P51575 |
| PDCD1LG2 | Q9BQ51 | PDPK1 | O15530 | PGAM1 | P18669 |
| PI3 | P19957 | PIK3AP1 | Q6ZUJ8 | PIK3CA | P42336 |
| PILRA | Q9UKJ1 | PIN1 | Q13526 | PJA1 | Q8NG27 |
| PLAUR | Q03405 | PLPP5 | Q8NEB5 | PML | P29590 |
| POLR1E | O15160 | POLR3H | Q9Y535 | PPBP | P02775 |
| PPIA | P62937, Q9UNP9 | PPP2R5D | Q14738 | PPP3CA | Q08209 |
| PPP3CB | P16298 | PPP3CC | P16298 | PPP3R1 | P63098 |
| PRKCB | P05771 | PRKCD | Q05655 | PRKCE | Q02156 |
| PRKCSH | P14314 | PRKDC | P78527 | PRR5 | P85299 |
| PSMB8 | P28062 | PSMB9 | P28065 | PSME1 | Q06323 |
| PSME2 | Q9UL46 | PTK2B | Q14289 | PTPN2 | P17706, P17706-2 |
| PTPN22 | Q9Y2R2 | PTPRN2 | Q92932 | RAB24 | Q969Q5 |
| RAB3D | O95716 | RBCK1 | Q9BYM8 | REG4 | Q06141 |
| REL | Q01201, Q04864 | RIGI | O95786 | RLIM | Q9NVW2 |
| RNASE2 | P10153 | RNF213 | Q63HN8 | RNF38 | Q9P000 |
| RORA | P35398 | RORC | P51449 | RPLP0 | P05388 |
| RPN2 | Q99460 | RPS27A | P62979, P62987 | RSAD2 | Q8WXG1 |
| S100B | P04271 | SDCBP | O00560 | SEC24D | O94855 |
| SEC61G | P60059 | SELL | P14151 | SERPING1 | P05155 |
| SH2D1B | O14796 | SIGIRR | Q6IA17 | SIGLEC1 | Q9BZZ2 |

| Input | UniProt Id | Input | UniProt Id | Input | UniProt Id |
|---|---|---|---|---|---|
| SIGLEC14 | Q08ET2 | SIGLEC5 | O15389, O43699 | SIGLEC7 | Q9Y286 |
| SIGLEC8 | Q96PQ1, Q9NYZ4 | SIRPA | P78324 | SLC15A4 | Q8N697 |
| SLC27A3 | O14975 | SOCS1 | O15524 | SOCS2 | O14508 |
| SOCS3 | O14543 | SOS2 | Q07890 | SP100 | P23497 |
| SP110 | P23497 | SPSB2 | Q99619 | SQSTM1 | Q13501 |
| STAT1 | P42224, P42224-1, P42224-2 | STAT2 | P52630 | SYK | P43405 |
| TAB3 | Q8N5C8 | TAP1 | Q03518 | TAP2 | Q03519 |
| TARBP2 | Q15633 | TCN1 | P20061 | TIFA | Q96CG3 |
| TIMP2 | P16035 | TLR1 | Q15399 | TLR2 | O60603 |
| TLR4 | O00206 | TLR5 | O60602 | TLR7 | Q9NYK1 |
| TLR8 | Q9NR97 | TLR9 | Q9NR96 | TNFAIP6 | P98066 |
| TNFRSF17 | Q02223 | TNFRSF25 | Q93038 | TNFSF13 | O75888 |
| TNFSF13B | Q9Y275 | TNFSF14 | O43557 | TOM1L2 | O60784 |
| TRAF3 | Q13114 | TREML1 | Q86YW5 | TREX1 | Q9NSU2 |
| TRIM14 | Q14142 | TRIM21 | P19474 | TRIM22 | Q8IYM9 |
| TRIM25 | Q14258 | TRIM34 | Q9BYJ4 | TRIM38 | O00635 |
| TRIM5 | Q9C035 | TRIM56 | Q9BRZ2 | TRIM6 | Q9C030 |
| TRIM69 | Q86WT6 | TRIP12 | Q14669 | TUBA4A | P68366 |
| TXK | P42681 | TXNDC5 | Q8NBS9 | UBA6 | A0AVT1 |
| UBA7 | P41226 | UBE2C | O00762 | UBE2G1 | P62253 |
| UBE2J2 | Q8N2K1 | UBE2L6 | O14933 | UBOX5 | O94941 |
| UFL1 | O94874 | UNC93B1 | Q9H1C4 | USP18 | Q3LFD5, Q9UMW8 |
| USP41 | Q3LFD5 | VIM | P08670 | VRK3 | Q8IV63 |
| WIPF1 | O43516 | WSB1 | Q9Y6I7 | XAF1 | Q6GPH4 |
| ZBP1 | Q9H171 | | | | |

| Input | Ensembl Id | Input | Ensembl Id | Input | Ensembl Id |
|---|---|---|---|---|---|
| ADAR | ENSG00000160710 | ALOX15 | ENSG00000161905 | ARF1 | ENSG00000143761 |
| B2M | ENSG00000166710 | BST2 | ENSG00000130303 | CCL2 | ENSG00000108691 |
| CCL20 | ENSG00000115009 | CCR1 | ENSG00000163823 | CDKN1A | ENSG00000124762 |
| CISH | ENSG00000114737 | CXCL10 | ENSG00000169245 | EGR1 | ENSG00000120738 |
| EIF2AK2 | ENSG00000055332 | FCGR1A | ENSG00000150337 | FCGR1B | ENSG00000198019 |
| GBP1 | ENSG00000117228 | GBP2 | ENSG00000162645 | GBP3 | ENSG00000117226 |
| GBP4 | ENSG00000162654 | GBP5 | ENSG00000154451 | GBP6 | ENSG00000183347 |
| HLA-A | ENSG00000206503 | HLA-B | ENSG00000234745 | HLA-C | ENSG00000204525 |
| HLA-DQA1 | ENSG00000196735 | HLA-DQA2 | ENSG00000237541 | HLA-DRA | ENSG00000204287 |
| HLA-DRB4 | ENSG00000227357 | HLA-G | ENSG00000204632 | HLA-H | ENSG00000206341 |
| HNRNPA2B1 | ENSG00000122566 | IFI27 | ENSG00000165949 | IFI30 | ENSG00000216490 |
| IFI35 | ENSG00000068079 | IFI6 | ENSG00000126709 | IFIT1 | ENSG00000185745 |
| IFIT2 | ENSG00000119922 | IFIT3 | ENSG00000119917 | IFIT5 | ENSG00000152778 |
| IFITM1 | ENSG00000185885 | IFITM2 | ENSG00000185201 | IFITM3 | ENSG00000142089 |
| IFNG | ENSG00000111537 | IGHE | ENSG00000211891 | IGHG1 | ENSG00000211896 |
| IGHG4 | ENSG00000211892 | IKBIP | ENSG00000166130 | IL1RN | ENSG00000136689 |
| IL4R | ENSG00000077238 | IL6 | ENSG00000136244 | IL6R | ENSG00000160712 |
| IP6K2 | ENSG00000068745 | IRF1 | ENSG00000125347 | IRF5 | ENSG00000128604 |
| IRF7 | ENSG00000185507 | IRF9 | ENSG00000213928 | ISG15 | ENSG00000187608 |
| ISG20 | ENSG00000172183 | ITGB1 | ENSG00000150093 | LIF | ENSG00000128342 |
| LMNB1 | ENSG00000113368 | MT2A | ENSG00000125148 | MUC1 | ENSG00000185499 |
| MX1 | ENSG00000157601 | MX2 | ENSG00000183486 | NCAM1 | ENSG00000149294 |

| Input | Ensembl Id | Input | Ensembl Id | Input | Ensembl Id |
|---|---|---|---|---|---|
| OAS1 | ENSG00000089127 | OAS2 | ENSG00000111335 | OAS3 | ENSG00000111331 |
| OASL | ENSG00000135114 | PML | ENSG00000140464 | PPIA | ENSG00000196262 |
| PSMB8 | ENSG00000204264 | PSME2 | ENSG00000100911 | PTPN2 | ENSG00000175354 |
| RORA | ENSG00000069667 | RORC | ENSG00000143365 | RPLP0 | ENSG00000089157 |
| RSAD2 | ENSG00000134321 | SOCS1 | ENSG00000185338 | SOCS2 | ENSG00000120833 |
| SOCS3 | ENSG00000184557, ENST00000330871 | SP100 | ENSG00000067066 | STAT1 | ENSG00000115415 |
| TRIM14 | ENSG00000106785 | TRIM21 | ENSG00000132109 | TRIM22 | ENSG00000132274 |
| TRIM25 | ENSG00000121060 | TRIM34 | ENSG00000258659 | TRIM38 | ENSG00000112343 |
| TRIM5 | ENSG00000132256 | TRIM6 | ENSG00000121236 | VIM | ENSG00000026025 |
| XAF1 | ENSG00000132530 | | | | |

#### 15. Eukaryotic Translation Termination (R-HSA-72764)

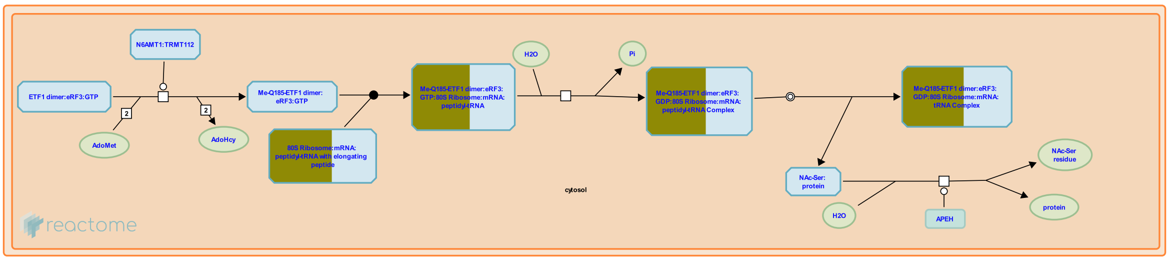


**Cellular compartments:** cytosol.

The arrival of any of the three stop codons (UAA, UAG and UGA) into the ribosomal A-site triggers the binding of a release factor (RF) to the ribosome and subsequent polypeptide chain release. In eukaryotes, the RF is composed of two proteins, eRF1 and eRF3. eRF1 is responsible for the hydrolysis of the peptidyl-tRNA, while eRF3 provides a GTP-dependent function. The ribosome releases the mRNA and dissociates into its two complex subunits, which can reassemble on another molecule to begin a new round of protein synthesis. It should be noted that at present, there is no factor identified in eukaryotes that would be the functional equivalent of the bacterial ribosome release (or recycling) factor, RRF, that catalyzes dissociation of the ribosome from the mRNA following release of the polypeptide

##### References

Salas-Marco J & Bedwell DM (2004). GTP hydrolysis by eRF3 facilitates stop codon decoding during eukaryotic translation termination. Mol Cell Biol, 24, 7769-78.

Ichikawa S & Kaji A (1989). Molecular cloning and expression of ribosome releasing factor. J Biol Chem, 264, 20054-9.

##### Edit history

| Date | Action | Author |
|---|---|---|
| 2004-11-09 | Authored | Bedwell DM |
| 2005-01-03 | Created | Merrick WC, Bedwell DM |
| 2023-08-24 | Edited | Gillespie ME |
| 2023-08-31 | Modified | Wright A |

##### 52 submitted entities found in this pathway, mapping to 66 Reactome entities

| Input | UniProt Id | Input | UniProt Id | Input | UniProt Id |
|---|---|---|---|---|---|
| AGRN | P32969 | FAU | P62861 | RPL10 | P27635, Q96L21 |
| RPL10A | P61313, P62906 | RPL13 | P26373, P40429 | RPL13A | P40429, P61313 |
| RPL15 | P61313 | RPL18 | Q07020 | RPL21 | P46778 |
| RPL27A | P46776, P61353 | RPL29 | P47914 | RPL3 | P39023, Q92901 |
| RPL30 | P62888 | RPL31 | P62899 | RPL32 | P62888, P62910 |
| RPL34 | P49207, P62899 | RPL35 | P42766 | RPL35A | P18077 |
| RPL36 | Q9Y3U8 | RPL38 | P63173 | RPL3L | Q92901 |
| RPL4 | P36578 | RPL5 | P46777 | RPL7 | P18124 |
| RPL7A | P18124, P62424 | RPL9P8 | P32969 | RPLP0 | P05388 |
| RPLP2 | P05387 | RPS10 | P46783 | RPS11 | P62280 |

| Input | UniProt Id | Input | UniProt Id | Input | UniProt Id |
|---|---|---|---|---|---|
| RPS12 | P25398 | RPS13 | P62277 | RPS14 | P62263 |
| RPS15 | P62841 | RPS15A | P62244 | RPS16 | P62249 |
| RPS17 | P08708 | RPS18 | P62269 | RPS2 | P15880, P46782 |
| RPS23 | P62266 | RPS27 | P42677, Q71UM5 | RPS27A | P42677, P62979, P62987 |
| RPS29 | P62273 | RPS3 | P23396 | RPS3A | P61247 |
| RPS4X | P62701, Q8TD47 | RPS5 | P46782 | RPS6 | P62753 |
| RPS8 | P62241 | RPS9 | P46781 | RPSA | P08865 |
| SOS2 | P42766 | | | | |

#### 16. Response of EIF2AK4 (GCN2) to amino acid deficiency (R-HSA-9633012)

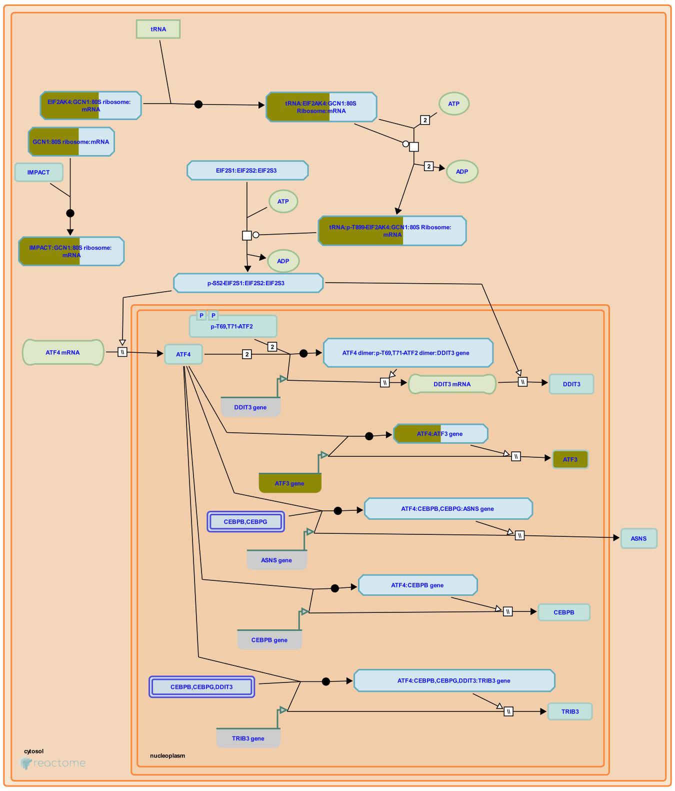


EIF2AK4 (GCN2) senses amino acid deficiency by binding uncharged tRNAs near the ribosome and responds by phosphorylating EIF2S1, the alpha subunit of the translation initiation factor EIF2 (inferred from yeast homologs and mouse homologs, reviewed in Chaveroux et al. 2010, Castilho et al. 2014, Gallinetti et al. 2013, Bröer and Bröer 2017, Wek 2018). Phosphorylated EIF2S1 reduces translation of most mRNAs but increases translation of downstream ORFs in mRNAs such as ATF4 that contain upstream ORFs (inferred from mouse homologs in Vattem and Wek 2004, reviewed in Hinnebusch et al. 2016, Sonenberg and Hinnebusch 2009). ATF4, in turn, activates expression of genes involved in responding to amino acid deficiency such as DDIT3 (CHOP), ASNS (asparagine synthetase), CEBPB, and ATF3 (reviewed in Kilberg et al. 2012, Wortel et al. 2017). In mice, EIF2AK4 in the brain may responsible for avoidance of diets lacking essential amino acids (Hao et al. 2005, Maurin et al. 2005, see also Leib and Knight 2015, Gietzen et al. 2016, reviewed in Dever and Hinnebusch 2005).

EIF2AK4 is bound to both the ribosome and GCN1, which is required for activation of EIF2AK4 and may act by shuttling uncharged tRNAs from the A site of the ribosome to EIF2AK4. Upon binding tRNA, EIF2AK4 trans-autophosphorylates. Phosphorylated EIF2AK4 then phosphorylates EIF2S1 on serine-52, the same serine residue phosphorylated by other kinases of the integrated stress response: EIF2AK1 (HRI, activated by heme deficiency and other stresses), EIF2AK2 (PKR, activated by double-stranded RNA), and EIF2AK3 (PERK, activated by unfolded proteins) (reviewed in Hinnebusch 1994, Wek et al. 2006, Donnelly et al. 2013, Pakos-Zebrucka et al. 2016, Wek 2018),

##### References


Koryga I, Pakos-Zebrucka K, Gorman AM, Samali A, Mnich K & Ljujic M (2016). The integrated stress response. EMBO Rep., 17, 1374-1395.

McGrath BC, Ross-Inta CM, Hao S, Koehnle TJ, McDaniel BJ, Sharp JW, ... Anthony TG (2005). Uncharged tRNA and sensing of amino acid deficiency in mammalian piriform cortex. Science, 307 , 1776-8.

Leib DE & Knight ZA (2015). Re-examination of Dietary Amino Acid Sensing Reveals a GCN2-Independent Mechanism. Cell Rep, 13, 1081-1089.

Shanmugam R, Himme BM, Sattlegger E, Silva RC, Ramesh R & Castilho BA (2014). Keeping the eIF2 alpha kinase Gcn2 in check. Biochim. Biophys. Acta, 1843, 1948-68.

Mitchell JR, Gallinetti J & Harputlugil E (2013). Amino acid sensing in dietary-restriction-mediated longevity: roles of signal-transducing kinases GCN2 and TOR. Biochem. J., 449, 1-10.

#### Edit history

| Date | Action | Author |
|---|---|---|
| 2018-12-28 | Edited | May B |
| 2018-12-28 | Authored | May B |
| 2018-12-28 | Created | May B |
| 2019-09-15 | Reviewed | Bruhat A |
| 2019-11-20 | Reviewed | Staschke KA |
| 2023-03-08 | Modified | Matthews L |

#### 53 submitted entities found in this pathway, mapping to 68 Reactome entities

| Input | UniProt Id | Input | UniProt Id | Input | UniProt Id |
|---|---|---|---|---|---|
| AGRN | P32969 | ATF3 | P18847 | FAU | P62861 |
| RPL10 | P27635, Q96L21 | RPL10A | P61313, P62906 | RPL13 | P26373, P40429 |
| RPL13A | P40429, P61313 | RPL15 | P61313 | RPL18 | Q07020 |
| RPL21 | P46778 | RPL27A | P46776, P61353 | RPL29 | P47914 |
| RPL3 | P39023, Q92901 | RPL30 | P62888 | RPL31 | P62899 |
| RPL32 | P62888, P62910 | RPL34 | P49207, P62899 | RPL35 | P42766 |
| RPL35A | P18077 | RPL36 | Q9Y3U8 | RPL38 | P63173 |
| RPL3L | Q92901 | RPL4 | P36578 | RPL5 | P46777 |
| RPL7 | P18124 | RPL7A | P18124, P62424 | RPL9P8 | P32969 |
| RPLP0 | P05388 | RPLP2 | P05387 | RPS10 | P46783 |
| RPS11 | P62280 | RPS12 | P25398 | RPS13 | P62277 |
| RPS14 | P62263 | RPS15 | P62841 | RPS15A | P62244 |
| RPS16 | P62249 | RPS17 | P08708 | RPS18 | P62269 |
| RPS2 | P15880, P46782 | RPS23 | P62266 | RPS27 | P42677, Q71UM5 |
| RPS27A | P42677, P62979, P62987 | RPS29 | P62273 | RPS3 | P23396 |
| RPS3A | P61247 | RPS4X | P62701, Q8TD47 | RPS5 | P46782 |
| RPS6 | P62753 | RPS8 | P62241 | RPS9 | P46781 |
| RPSA | P08865 | SOS2 | P42766 | | |

| Input | Ensembl Id |
|---|---|
| ATF3 | ENSG00000162772 |

#### 17. Selenocysteine synthesis (R-HSA-2408557)

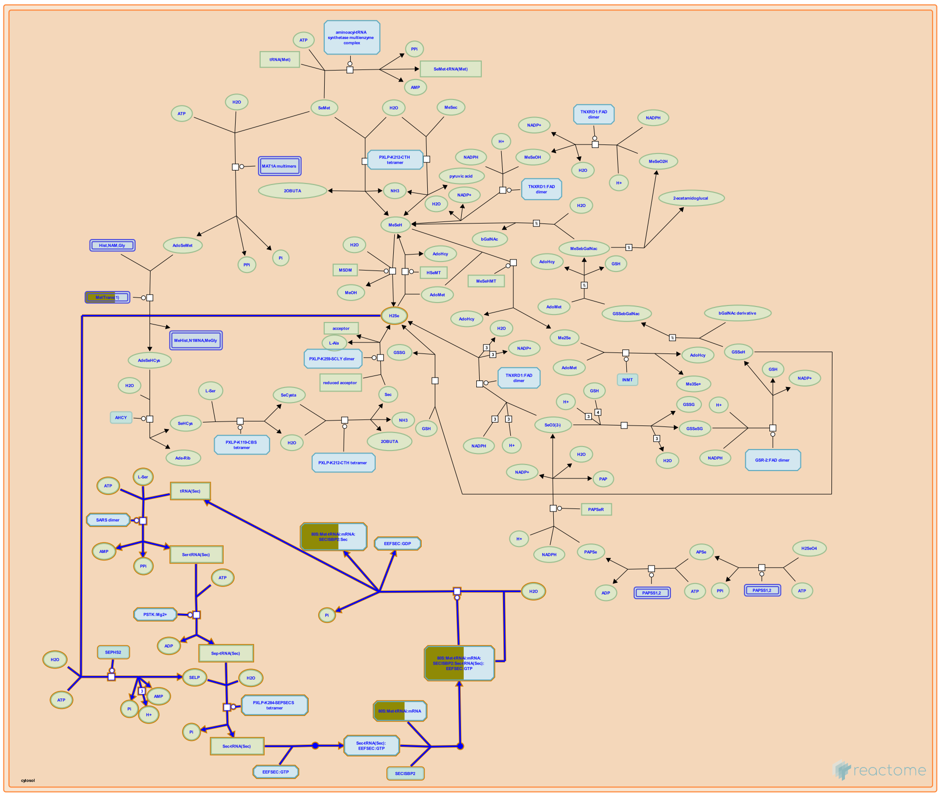

Selenocysteine, the 21st genetically encoded amino acid, is the major form of the antioxidant trace element selenium in the human body. In eukaryotes and archaea its synthesis proceeds through a phosphorylated intermediate in a tRNA-dependent fashion. The final step of selenocysteine formation is catalyzed by O-phosphoseryl-tRNA:selenocysteinyl-tRNA synthase (SEPSECS) that converts phosphoseryl-tRNA(Sec) to selenocysteinyl-tRNA(Sec).

##### References


Donovan J & Copeland PR (2010). Threading the needle: getting selenocysteine into proteins. Antioxid. Redox Signal., 12, 881-92.

Palioura S, Herkel J, Simonovic M, Lohse AW & Söll D (2010). Human SepSecS or SLA/LP: selenocysteine formation and autoimmune hepatitis. Biol. Chem., 391, 771-6.

Sheppard K, Yuan J, Devine KM, Jester B, Söll D & Hohn MJ (2008). From one amino acid to another: tRNA-dependent amino acid biosynthesis. Nucleic Acids Res., 36, 1813-25.


##### Edit history

| Date | Action | Author |
|---|---|---|
| 2012-07-17 | Created | Williams MG |
| 2014-05-06 | Authored | Williams MG |
| 2015-08-29 | Edited | D'Eustachio P |
| 2015-08-30 | Reviewed | Rush MG |
| 2023-08-26 | Modified | Wright A |

##### 52 submitted entities found in this pathway, mapping to 66 Reactome entities

| Input | UniProt Id | Input | UniProt Id | Input | UniProt Id |
|---|---|---|---|---|---|
| AGRN | P32969 | FAU | P62861 | RPL10 | P27635, Q96L21 |
| RPL10A | P61313, P62906 | RPL13 | P26373, P40429 | RPL13A | P40429, P61313 |
| RPL15 | P61313 | RPL18 | Q07020 | RPL21 | P46778 |
| RPL27A | P46776, P61353 | RPL29 | P47914 | RPL3 | P39023, Q92901 |
| RPL30 | P62888 | RPL31 | P62899 | RPL32 | P62888, P62910 |
| RPL34 | P49207, P62899 | RPL35 | P42766 | RPL35A | P18077 |
| RPL36 | Q9Y3U8 | RPL38 | P63173 | RPL3L | Q92901 |
| RPL4 | P36578 | RPL5 | P46777 | RPL7 | P18124 |
| RPL7A | P18124, P62424 | RPL9P8 | P32969 | RPLP0 | P05388 |
| RPLP2 | P05387 | RPS10 | P46783 | RPS11 | P62280 |
| RPS12 | P25398 | RPS13 | P62277 | RPS14 | P62263 |
| RPS15 | P62841 | RPS15A | P62244 | RPS16 | P62249 |
| RPS17 | P08708 | RPS18 | P62269 | RPS2 | P15880, P46782 |
| RPS23 | P62266 | RPS27 | P42677, Q71UM5 | RPS27A | P42677, P62979, P62987 |
| RPS29 | P62273 | RPS3 | P23396 | RPS3A | P61247 |
| RPS4X | P62701, Q8TD47 | RPS5 | P46782 | RPS6 | P62753 |
| RPS8 | P62241 | RPS9 | P46781 | RPSA | P08865 |
| SOS2 | P42766 | | | | |

#### 18. Viral mRNA Translation (R-HSA-192823)

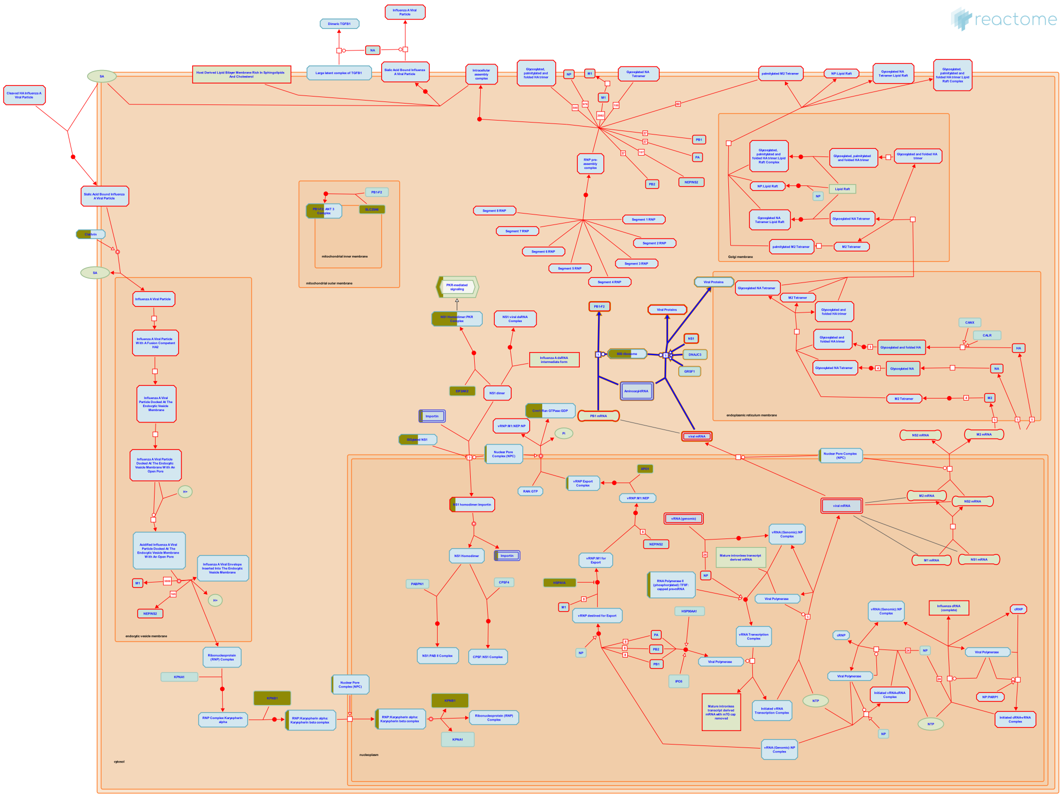

**Cellular compartments:** cytosol.

**Diseases:** influenza.

Spliced and unspliced viral mRNA in the cytoplasm are translated by host cell ribosomal translation machinery (reviewed in Kash, 2006). At least ten viral proteins are synthesized: HA, NA, PB1, PB2, PA, NP, NS1, NEP/NS2, M1, and M2. Viral mRNA translation is believed to be enhanced by conserved 5'UTR sequences that interact with the ribosomal machinery and at least one cellular RNA-binding protein, G-rich sequence factor 1 (GRSF-1), has been found to specifically interact with the viral 5' UTRs. (Park, 1995; Park, 1999). The viral NS1 protein and the cellular protein P58(IPK) enhance viral translation indirectly by preventing the activation of the translational inhibitor PKR (Salvatore, 2002; Goodman, 2006). The viral NS1 protein has also been proposed to specifically enhance translation through interaction with host poly(A)-binding protein 1 (PABP1) (Burgui, 2003). Simultaneously, host cell protein synthesis is downregulated in influenza virus infection through still uncharacterized mechanisms (Katze, 1986; Garfinkel, 1992; Kash, 2006). In most human influenza A strains (such as PR8), the PB1 mRNA segment is capable of producing a second protein, PB1-F2, from a short +1 open reading frame initiating downstream of the PB1 ORF initiation codon (Chen, 2001).

##### References


Korth MJ, Katze MG, Kash JC & Goodman AG (2006). Hijacking of the host-cell response and translational control during influenza virus infection. Virus Res, 119, 111-20.


##### Edit history

| Date | Action | Author |
|---|---|---|
| 2007-02-07 | Created | Gillespie ME |

| Date | Action | Author |
|---|---|---|
| 2007-02-13 | Reviewed | Squires B |
| 2007-02-13 | Authored | Garcia-Sastre A, Bortz E |
| 2023-08-31 | Modified | Wright A |

#### 52 submitted entities found in this pathway, mapping to 66 Reactome entities

| Input | UniProt Id | Input | UniProt Id | Input | UniProt Id |
|---|---|---|---|---|---|
| AGRN | P32969 | FAU | P62861 | RPL10 | P27635, Q96L21 |
| RPL10A | P61313, P62906 | RPL13 | P26373, P40429 | RPL13A | P40429, P61313 |
| RPL15 | P61313 | RPL18 | Q07020 | RPL21 | P46778 |
| RPL27A | P46776, P61353 | RPL29 | P47914 | RPL3 | P39023, Q92901 |
| RPL30 | P62888 | RPL31 | P62899 | RPL32 | P62888, P62910 |
| RPL34 | P49207, P62899 | RPL35 | P42766 | RPL35A | P18077 |
| RPL36 | Q9Y3U8 | RPL38 | P63173 | RPL3L | Q92901 |
| RPL4 | P36578 | RPL5 | P46777 | RPL7 | P18124 |
| RPL7A | P18124, P62424 | RPL9P8 | P32969 | RPLP0 | P05388 |
| RPLP2 | P05387 | RPS10 | P46783 | RPS11 | P62280 |
| RPS12 | P25398 | RPS13 | P62277 | RPS14 | P62263 |
| RPS15 | P62841 | RPS15A | P62244 | RPS16 | P62249 |
| RPS17 | P08708 | RPS18 | P62269 | RPS2 | P15880, P46782 |
| RPS23 | P62266 | RPS27 | P42677, Q71UM5 | RPS27A | P42677, P62979, P62987 |
| RPS29 | P62273 | RPS3 | P23396 | RPS3A | P61247 |
| RPS4X | P62701, Q8TD47 | RPS5 | P46782 | RPS6 | P62753 |
| RPS8 | P62241 | RPS9 | P46781 | RPSA | P08865 |
| SOS2 | P42766 | | | | |

#### 19. Eukaryotic Translation Initiation (R-HSA-72613)

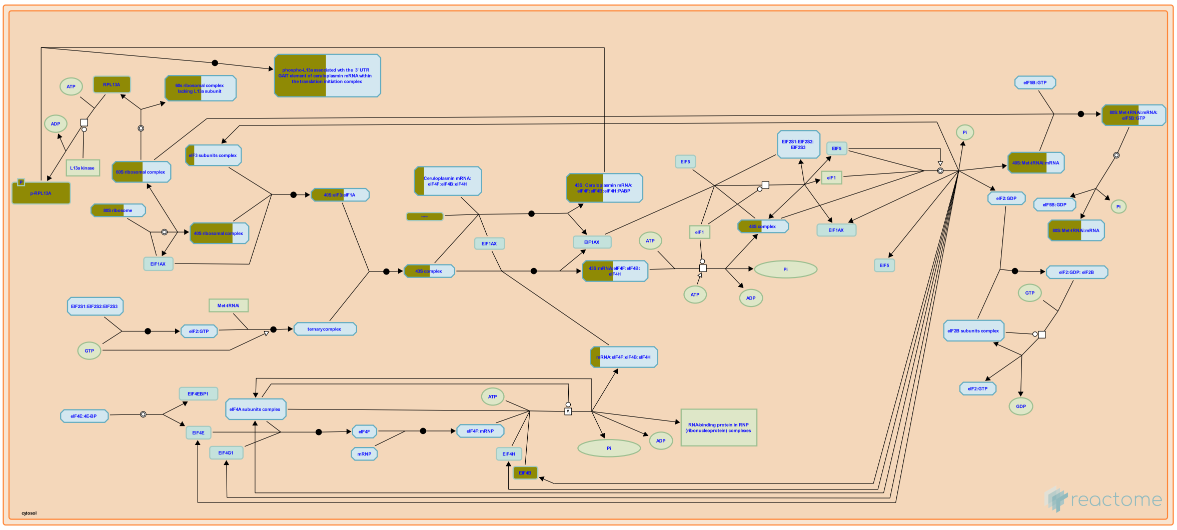

Initiation of translation in the majority of eukaryotic cellular mRNAs depends on the 5'-cap (m7GpppN) and involves ribosomal scanning of the 5' untranslated region (5'-UTR) for an initiating AUG start codon. Therefore, this mechanism is often called cap-dependent translation initiation. Proximity to the cap, as well as the nucleotides surrounding an AUG codon, influence the efficiency of the start site recognition during the scanning process. However, if the recognition site is poor enough, scanning ribosomal subunits will ignore and skip potential starting AUGs, a phenomenon called leaky scanning. Leaky scanning allows a single mRNA to encode several proteins that differ in their amino-termini. Merrick (2010) provides an overview of this process and hghlights several features of it that remain incompletely understood.

Several eukaryotic cell and viral mRNAs initiate translation by an alternative mechanism that involves internal initiation rather than ribosomal scanning. These mRNAs contain complex nucleotide sequences, called internal ribosomal entry sites, where ribosomes bind in a cap-independent manner and start translation at the closest downstream AUG codon.

Initiation on several viral and cellular mRNAs is cap-independent and is mediated by binding of the ribosome to internal ribosome entry site (IRES) elements. These elements are often found in characteristically long structured regions on the 5'-UTR of an mRNA that may or may not have regulatory upstream open reading frames (uORFs). Both of these features on the 5'-end of the mRNA hinder ribosomal scanning, and thus promote a cap-independent translation initiation mechanism. IRESs act as specific translational enhancers that allow translation initiation to occur in response to specific stimuli and under the control of different trans-acting factors, as for example when cap-dependent protein synthesis is shut off during viral infection. Such regulatory elements have been identified in the mRNAs of growth factors, protooncogenes, angiogenesis factors, and apoptosis regulators, which are translated under a variety of stress conditions, including hypoxia, serum deprivation, irradiation and apoptosis. Thus, cap-independent translational control might have evolved to regulate cellular responses in acute but transient stress conditions that would otherwise lead to cell death, while the same mechanism is of major importance for viral mRNAs to bypass the shutting-off of host protein synthesis after infection. Encephalomyocarditis virus (EMCV) and hepatitis C virus exemplify two distinct mechanisms of IRES-mediated initiation. In contrast to cap-dependent initiation, the eIF4A and eIF4G subunits of eIF4F bind immediately upstream of the EMCV initiation codon and promote binding of a 43S complex. Accordingly, EMCV initiation does not involve scanning and does not require eIF1, eIF1A, and the eIF4E subunit of eIF4F. Nonetheless, initiation on some EMCV-like IRESs requires additional non-canonical initiation factors, which alter IRES conformation and promote binding of eIF4A/eIF4G. Initiation on the hepatitis C virus IRES is simpler: a 43S complex containing only eIF2 and eIF3 binds directly to the initiation codon as a result of specific interaction of the IRES and the 40S subunit.

#### References


Merrick WC (2010). Eukaryotic protein synthesis: still a mystery. J Biol Chem, 285, 21197-201.


#### Edit history

| Date | Action | Author |
|---|---|---|
| 2002-12-16 | Created | Merrick WC |
| 2023-08-26 | Modified | Wright A |

#### 56 submitted entities found in this pathway, mapping to 70 Reactome entities

| Input | UniProt Id | Input | UniProt Id | Input | UniProt Id |
|---|---|---|---|---|---|
| AGRN | P32969 | EIF3F | O00303 | EIF3L | Q9Y262 |
| EIF4B | P23588 | FAU | P62861 | PABPC1 | P11940 |
| RPL10 | P27635, Q96L21 | RPL10A | P61313, P62906 | RPL13 | P26373, P40429 |
| RPL13A | P40429, P61313 | RPL15 | P61313 | RPL18 | Q07020 |
| RPL21 | P46778 | RPL27A | P46776, P61353 | RPL29 | P47914 |
| RPL3 | P39023, Q92901 | RPL30 | P62888 | RPL31 | P62899 |
| RPL32 | P62888, P62910 | RPL34 | P49207, P62899 | RPL35 | P42766 |
| RPL35A | P18077 | RPL36 | Q9Y3U8 | RPL38 | P63173 |
| RPL3L | Q92901 | RPL4 | P36578 | RPL5 | P46777 |
| RPL7 | P18124 | RPL7A | P18124, P62424 | RPL9P8 | P32969 |
| RPLP0 | P05388 | RPLP2 | P05387 | RPS10 | P46783 |
| RPS11 | P62280 | RPS12 | P25398 | RPS13 | P62277 |
| RPS14 | P62263 | RPS15 | P62841 | RPS15A | P62244 |
| RPS16 | P62249 | RPS17 | P08708 | RPS18 | P62269 |

| Input | UniProt Id | Input | UniProt Id | Input | UniProt Id |
|---|---|---|---|---|---|
| RPS2 | P15880, P46782 | RPS23 | P62266 | RPS27 | P42677, Q71UM5 |
| RPS27A | P42677, P62979, P62987 | RPS29 | P62273 | RPS3 | P23396 |
| RPS3A | P61247 | RPS4X | P62701, Q8TD47 | RPS5 | P46782 |
| RPS6 | P62753 | RPS8 | P62241 | RPS9 | P46781 |
| RPSA | P08865 | SOS2 | P42766 | | |

#### 20. Cap-dependent Translation Initiation (R-HSA-72737)

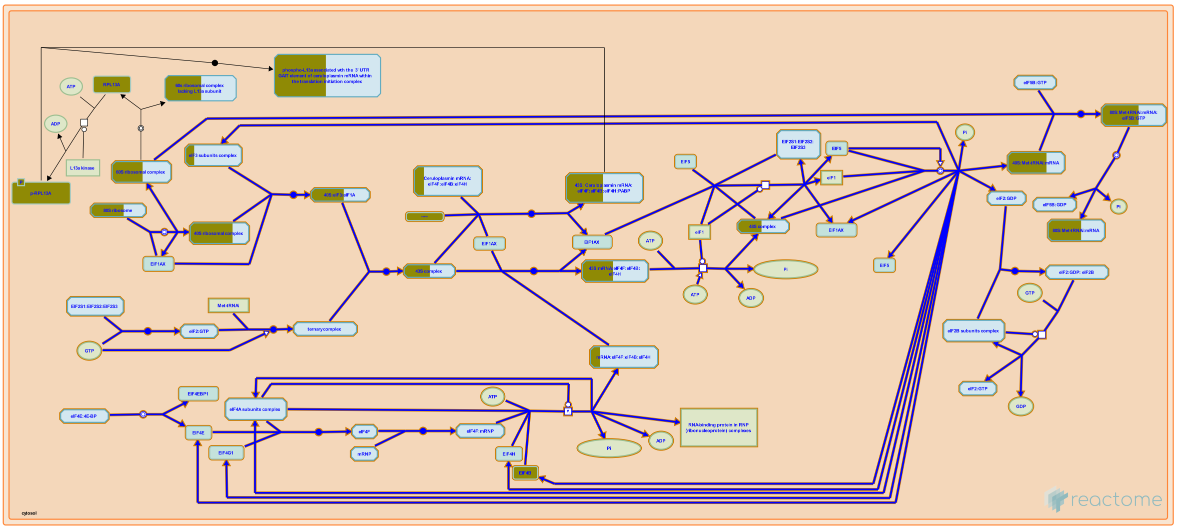


Translation initiation is a complex process in which the Met-tRNAi initiator, 40S, and 60S ribosomal subunits are assembled by eukaryotic initiation factors (eIFs) into an 80S ribosome at the start codon of an mRNA. The basic mechanism for this process can be described as a series of five steps: 1) formation of a pool of free 40S subunits, 2) formation of the ternary complex (Met-tRNAi/eIF2/GTP), and subsequently, the 43S complex (comprising the 40S subunit, Met-tRNAi/eIF2/GTP, eIF3 and eIF1A), 3) activation of the mRNA upon binding of the cap-binding complex eIF4F, and factors eIF4A, eIF4B and eIF4H, with subsequent binding to the 43S complex, 4) ribosomal scanning and start codon recognition, and 5) GTP hydrolysis and joining of the 60S ribosomal subunit.

##### References

##### Edit history

| Date | Action | Author |
|---|---|---|
| 2002-12-16 | Created | Merrick WC |
| 2023-08-26 | Modified | Wright A |

##### 56 submitted entities found in this pathway, mapping to 70 Reactome entities

| Input | UniProt Id | Input | UniProt Id | Input | UniProt Id |
|---|---|---|---|---|---|
| AGRN | P32969 | EIF3F | O00303 | EIF3L | Q9Y262 |
| EIF4B | P23588 | FAU | P62861 | PABPC1 | P11940 |
| RPL10 | P27635, Q96L21 | RPL10A | P61313, P62906 | RPL13 | P26373, P40429 |
| RPL13A | P40429, P61313 | RPL15 | P61313 | RPL18 | Q07020 |
| RPL21 | P46778 | RPL27A | P46776, P61353 | RPL29 | P47914 |
| RPL3 | P39023, Q92901 | RPL30 | P62888 | RPL31 | P62899 |
| RPL32 | P62888, P62910 | RPL34 | P49207, P62899 | RPL35 | P42766 |
| RPL35A | P18077 | RPL36 | Q9Y3U8 | RPL38 | P63173 |
| RPL3L | Q92901 | RPL4 | P36578 | RPL5 | P46777 |
| RPL7 | P18124 | RPL7A | P18124, P62424 | RPL9P8 | P32969 |
| RPLP0 | P05388 | RPLP2 | P05387 | RPS10 | P46783 |
| RPS11 | P62280 | RPS12 | P25398 | RPS13 | P62277 |
| RPS14 | P62263 | RPS15 | P62841 | RPS15A | P62244 |

| Input | UniProt Id | Input | UniProt Id | Input | UniProt Id |
|---|---|---|---|---|---|
| RPS16 | P62249 | RPS17 | P08708 | RPS18 | P62269 |
| RPS2 | P15880, P46782 | RPS23 | P62266 | RPS27 | P42677, Q71UM5 |
| RPS27A | P42677, P62979, P62987 | RPS29 | P62273 | RPS3 | P23396 |
| RPS3A | P61247 | RPS4X | P62701, Q8TD47 | RPS5 | P46782 |
| RPS6 | P62753 | RPS8 | P62241 | RPS9 | P46781 |
| RPSA | P08865 | SOS2 | P42766 | | |

#### 21. Immunoregulatory interactions between a Lymphoid and a non-Lymphoid cell (R-HSA-198933)

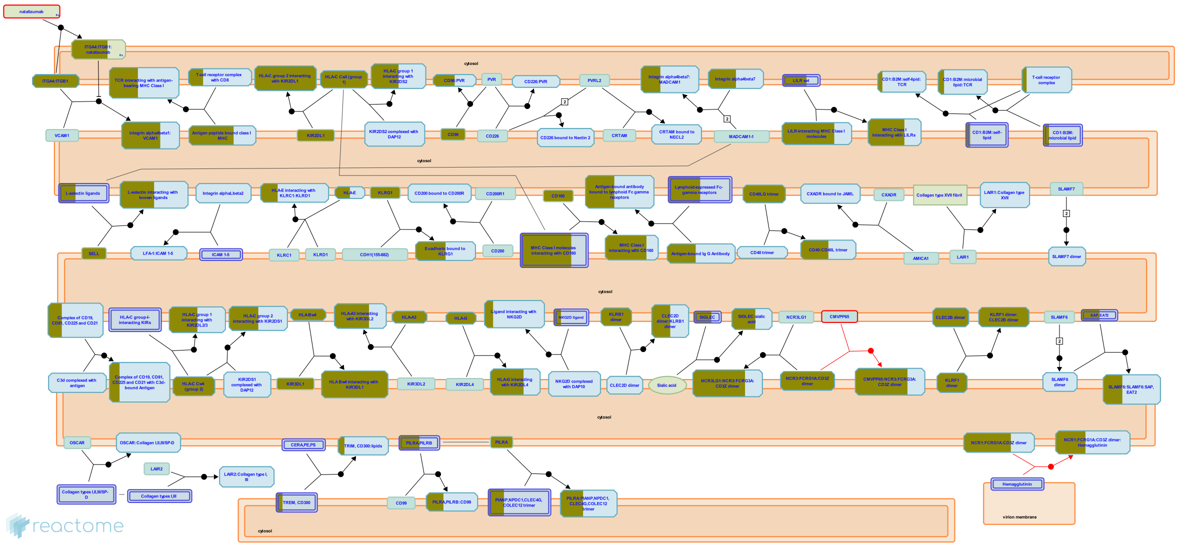

A number of receptors and cell adhesion molecules play a key role in modifying the response of cells of lymphoid origin (such as B-, T- and NK cells) to self and tumor antigens, as well as to pathogenic organisms.

Molecules such as KIRs and LILRs form part of a crucial surveillance system that looks out for any derangement, usually caused by cancer or viral infection, in MHC Class I presentation. Somatic cells are also able to report internal functional impairment by displaying surface stress markers such as MICA. The presence of these molecules on somatic cells is picked up by C-lectin NK immune receptors.

Lymphoid cells are able to regulate their location and movement in accordance to their state of activation, and home in on tissues expressing the appropriate complementary ligands. For example, lymphoid cells may fine tune the presence and concentration of adhesion molecules belonging to the IgSF, Selectin and Integrin class that interact with a number of vascular markers of inflammation.

Furthermore, there are a number of avenues through which lymphoid cells may interact with antigen. This may be presented directly to a specific T-cell receptor in the context of an MHC molecule. Antigen-antibody complexes may anchor to the cell via a small number of lymphoid-specific Fc receptors that may, in turn, influence cell function further. Activated complement factor C3d binds to both antigen and to cell surface receptor CD21. In such cases, the far-reaching influence of CD19 on B-lymphocyte function is tempered by its interaction with CD21.

##### References


Kelley J, Trowsdale J & Walter L (2005). Comparative genomics of natural killer cell receptor gene clusters. PLoS Genet., 1, 129-39.

Tomasello E, Walzer T, Vivier E, Baratin M & Ugolini S (2008). Functions of natural killer cells. Nat. Immunol., 9, 503-10.

Vivier E, Harris J, Trowsdale J, Vely F, Nedvetzki S, Davis DM, ... Pende D (2007). Reciprocal regulation of human natural killer cells and macrophages associated with distinct immune synapses. Blood, 109, 3776-85.

Batista FD & Carrasco YR (2006). B cell recognition of membrane-bound antigen: an exquisite way of sensing ligands. Curr Opin Immunol, 18, 286-91.

Shaw A & Cemerski S (2006). Immune synapses in T-cell activation. Curr Opin Immunol, 18, 298-304 .

#### Edit history

| Date | Action | Author |
|---|---|---|
| 2007-07-08 | Authored | de Bono B |
| 2007-07-08 | Created | de Bono B |
| 2007-08-06 | Reviewed | Trowsdale J |
| 2015-03-27 | Authored | Garapati P V |
| 2015-05-13 | Reviewed | Barrow AD |
| 2023-08-26 | Modified | Wright A |

#### 77 submitted entities found in this pathway, mapping to 86 Reactome entities

| Input | UniProt Id | Input | UniProt Id | Input | UniProt Id |
|---|---|---|---|---|---|
| B2M | P61769 | CD160 | O95971 | CD1C | P29017 |
| CD22 | P20273 | CD247 | P20963-1 | CD300E | Q496F6 |
| CD34 | P28906-1 | CD40LG | P29965 | CD8B | P10966 |
| CD96 | P40200 | CLEC2B | Q92478 | FCGR1A | P12314 |
| FCGR3A | P08637 | HLA-A | P04439 | HLA-B | P01889 |
| HLA-C | P10321 | HLA-G | P17693 | HLA-H | P01893 |
| IFITM1 | P13164 | IFITM3 | P13164 | IGHV1-69 | P01742 |
| IGHV3-11 | P01762 | IGHV3-23 | P01764 | IGHV3-30 | P01768 |
| IGHV3-33 | P01772 | IGHV4-34 | P06331 | IGHV4-39 | P01824 |
| IGHV4-59 | P01825 | IGKC | P01834 | IGKV1-12 | A0A0C4DH73 |
| IGKV1-17 | P01599 | IGKV1-33 | P01594 | IGKV1-5 | P01602 |
| IGKV1D-39 | P04432 | IGKV3-11 | P04433 | IGKV3-15 | P01624 |
| IGKV3-20 | P01619 | IGLC1 | P0CG04 | IGLC2 | P0DOY2 |
| IGLC3 | P0DOY3 | IGLC7 | A0M8Q6 | IGLV1-40 | P01703, Q5NV69 |
| IGLV1-44 | P01699, Q5NV81 | IGLV1-47 | P01700 | IGLV1-51 | P01701 |
| IGLV2-14 | P01704 | IGLV2-23 | P01705, Q5NV89 | IGLV3-1 | P01715 |
| IGLV3-19 | P01714 | IGLV3-21 | P80748 | IGLV3-25 | P01717, Q5NV90 |
| IGLV6-57 | P01721 | ITGA4 | P13612 | ITGB1 | P05556 |
| KIR2DL1 | P43626, P43629 | KIR3DL1 | P43629 | KLRB1 | Q12918 |
| KLRF1 | Q9NZS2 | KLRG1 | Q96E93 | LILRA4 | P59901 |
| LILRA5 | A6NI73, Q8N149 | LILRA6 | Q6PI73 | LILRB1 | Q8NHL6 |
| LILRB2 | Q8N423 | LILRB4 | Q8N149, Q8NHJ6 | MAL | P01732 |
| MICB | Q29980 | NCR3 | O14931 | NPDC1 | Q9NQX5 |
| PILRA | Q9UKJ1 | SELL | P14151 | SH2D1B | O14796 |
| SIGLEC1 | Q9BZZ2 | SIGLEC5 | O15389, O43699 | SIGLEC7 | Q9Y286 |
| SIGLEC8 | Q96PQ1, Q9NYZ4 | TREML1 | Q86YW5 | | |

#### 22. Regulation of expression of SLITs and ROBOs (R-HSA-9010553)

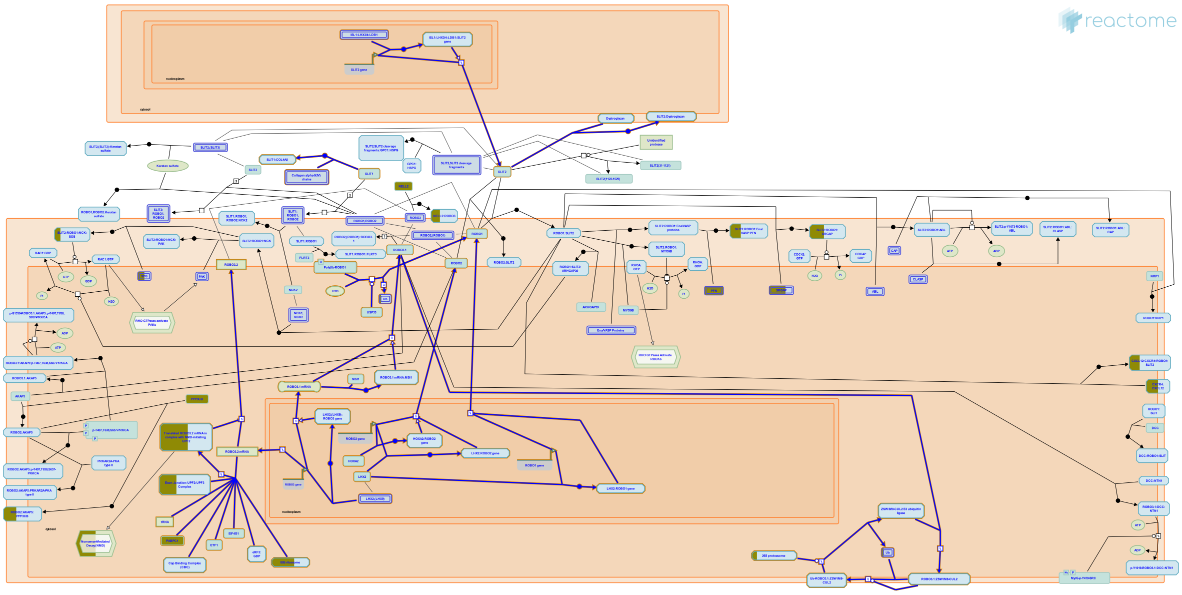

Expression of SLIT and ROBO proteins is regulated at the level of transcription, translation and protein localization and stability. LIM-homeodomain transcription factors LHX2, LHX3, LHX4, LHX9 and ISL1 have so far been implicated in a cell type-dependent transcriptional regulation of ROBO1, ROBO2, ROBO3 and SLIT2 (Wilson et al. 2008, Marcos-Mondejar et al. 2012, Kim et al. 2016). Homeobox transcription factor HOXA2 is involved in transcriptional regulation of ROBO2 (Geisen et al. 2008). Transcription of SLIT1 during optic tract development in Xenopus is stimulated by FGF signaling and may also involve the transcription factor HOXA2, but the mechanism has not been established (Atkinson-Leadbeater et al. 2010). PAX6 and the homeodomain transcription factor NKX2.2 are also implicated in regulation of SLIT1 transcription (Genethliou et al. 2009). An RNA binding protein, MSI1, binds ROBO3 mRNA and promotes its translation, thus increasing ROBO3 protein levels (Kuwako et al. 2010). A poorly studied E3 ubiquitin ligase ZSWIM8 promotes degradation of ROBO3 (Wang et al. 2013). ROBO1 is protein half-life is increased via deubiquitination of ROBO1 by a ubiquitin protease USP33 (Yuasa-Kawada et al. 2009, Huang et al. 2015). Interaction of SLIT2 with DAG1 (dystroglycan) is important for proper localization of SLIT2 at the floor plate (Wright et al. 2012). Interaction of SLIT1 with a type IV collagen COL4A5 is important for localization of SLIT1 to the basement membrane of the optical tectum (Xiao et al. 2011).

##### References


Rao Y, Kinoshita-Kawada M, Wu JY & Yuasa-Kawada J (2009). Deubiquitinating enzyme USP33/VDU1 is required for Slit signaling in inhibiting breast cancer cell migration. Proc. Natl. Acad. Sci. U.S.A., 106, 14530-5.

Ma L, Leung H, Lyon KA, Leahy DJ, Wright KM & Ginty DD (2012). Dystroglycan organizes axon guidance cue localization and axonal pathfinding. Neuron, 76, 931-44.

Brunet JF, Rijli FM, Pasqualetti M, Geisen MJ, Chédotal A, Di Meglio T & Ducret S (2008). Hox paralog group 2 genes control the migration of mouse pontine neurons through slit-robo signaling. PLoS Biol., 6, e142.

Dodd J, Shafer B, Lee KJ & Wilson SI (2008). A molecular program for contralateral trajectory: Rig-1 control by LIM homeodomain transcription factors. Neuron, 59, 413-24.

Zhu L, Wen P, Liu J, Kong R, Wu JY, Chen X, ... Quan C (2015). USP33 mediates Slit-Robo signaling in inhibiting colorectal cancer cell migration. Int. J. Cancer, 136, 1792-802.

#### Edit history

| Date | Action | Author |
|---|---|---|
| 2017-06-27 | Authored | Orlic-Milacic M |
| 2017-06-27 | Created | Orlic-Milacic M |
| 2017-07-31 | Reviewed | Jaworski A |
| 2017-08-04 | Edited | Orlic-Milacic M |
| 2023-08-26 | Modified | Wright A |

#### 61 submitted entities found in this pathway, mapping to 75 Reactome entities

| Input | UniProt Id | Input | UniProt Id | Input | UniProt Id |
|---|---|---|---|---|---|
| AGRN | P32969 | CASC3 | O15234 | FAU | P62861 |
| MAGOH | P61326 | PABPC1 | P11940 | PSMB8 | P28062 |
| PSMB9 | P28065 | PSME1 | Q06323 | PSME2 | Q9UL46 |
| RPL10 | P27635, Q96L21 | RPL10A | P61313, P62906 | RPL13 | P26373, P40429 |
| RPL13A | P40429, P61313 | RPL15 | P61313 | RPL18 | Q07020 |
| RPL21 | P46778 | RPL27A | P46776, P61353 | RPL29 | P47914 |
| RPL3 | P39023, Q92901 | RPL30 | P62888 | RPL31 | P62899 |
| RPL32 | P62888, P62910 | RPL34 | P49207, P62899 | RPL35 | P42766 |
| RPL35A | P18077 | RPL36 | Q9Y3U8 | RPL38 | P63173 |
| RPL3L | Q92901 | RPL4 | P36578 | RPL5 | P46777 |
| RPL7 | P18124 | RPL7A | P18124, P62424 | RPL9P8 | P32969 |
| RPLP0 | P05388 | RPLP2 | P05387 | RPN2 | Q99460 |
| RPS10 | P46783 | RPS11 | P62280 | RPS12 | P25398 |
| RPS13 | P62277 | RPS14 | P62263 | RPS15 | P62841 |
| RPS15A | P62244 | RPS16 | P62249 | RPS17 | P08708 |
| RPS18 | P62269 | RPS2 | P15880, P46782 | RPS23 | P62266 |
| RPS27 | P42677, Q71UM5 | RPS27A | P42677, P62979, P62987 | RPS29 | P62273 |
| RPS3 | P23396 | RPS3A | P61247 | RPS4X | P62701, Q8TD47 |
| RPS5 | P46782 | RPS6 | P62753 | RPS8 | P62241 |
| RPS9 | P46781 | RPSA | P08865 | SOS2 | P42766 |
| UPF3B | Q9BZI7 | | | | |

#### 23. Cellular response to starvation (R-HSA-9711097)

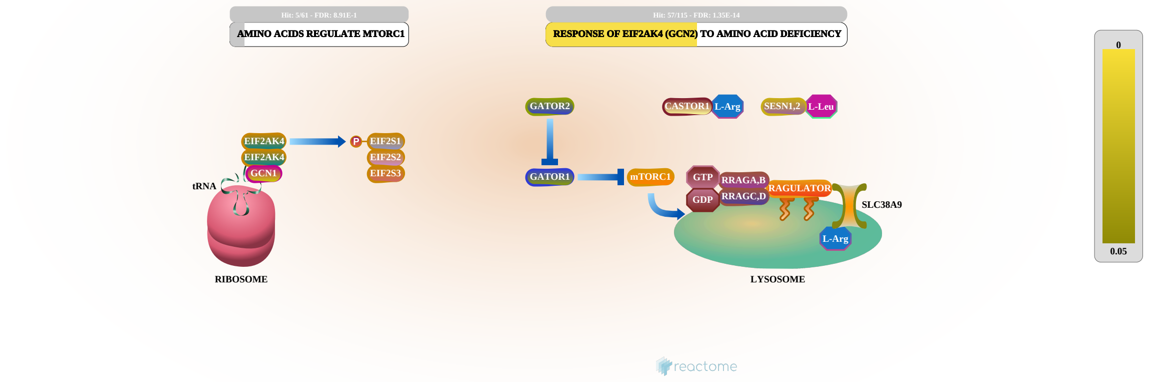


Deprivation of nutrients triggers diverse short- and long terms adaptations in cells. Here we have annotated two aspects of cellular responses to amino acid deprivation, ones mediated by EIF2AK4 and ones mediated by mTORC.

EIF2AK4 (GCN2) senses amino acid deficiency by binding uncharged tRNAs near the ribosome, and phosphorylating EIF2S1 (reviewed in Chaveroux et al. 2010, Castilho et al. 2014, Gallinetti et al. 2013, Bröer and Bröer 2017, Wek 2018). This reduces translation of most mRNAs but increases translation of mRNAs, notably ATF4, that mediate stress responses (reviewed in Kilberg et al. 2012, Wortel et al. 2017; Dever and Hinnebusch 2005).

The mTORC1 complex acts as an integrator that regulates translation, lipid synthesis, autophagy, and cell growth in response to multiple inputs, notably glucose, oxygen, amino acids, and growth factors such as insulin (reviewed in Sabatini 2017, Meng et al. 2018, Kim and Guan 2019).

MTOR, the kinase subunit of mTORC1, is activated by interaction with RHEB:GTP at the cytosolic face of lysosomal membrane (Long et al. 2005, Tee et al. 2005, Long et al. 2007, Yang et al. 2017). This process is regulated by various individual amino acids (reviewed in Zhuang et al. 2019, Wolfson and Sabatini 2017, Yao et al. 2017) and is reversed in response to the removal of amino acids, through the action of TSC1 (Demetriades et al. 2014).

##### References


Busch S, Lin Y, Long X, Avruch J & Ortiz-Vega S (2007). The Rheb switch 2 segment is critical for signaling to target of rapamycin complex 1. J. Biol. Chem., 282, 18542-51.

Demetriades C, Teleman AA & Doumpas N (2014). Regulation of TORC1 in response to amino acid starvation via lysosomal recruitment of TSC2. Cell, 156, 786-99.

Sabatini DM (2017). Twenty-five years of mTOR: Uncovering the link from nutrients to growth. Proc. Natl. Acad. Sci. U.S.A., 114, 11818-11825.

Sabatini DM & Wolfson RL (2017). The Dawn of the Age of Amino Acid Sensors for the mTORC1 Pathway. Cell Metab., 26, 301-309.

Yang A, Yang HJ, Jiang X, Li B, Pavletich NP, Yang H, ... Miller M (2017). Mechanisms of mTORC1 activation by RHEB and inhibition by PRAS40. Nature, 552, 368-373.

#### Edit history

| Date | Action | Author |
|---|---|---|
| 2020-11-19 | Authored | Stephan R |
| 2021-01-13 | Created | Stephan R |
| 2021-02-19 | Edited | D'Eustachio P |
| 2023-08-26 | Modified | Wright A |

#### 58 submitted entities found in this pathway, mapping to 73 Reactome entities

| Input | UniProt Id | Input | UniProt Id | Input | UniProt Id |
|---|---|---|---|---|---|
| AGRN | P32969 | ATF3 | P18847 | ATP6V0E2 | Q8NHE4 |
| FAU | P62861 | FLCN | Q8NFG4 | KPTN | Q9Y664 |
| LAMTOR1 | Q6IAA8 | RPL10 | P27635, Q96L21 | RPL10A | P61313, P62906 |
| RPL13 | P26373, P40429 | RPL13A | P40429, P61313 | RPL15 | P61313 |
| RPL18 | Q07020 | RPL21 | P46778 | RPL27A | P46776, P61353 |
| RPL29 | P47914 | RPL3 | P39023, Q92901 | RPL30 | P62888 |
| RPL31 | P62899 | RPL32 | P62888, P62910 | RPL34 | P49207, P62899 |
| RPL35 | P42766 | RPL35A | P18077 | RPL36 | Q9Y3U8 |
| RPL38 | P63173 | RPL3L | Q92901 | RPL4 | P36578 |
| RPL5 | P46777 | RPL7 | P18124 | RPL7A | P18124, P62424 |
| RPL9P8 | P32969 | RPLP0 | P05388 | RPLP2 | P05387 |
| RPS10 | P46783 | RPS11 | P62280 | RPS12 | P25398 |
| RPS13 | P62277 | RPS14 | P62263 | RPS15 | P62841 |
| RPS15A | P62244 | RPS16 | P62249 | RPS17 | P08708 |
| RPS18 | P62269 | RPS2 | P15880, P46782 | RPS23 | P62266 |
| RPS27 | P42677, Q71UM5 | RPS27A | P42677, P62979, P62987 | RPS29 | P62273 |
| RPS3 | P23396 | RPS3A | P61247 | RPS4X | P62701, Q8TD47 |
| RPS5 | P46782 | RPS6 | P62753 | RPS8 | P62241 |
| RPS9 | P46781 | RPSA | P08865 | SESN1 | Q9Y6P5 |
| SOS2 | P42766 | | | | |

| Input | Ensembl Id |
|---|---|
| ATF3 | ENSG00000162772 |

#### 24. SARS-CoV-1 modulates host translation machinery (R-HSA-9735869)

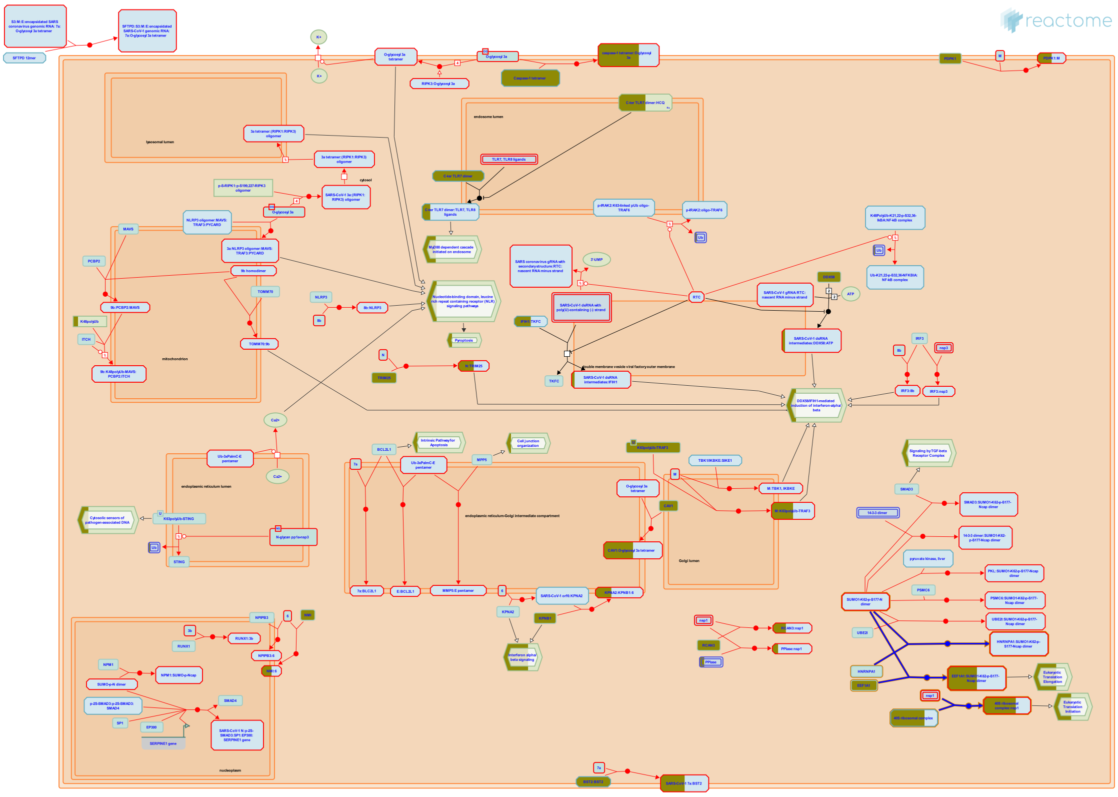

**Diseases:** severe acute respiratory syndrome.

Severe acute respiratory syndrome coronavirus type 1 (SARS-CoV-1) nonstructural protein 1 (nsp1) and nucleocapsid protein (N) disrupt mRNA translation upon SARS-CoV-1 infection in human cells.

##### References


Huang C, Makino S, Lokugamage KG & Narayanan K (2012). Severe acute respiratory syndrome coronavirus protein nsp1 is a novel eukaryotic translation inhibitor that represses multiple steps of translation initiation. J Virol, 86, 13598-608.

Cao C, Wang Q, Zhou B, Li X, Ma Q, Liu J, ... Liu X (2008). The nucleocapsid protein of severe acute respiratory syndrome coronavirus inhibits cell cytokinesis and proliferation by interacting with translation elongation factor 1alpha. J Virol, 82, 6962-71.


##### Edit history

| Date | Action | Author |
|---|---|---|
| 2020-06-25 | Authored | Shamovsky V |
| 2021-01-26 | Reviewed | D'Eustachio P |
| 2021-05-12 | Authored | Stephan R |
| 2021-07-05 | Created | Shamovsky V |
| 2022-08-11 | Edited | Shamovsky V |
| 2023-03-08 | Modified | Matthews L |

##### 25 submitted entities found in this pathway, mapping to 29 Reactome entities

| Input | UniProt Id | Input | UniProt Id | Input | UniProt Id |
|---|---|---|---|---|---|
| EEF1A1 | P68104 | FAU | P62861 | RPS10 | P46783 |
| RPS11 | P62280 | RPS12 | P25398 | RPS13 | P62277 |
| RPS14 | P62263 | RPS15 | P62841 | RPS15A | P62244 |
| RPS16 | P62249 | RPS17 | P08708 | RPS18 | P62269 |
| RPS2 | P15880, P46782 | RPS23 | P62266 | RPS27 | P42677, Q71UM5 |
| RPS27A | P42677, P62979 | RPS29 | P62273 | RPS3 | P23396 |
| RPS3A | P61247 | RPS4X | P62701, Q8TD47 | RPS5 | P46782 |
| RPS6 | P62753 | RPS8 | P62241 | RPS9 | P46781 |
| RPSA | P08865 | | | | |

#### 25. FCGR activation (R-HSA-2029481)

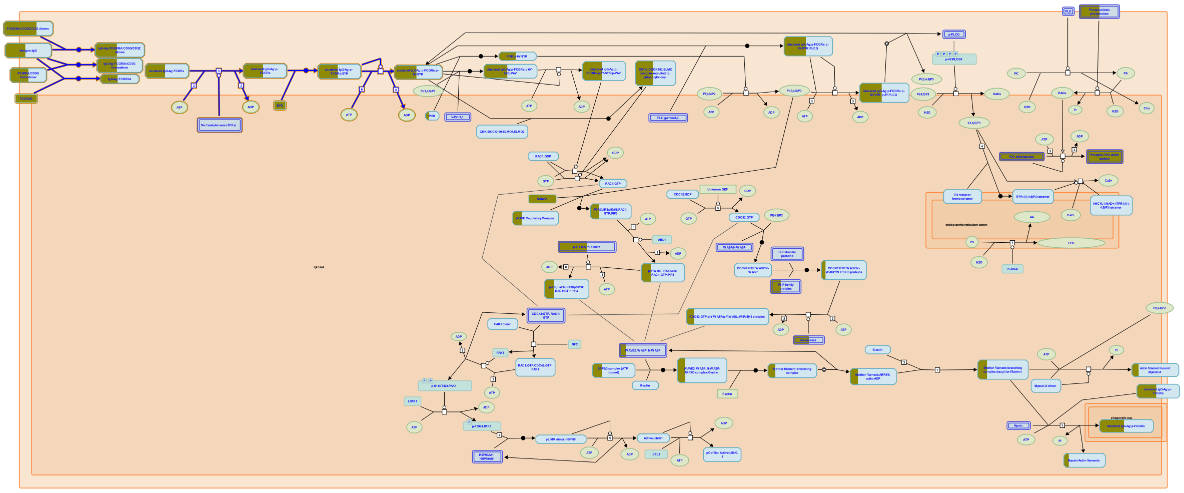

**Cellular compartments:** plasma membrane, cytosol.

Cross-linking of FCGRs with IgG coated immune complexes results in tyrosine phosphorylation of the immuno tyrosine activation motif (ITAMs) of the rececptor by membrane-bound tyrosine kinases of the SRC family. The phosphorylated ITAM tyrosines serve as docking sites for Src homology 2 (SH2) domain-containing SYK kinase. Recruitment and activation of SYK is critical for FCGR-mediated signaling in phagocytosis, but the exact role of SYK in this process is unclear. Activated SYK then transmits downstream signals leading to actin polymerization and particle internalization.

##### References


Rosales C & García-García E (2002). Signal transduction during Fc receptor-mediated phagocytosis. J Leukoc Biol, 72, 1092-108.

Tridandapani S, Joshi T & Butchar JP (2006). Fcgamma receptor signaling in phagocytes. Int J Hematol, 84, 210-6.

Dart AE, Caron E, Groves E & Covarelli V (2008). Molecular mechanisms of phagocytic uptake in mammalian cells. Cell Mol Life Sci, 65, 1957-76.


##### Edit history

| Date | Action | Author |
| --- | --- | --- |
| 2012-01-04 | Edited | Garapati P V |
| 2012-01-04 | Authored | Garapati P V |
| 2012-01-04 | Created | Garapati P V |
| 2012-05-15 | Reviewed | Rosales C |
| 2023-08-26 | Modified | Wright A |

##### 40 submitted entities found in this pathway, mapping to 45 Reactome entities

| Input | UniProt Id | Input | UniProt Id | Input | UniProt Id |
| --- | --- | --- | --- | --- | --- |
| CD247 | P20963-1 | FCGR1A | P12314 | FCGR2A | P12318 |

| Input | UniProt Id | Input | UniProt Id | Input | UniProt Id |
|---|---|---|---|---|---|
| FCGR3A | P08637 | IGHG1 | P01857, P01861 | IGHG3 | P01860 |
| IGHG4 | P01861 | IGHV1-69 | P01742 | IGHV3-11 | P01762 |
| IGHV3-23 | P01764 | IGHV3-30 | P01768 | IGHV3-33 | P01772 |
| IGHV4-34 | P06331 | IGHV4-39 | P01824 | IGHV4-59 | P01825 |
| IGKC | P01834 | IGKV1-12 | A0A0C4DH73 | IGKV1-17 | P01599 |
| IGKV1-33 | P01594 | IGKV1-5 | P01602 | IGKV1D-39 | P04432 |
| IGKV3-11 | P04433 | IGKV3-15 | P01624 | IGKV3-20 | P01619 |
| IGLC1 | P0CG04 | IGLC2 | P0DOY2 | IGLC3 | P0DOY3 |
| IGLC7 | A0M8Q6 | IGLV1-40 | P01703, Q5NV69 | IGLV1-44 | P01699, Q5NV81 |
| IGLV1-47 | P01700 | IGLV1-51 | P01701 | IGLV2-14 | P01704 |
| IGLV2-23 | P01705, Q5NV89 | IGLV3-1 | P01715 | IGLV3-19 | P01714 |
| IGLV3-21 | P80748 | IGLV3-25 | P01717, Q5NV90 | IGLV6-57 | P01721 |
| SYK | P43405 | | | | |

### 6. Identifiers found

Below is a list of the input identifiers that have been found or mapped to an equivalent element in Reactome, classified by resource.

##### 1084 of the submitted entities were found, mapping to 1396 Reactome entities

| Input | UniProt Id | Input | UniProt Id | Input | UniProt Id |
|---|---|---|---|---|---|
| AACS | Q86V21 | ABAT | P80404 | ABCA1 | O95477 |
| ABCB1 | P08183 | ABCB10 | Q9NRK6 | ABCG1 | P45844 |
| ABHD17A | Q96GS6 | ABHD4 | Q8TB40 | ABLIM1 | O14639 |
| ACOT8 | O14734 | ACOT9 | Q9Y305 | ACP6 | Q9NPH0 |
| ACSL1 | P33121 | ACTA2 | P62736 | ACTR3 | P61158 |
| ADA2 | Q9NZK5 | ADAM12 | O43184 | ADAM9 | Q13443 |
| ADAMTS1 | Q9UHI8 | ADAMTS10 | Q9H324 | ADAMTS2 | O95450 |
| ADAMTSL4 | Q6UY14 | ADAR | P55265 | ADCY4 | Q8NFM4 |
| ADGRE3 | Q9BY15 | ADGRG1 | Q86Y34 | ADM | P35318 |
| ADRA1D | P25100 | ADRA2B | P18089 | AGRN | P32969 |
| AIM2 | O14862 | AK4 | P27144 | AK5 | Q9Y6K8 |
| AKAP1 | Q92667 | AKR1B1 | P15121 | AKR1C3 | P42330 |
| ALDOA | P04075 | ALDOC | P09972 | ALKBH2 | Q6NS38 |
| ALOX15 | P16050 | ALPL | P05186 | ALS2CL | Q60I27 |
| AMY1A | P0DUB6 | AMY2A | P04746, P19961 | AMY2B | P19961 |
| ANKFY1 | Q9P2R3 | ANKH | Q9HCJ1 | ANPEP | P15144 |
| AP1B1 | Q10567 | APBA2 | Q99767 | APBB1 | O00213 |
| APOBEC3A | P31941 | APOBEC3B | Q9UH17 | APOBEC3G | Q9HC16-1, Q9HC16-3 |
| APOBEC3H | Q6NTF7 | APOD | P05090 | APOL1 | O14791 |
| ARF1 | P84077 | ARF4 | P18085 | ARL2 | P36404 |
| ARL3 | Q13795 | ARPC1B | O15143 | ARRB1 | P49407 |
| ASCC2 | Q9H1I8 | ASIC1 | P78348 | ATAD2 | Q6PL18 |
| ATF3 | P18847 | ATG4B | Q96DT6, Q9Y4P1 | ATG9B | Q674R7 |
| ATP1A1 | P05023 | ATP1B1 | P05026 | ATP1B3 | P54709 |
| ATP2C1 | P98194 | ATP5MC2 | Q06055 | ATP5PD | O75947 |
| ATP6V0E2 | Q8NHE4 | AURKA | O14965 | AUTS2 | Q8WXX7 |
| AXIN2 | Q9Y2T1 | B2M | P61769 | B3GALT2 | O43825 |
| BABAM1 | Q9NWV8 | BAG3 | O95817 | BAG6 | P46379 |
| BAIAP2 | Q9UQB8 | BANP | Q8N9N5 | BBS7 | Q8IWZ6 |
| BCL2A1 | Q16548 | BCL2L14 | Q9BZR8 | BCL9 | O00512 |
| BCR | P11274 | BIRC2 | Q13489, Q13490 | BLVRA | P53004 |
| BMPR2 | Q13873 | BRCA1 | P38398 | BRCA2 | P51587 |
| BRI3 | O95415 | BST1 | Q10588 | BST2 | Q10589 |
| BTF3 | O00478, P78410 | BTK | Q06187 | BTN2A2 | Q8WVV5, Q9UIR0 |
| BTN3A1 | O00481 | BTNL8 | Q6UX41 | BUB1 | O43683, O60566 |
| C1GALT1 | Q9NS00 | C1QA | P02745 | C1QB | P02746 |
| C1QC | P02747 | C2 | P06681 | C3AR1 | Q16581 |
| C5 | P01031 | C5AR2 | Q9P296 | CA6 | P23280 |
| CACNA1A | O00555 | CACNA1E | Q15878 | CACNA1I | Q9P0X4 |
| CAMK4 | Q16566 | CAMKMT | Q7Z624 | CAMLG | P49069 |
| CAPN3 | P20807 | CAPN5 | O15484 | CAPNS1 | P04632, Q96L46 |

| Input | UniProt Id | Input | UniProt Id | Input | UniProt Id |
|---|---|---|---|---|---|
| CARNS1 | A5YM72 | CASC3 | O15234 | CASP1 | P29466 |
| CASP4 | P49662 | CASP5 | P51878 | CASP7 | P55210 |
| CAT | P04040 | CAV1 | Q03135 | CCL2 | P13500 |
| CCL20 | P78556 | CCL23 | P55773-2 | CCL8 | P80098 |
| CCNA1 | P78396 | CCNA2 | P20248 | CCR1 | P32246 |
| CCR6 | P51684 | CCR7 | P32248 | CCRL2 | O00421 |
| CD160 | O95971 | CD163 | Q86VB7 | CD1C | P29017 |
| CD22 | P20273 | CD24 | P25063 | CD247 | P20963-1 |
| CD27 | P26842 | CD274 | Q9NZQ7 | CD300E | Q496F6 |
| CD34 | P28906-1 | CD38 | P28907 | CD40LG | P29965 |
| CD68 | P34810 | CD8B | P10966 | CD96 | P40200 |
| CDC14B | O60729 | CDC42EP2 | O14613 | CDC45 | O75419 |
| CDKN1A | P38936 | CDKN1C | P49918 | CEACAM1 | P13688 |
| CEP164 | Q9UPV0 | CEP41 | Q9BYV8 | CEP55 | Q92753 |
| CEP57 | Q86XR8 | CEP83 | Q9Y592 | CETP | P11597 |
| CFB | P00751 | CFP | P27918 | CHM | P24386 |
| CHMP2A | O43633 | CHMP5 | Q9NZZ3 | CHN1 | P15882 |
| CHTF8 | P0CG13 | CIR1 | P38117 | CISH | Q9NSE2 |
| CKAP4 | Q07065 | CLDN23 | Q96B33 | CLDN5 | O00501 |
| CLEC10A | Q6UW15, Q8IUN9 | CLEC12A | Q5QGZ9 | CLEC2B | Q92478 |
| CLEC4D | Q8WXI8 | CLEC4E | Q9UJ71, Q9ULY5 | CLEC7A | Q9BXN2 |
| CLOCK | O15516 | CLTC | Q00610, Q00610-1 | CNBP | P62633 |
| CNOT11 | Q9UKZ1 | CNTRL | Q7Z7A1-3 | COA6 | Q5JTJ3 |
| COG3 | Q96JB2 | COL13A1 | Q5TAT6 | COL16A1 | Q07092 |
| COL4A3 | Q01955 | COL5A3 | P25940 | COL6A2 | P12110 |
| COPE | O14579 | CPNE8 | Q86YQ8 | CPSF6 | Q16630 |
| CRAT | P43155 | CRTAP | O75718 | CSAD | Q9Y600 |
| CSTF3 | Q12996 | CTNNA1 | P35221 | CTNNBIP1 | Q9NSA3 |
| CTSF | Q9UBX1 | CTSL | O60911, P07711 | CUL9 | Q8IWT3 |
| CX3CR1 | P49238 | CXCL10 | P02778 | CXCL11 | O14625 |
| CXCL5 | P42830, P80162 | CXCL9 | Q07325 | CXCR2 | P25024, P25025 |
| CXCR4 | P61073 | CXCR6 | O00574 | CYBB | P04839 |
| CYFIP2 | Q96F07 | CYP2E1 | P05181 | CYP2S1 | Q96SQ9 |
| CYP4F22 | Q6NT55 | CYRIB | Q9NUQ9 | CYSLTR1 | Q9Y271 |
| CYTH3 | O43739 | DAPP1 | Q9UN19 | DBNL | Q9UJU6 |
| DBP | Q03518 | DCAF6 | Q58WW2 | DCLRE1A | Q6PJP8 |
| DDX17 | Q92841 | DDX58 | O95786 | DENND6A | Q8IWF6 |
| DHCR7 | Q9UBM7 | DHDDS | Q86SQ9 | DHRS3 | O75911 |
| DHRS9 | Q9BPW9 | DHX58 | Q96C10 | DIABLO | Q9NR28 |
| DIAPH2 | O60879-2, O60879-3 | DICER1 | Q9UPY3 | DLG5 | Q8TDM6 |
| DLGAP2 | Q9P1A6 | DNASE1L3 | P49184 | DOCK4 | Q8N1I0 |
| DOCK8 | Q8NF50 | DOLPP1 | Q86YN1 | DPYD | Q12882 |
| DPYSL2 | Q16555 | DRAP1 | Q14919 | DSC1 | Q08554 |
| DSEL | Q8IZU8 | DTX3L | Q8TDB6 | DUSP5 | Q16690 |
| DUSP6 | Q16828 | DUT | Q6SW70 | DYNC2H1 | Q8NCM8 |
| DYNLT1 | P63172 | E2F2 | Q14209 | ECT2 | Q9H8V3 |
| EDAR | Q9UNE0 | EEF1A1 | P68104 | EEF1A1P5 | Q5VTE0 |
| EEF1B2 | P24534 | EEF1G | P26641 | EEF2 | P13639 |
| EEPD1 | Q7L9B9 | EGLN1 | Q9GZT9 | EGR1 | P18146 |

| Input | UniProt Id | Input | UniProt Id | Input | UniProt Id |
|---|---|---|---|---|---|
| EIF2AK2 | P19525 | EIF3F | O00303 | EIF3L | Q9Y262 |
| EIF4B | P23588 | ELOVL4 | Q9GZR5 | ELOVL5 | Q9NYP7 |
| ELP5 | Q8TE02 | ELP6 | Q0PNE2 | EMD | P50402 |
| ENSA | O43768 | EPB41L3 | Q9Y2J2 | EPHA4 | P54764 |
| EPHB2 | P29323 | EPHX2 | P34913 | EPSTI1 | Q96J88 |
| ERF | P50548 | ESAM | Q96AP7 | ESYT2 | A0FGR8, A0FGR9 |
| ETNK1 | Q9HBU6 | ETV6 | P41212 | EXOC1 | Q9NV70 |
| EXTL3 | O43909 | EZH2 | Q15910 | F11R | Q9Y624 |
| FAAP20 | Q6NZ36 | FAM13A | O94988 | FAM20A | Q96MK3 |
| FANCA | O15360 | FANCD2 | Q9BXW9 | FANCL | Q9NW38 |
| FASTK | Q14296-4 | FAU | P62861 | FBL | P22087 |
| FBLN2 | P98095 | FBXL8 | Q96CD0 | FBXO30 | Q8TB52 |
| FBXO6 | Q9NRD1 | FBXO7 | Q9Y3I1 | FBXO9 | Q9UK97 |
| FBXW11 | Q9UKB1 | FCAR | P24071 | FCER1A | P12319 |
| FCGR1A | P12314 | FCGR1B | Q92637 | FCGR2A | P12318 |
| FCGR3A | P08637 | FCGR3B | O75015 | FCN1 | O00602 |
| FFAR2 | O15552 | FFAR3 | O14843 | FGD2 | Q7Z6J4 |
| FGD4 | Q96M96 | FGF9 | P31371 | FGL2 | Q14314 |
| FLCN | Q8NFG4 | FLI1 | Q01543 | FLOT1 | O75955 |
| FLT3LG | P49771 | FLT4 | P35916 | FMNL2 | Q96PY5 |
| FTH1 | P02794 | FXN | Q16595 | GADD45B | P41440 |
| GALNT12 | Q8IXK2 | GAMT | Q14353 | GAPDH | P04406 |
| GBP1 | P32455 | GBP2 | P32456 | GBP3 | Q9H0R5 |
| GBP4 | Q96PP9 | GBP5 | Q96PP8 | GBP6 | Q6ZN66 |
| GBX2 | P52951 | GCH1 | P30793 | GCHFR | P30047 |
| GCLC | P48506 | GCNT1 | Q02742 | GDI2 | P50395 |
| GET4 | Q7L5D6 | GFPT1 | Q06210 | GHRL | Q9UBU3-1, Q9UBU3-2 |
| GJC2 | Q5T442 | GLRX | P35754 | GLUL | P15104 |
| GM2A | P17900 | GNAQ | P50148 | GNB4 | Q9HAV0 |
| GNG7 | O60262 | GNLY | P22749 | GOLGA7 | Q7Z5G4 |
| GORASP1 | Q9BQQ3 | GPBAR1 | Q8TDU6 | GPC6 | Q9Y625 |
| GPD2 | P43304 | GPR183 | P32249 | GPR65 | Q8IYL9 |
| GPR68 | Q15743 | GPR84 | Q9NQS5 | GRAP | Q13588 |
| GRB14 | Q14449 | GRIN3A | Q8TCU5 | GRN | P28799 |
| GSTM2 | P28161 | GTPBP1 | Q9BX10 | GTPBP2 | Q9BX10 |
| GYG1 | P46976 | GZMM | P51124 | H1-2 | P16403 |
| H2BC12 | O60814, Q99877 | H2BC18 | Q5QNW6 | H2BC4 | P62807, Q93079 |
| H2BC5 | P58876 | H3-3A | P84243 | H3C4 | P68431, Q71DI3 |
| H4C8 | P62805 | HADHA | P40939 | HAS3 | O00219 |
| HBEGF | Q99075 | HBG1 | P69891 | HBG2 | P69892 |
| HDAC5 | Q9UQL6 | HELZ2 | Q9BYK8 | HERC4 | Q15034, Q5GLZ8 |
| HERC5 | Q9UII4 | HERC6 | Q8IVU3 | HERPUD1 | Q15011 |
| HGD | Q93099 | HK3 | P52790 | HLA-A | P04439 |
| HLA-B | P01889 | HLA-C | P10321 | HLA-DQA1 | P01909, P20036 |
| HLA-DQA2 | P01906 | HLA-DRA | P01903 | HLA-DRB4 | P13762 |
| HLA-G | P17693 | HLA-H | P01893 | HMBS | P08397 |
| HMGN2 | P05114 | HNMT | P50135 | HNRNPA2B1 | P22626 |
| HOMER2 | Q9NSB8 | HP | P00738 | HPS4 | Q9NQG7 |
| HSD17B13 | Q7Z5P4 | HSH2D | Q9NP31 | HSPA1A | P0DMV8 |

| Input | UniProt Id | Input | UniProt Id | Input | UniProt Id |
|---|---|---|---|---|---|
| HSPA6 | P17066 | HSPA7 | P48741 | ICE1 | Q9Y2F5 |
| ICOSLG | O75144 | ID2 | Q02363 | ID3 | Q02535 |
| IDO1 | P14902, Q6ZQW0 | IFI16 | Q16666 | IFI27 | P40305 |
| IFI30 | P13284 | IFI35 | P80217 | IFI6 | P09912 |
| IFIH1 | Q9BYX4 | IFIT1 | P09914 | IFIT2 | P09913 |
| IFIT3 | O14879 | IFIT5 | Q13325 | IFITM1 | P13164 |
| IFITM2 | Q01629 | IFITM3 | P13164, Q01628 | IFNG | P01579 |
| IFT122 | Q9HBG6 | IFT140 | Q96RY7 | IGF1R | P08069 |
| IGF2BP3 | O00425 | IGHA1 | P01876 | IGHE | P01854 |
| IGHG1 | P01857, P01861 | IGHG3 | P01860 | IGHG4 | P01861 |
| IGHV1-69 | P01742 | IGHV3-11 | P01762 | IGHV3-23 | P01764 |
| IGHV3-30 | P01768 | IGHV3-33 | P01772 | IGHV4-34 | P06331 |
| IGHV4-39 | P01824 | IGHV4-59 | P01825 | IGKC | P01834 |
| IGKV1-12 | A0A0C4DH73 | IGKV1-17 | P01599 | IGKV1-33 | P01594 |
| IGKV1-5 | P01602 | IGKV1D-39 | P04432 | IGKV3-11 | P04433 |
| IGKV3-15 | P01624 | IGKV3-20 | P01619 | IGLC1 | P0CG04 |
| IGLC2 | P0DOY2 | IGLC3 | P0DOY3 | IGLC7 | A0M8Q6 |
| IGLV1-40 | P01703, Q5NV69 | IGLV1-44 | P01699, Q5NV81 | IGLV1-47 | P01700 |
| IGLV1-51 | P01701 | IGLV2-14 | P01704 | IGLV2-23 | P01705, Q5NV89 |
| IGLV3-1 | P01715 | IGLV3-19 | P01714 | IGLV3-21 | P80748 |
| IGLV3-25 | P01717, Q5NV90 | IGLV6-57 | P01721 | IKBIP | Q70UQ0 |
| IL11RA | P08887, Q14626 | IL15 | P40933 | IL1RN | P18510 |
| IL27 | Q8NEV9 | IL2RB | P14784 | IL37 | Q9NZH6 |
| IL4R | P24394, Q01113 | IL5RA | P15509, Q01344 | IL6 | P05231 |
| IL6R | P08887, P08887-2 | IL7R | P16871 | IMPA2 | O14732 |
| IMPDH1 | P20839 | ING5 | Q8WYH8 | INPP5E | Q10713 |
| INTS7 | Q9NVH2 | IP6K1 | Q92551 | IP6K2 | Q9UHH9 |
| IQGAP1 | P46940 | IRF1 | P10914 | IRF5 | Q13568 |
| IRF7 | Q92985 | IRF9 | Q00978 | ISG15 | P05161 |
| ISG20 | Q96AZ6 | ITGA4 | P13612 | ITGA6 | P23229 |
| ITGB1 | P05556 | ITGB4 | P16144 | JADE1 | Q6IE81 |
| JAK2 | O60674 | JUP | P14923 | KAT8 | Q9H7Z6 |
| KCNH3 | Q9ULD8 | KCNJ15 | Q99712 | KCNQ3 | O43525 |
| KDM1B | Q8NB78 | KDM4B | O94953 | KHK | P50053 |
| KIR2DL1 | P43626, P43629 | KIR3DL1 | P43629 | KISS1R | Q969F8 |
| KIT | P10721 | KLF4 | O43474 | KLHL3 | Q9UH77 |
| KLRB1 | Q12918 | KLRF1 | Q9NZS2 | KLRG1 | Q96E93 |
| KMO | O15229 | KMT5B | Q4FZB7, Q86Y97 | KPNB1 | Q14974 |
| KPTN | Q9Y664 | KRT1 | P04264 | KRT5 | P13647 |
| KRT72 | Q14CN4 | KRT73 | Q86Y46 | KYNU | Q16719 |
| LAG3 | P18627 | LAMP3 | P34810 | LAMTOR1 | Q6IAA8 |
| LAP3 | Q13867 | LDLRAP1 | Q5SW96 | LEF1 | Q9UJU2 |
| LGALS3BP | Q08380 | LGALS9 | O00182 | LGMN | Q99538 |
| LHFPL2 | Q6ZUX7 | LIF | P15018 | LILRA4 | P59901 |
| LILRA5 | A6NI73, Q8N149 | LILRA6 | Q6PI73 | LILRB1 | Q8NHL6 |
| LILRB2 | Q8N423 | LILRB4 | Q8N149, Q8NHJ6 | LIPN | Q5VXI9 |
| LMNB1 | P20700 | LMO2 | P25791 | LPAR6 | P43657 |
| LRFN3 | Q9BTN0 | LRFN4 | Q6PJG9 | LRRK2 | Q5S007 |
| LRTOMT | Q8WZ04 | LTB | Q06643 | LTBP3 | Q14767, Q9NS15 |

| Input | UniProt Id | Input | UniProt Id | Input | UniProt Id |
|---|---|---|---|---|---|
| LY6E | Q16553 | LY96 | Q9Y6Y9 | LYPLA2 | O95372 |
| MAD1L1 | Q9Y6D9 | MAF | O75444 | MAFB | Q9Y5Q3 |
| MAGED1 | Q9Y5V3 | MAGED2 | Q9UNF1 | MAGOH | P61326 |
| MAL | P01732, P58753 | MAP2K3 | P46734 | MAP2K6 | P52564 |
| MAP3K1 | Q13233 | MAPK3 | P27361 | MARCKS | P29966 |
| MASTL | Q96GX5 | MATK | P42679 | MAX | P61244 |
| MBP | P13727 | MCAT | Q8IVS2 | MCM8 | Q9UJA3 |
| MCOLN3 | Q8TDD5 | MCRS1 | Q96EZ8 | MED16 | Q9Y2X0 |
| MED25 | Q71SY5, Q9NWA0 | MEF2C | Q06413 | MEFV | O15553 |
| MERTK | Q12866 | METTL1 | Q9UBP6 | METTL21A | Q8WXB1 |
| MFGE8 | Q08431 | MICAL1 | Q8TDZ2 | MICB | Q29980 |
| MIS18BP1 | Q6P0N0 | MLKL | Q8NB16 | MMP8 | P22894 |
| MNDA | P41218 | MOV10 | Q9HCE1 | MPST | P25325 |
| MRPL24 | Q96A35 | MRPL45 | Q9BRJ2 | MSR1 | P21757 |
| MSRA | Q9UJ68 | MT1A | P02795 | MT1E | P04732 |
| MT1F | P04733 | MT2A | P02795 | MTHFD2 | P13995 |
| MTMR12 | Q9C0I1 | MTX1 | Q13505 | MUC1 | P15941 |
| MUC6 | Q6W4X9 | MX1 | P20591 | MX2 | P20592 |
| MYBL1 | P10243 | MYBL2 | P10244 | MYD88 | Q99836 |
| NADK | O95544 | NAGK | Q9UJ70 | NAMPT | P43490 |
| NAPA | Q9Y2A7 | NBN | O60934 | NCALD | P61601 |
| NCAM1 | P13591 | NCAPG | Q9BPX3 | NCF1 | P14598 |
| NCF2 | P19878 | NCOA1 | Q15788 | NCOA3 | Q9Y6Q9 |
| NCR3 | O14931 | NDC80 | O14777 | NDEL1 | Q9GZM8 |
| NDUFA10 | O95299 | NDUFA3 | O95167 | NDUFAF1 | Q9Y375 |
| NDUFAF5 | Q5TEU4 | NDUFAF6 | Q330K2 | NDUFAF7 | Q7L592 |
| NDUFB4 | O95168 | NELL2 | Q99435 | NEO1 | Q92859 |
| NF1 | P21359 | NFIL3 | Q16649 | NHLRC3 | Q5JS37 |
| NLRC5 | Q86WI3 | NLRX1 | Q86UT6 | NMB | Q14956 |
| NMI | Q13287 | NMNAT3 | Q96T66 | NMUR1 | O94822 |
| NNMT | P40261 | NOC4L | Q9BVI4 | NOD2 | Q9HC29 |
| NOG | Q13253 | NOSIP | Q9Y314 | NPAS2 | Q99743 |
| NPC2 | P61916 | NPDC1 | Q9NQX5 | NR1D1 | P20393 |
| NR3C2 | P08235 | NRCAM | Q92823 | NRIP1 | P48552 |
| NSD3 | Q9BZ95 | NT5C3A | Q9H0P0-4 | NT5E | P21589 |
| NTNG2 | Q96CW9 | NUB1 | Q9Y5A7-1, Q9Y5A7-2 | NUDT5 | Q9UKK9 |
| NUP188 | Q5SRE5 | OAS1 | P00973 | OAS2 | P29728 |
| OAS3 | Q9Y6K5 | OASL | Q15646 | OAZ1 | P54368 |
| OGG1 | O15527, O15527-4 | OR2AG1 | Q9H205 | OSBPL6 | Q9BZF3 |
| OTOF | Q9HC10 | P2RX1 | P51575 | P2RY13 | Q9BPV8 |
| P3H2 | Q8IVL5 | PABPC1 | P11940 | PACSIN1 | Q9BY11 |
| PAFAH1B3 | Q15102 | PANX2 | Q96RD6 | PARD6A | Q9NPB6 |
| PARP10 | Q53GL7 | PARP14 | Q460N5 | PARP16 | Q8N5Y8 |
| PARP9 | Q8IXQ6 | PATL1 | Q86TB9 | PCGF5 | Q86SE9 |
| PCK2 | Q16822 | PCSK5 | Q92824 | PDCD1LG2 | Q9BQ51 |
| PDGFD | Q9GZP0 | PDGFRB | P09619 | PDK4 | Q16654 |
| PDLIM7 | Q9NR12 | PDPK1 | O15530 | PDSS1 | Q5T2R2 |
| PEX16 | Q9Y5Y5 | PFKP | Q01813 | PFN2 | P07737, P35080 |
| PGAM1 | P18669 | PGAP1 | Q75T13 | PHC3 | Q8NDX5 |

| Input | UniProt Id | Input | UniProt Id | Input | UniProt Id |
|---|---|---|---|---|---|
| PHF1 | O43189 | PHF19 | Q5T6S3 | PHF20L1 | A8MW92 |
| PHGDH | O43175 | PHYH | O14832 | PI3 | P19957 |
| PI4K2B | Q8TCG2 | PIGP | P57054 | PIGQ | Q9BRB3-2 |
| PIK3AP1 | Q6ZUJ8 | PIK3CA | P42336 | PILRA | Q9UKJ1 |
| PIN1 | Q13526 | PIWIL4 | Q7Z3Z4 | PJA1 | Q8NG27 |
| PKD2 | Q13563 | PLAAT4 | P53816, Q9UL19 | PLAUR | Q03405 |
| PLEKHA1 | Q9HB21 | PLEKHG1 | Q9ULL1 | PLOD1 | Q02809 |
| PLPP5 | Q8NEB5 | PML | P29590 | PMM2 | O15305 |
| PNPT1 | Q8TCS8 | POLQ | O75417 | POLR1E | O15160 |
| POLR3H | Q9Y535 | POTEKP | Q9BYX7 | PPARGC1B | Q86YN6 |
| PPBP | P02775 | PPIA | P62937 | PPP2R2D | Q66LE6 |
| PPP2R5D | Q14738 | PPP3CA | Q08209 | PPP3CB | P16298 |
| PPP3CC | P16298 | PPP3R1 | P63098 | PRAG1 | Q86YV5 |
| PRC1 | O43663 | PRKCB | P05771 | PRKCD | Q05655 |
| PRKCE | Q02156 | PRKCSH | P14314 | PRKDC | P78527 |
| PRPF4B | Q13523 | PRR5 | P85299 | PSMB8 | P28062 |
| PSMB9 | P28065 | PSME1 | Q06323 | PSME2 | Q9UL46 |
| PTCH1 | Q13635 | PTGDR | Q13258 | PTGDS | P41222 |
| PTGES3 | Q15185 | PTGFR | P43088 | PTK2B | Q14289 |
| PTPN2 | P17706-2 | PTPN22 | Q9Y2R2 | PTPRF | P10586 |
| PTPRN2 | Q92932 | PTPRO | Q92729 | PTPRS | P10586, Q13332 |
| QTRT1 | Q9BXR0 | RAB24 | Q969Q5 | RAB33A | Q14088 |
| RAB3D | O95716 | RAB43 | Q86YS6 | RAD21 | O60216 |
| RAD51AP1 | Q96B01 | RASGRF2 | O14827 | RBBP8 | Q99708 |
| RBCK1 | Q9BYM8 | RCAN3 | Q9UKA8 | RDX | P35241 |
| REC8 | O95072 | REG4 | Q06141 | REL | Q01201 |
| RGCC | Q9H4X1 | RGL1 | Q9NZL6 | RHBDF2 | Q6PJF5 |
| RHOBTB3 | O94955 | RHOT1 | Q8IXI2 | RIGI | O95786 |
| RIN2 | Q8WYP3 | RLIM | Q9NVW2 | RMI2 | Q96E14 |
| RNASE2 | P10153 | RNF213 | Q63HN8 | RNF31 | Q96EP0 |
| RNF38 | Q9P000 | RORA | P35398 | RORC | P51449 |
| RPA1 | P27694 | RPL10 | P27635, Q96L21 | RPL10A | P61313, P62906 |
| RPL13 | P26373, P40429 | RPL13A | P40429, P61313 | RPL15 | P61313 |
| RPL18 | Q07020 | RPL21 | P46778 | RPL27A | P46776, P61353 |
| RPL29 | P47914 | RPL3 | P39023, Q92901 | RPL30 | P62888 |
| RPL31 | P62899 | RPL32 | P62888, P62910 | RPL34 | P49207, P62899 |
| RPL35 | P42766 | RPL35A | P18077 | RPL36 | Q9Y3U8 |
| RPL38 | P63173 | RPL3L | Q92901 | RPL4 | P36578 |
| RPL5 | P46777 | RPL7 | P18124 | RPL7A | P18124, P62424 |
| RPL9P8 | P32969 | RPLP0 | P05388 | RPLP2 | P05387 |
| RPN2 | P04844 | RPS10 | P46783 | RPS11 | P62280 |
| RPS12 | P25398 | RPS13 | P62277 | RPS14 | P62263 |
| RPS15 | P62841 | RPS15A | P62244 | RPS16 | P62249 |
| RPS17 | P08708 | RPS18 | P62269 | RPS2 | P15880, P46782 |
| RPS23 | P62266 | RPS27 | P42677, Q71UM5 | RPS27A | P42677, P62979, P62987 |
| RPS29 | P62273 | RPS3 | P23396 | RPS3A | P61247 |
| RPS4X | P62701, Q8TD47 | RPS5 | P46782 | RPS6 | P62753 |
| RPS8 | P62241 | RPS9 | P46781 | RPSA | P08865 |
| RRAS | P10301 | RRM2 | P31350 | RSAD2 | Q8WXG1 |

| Input | UniProt Id | Input | UniProt Id | Input | UniProt Id |
|---|---|---|---|---|---|
| RSF1 | Q96T23 | RTN3 | O95197 | RTN4 | Q9NQC3 |
| RTN4R | Q9BZR6 | S100B | P04271 | S1PR5 | Q9H228 |
| SAP30 | O75446, Q9HAJ7 | SAT1 | Q9H2H9 | SCARB2 | Q14108 |
| SCO2 | P78563 | SDC3 | O75056 | SDCBP | O00560 |
| SDK2 | Q58EX2 | SEC24D | O94855 | SEC61G | P60059 |
| SELENOI | Q9C0D9 | SELL | P14151 | SEPTIN4 | O43236-6 |
| SERPINE2 | P07093 | SERPING1 | P05155 | SERPINH1 | P50454 |
| SESN1 | Q9Y6P5 | SGF29 | Q96ES7 | SH2D1B | O14796 |
| SHH | Q15465 | SHISA5 | Q8N114 | SIGIRR | Q6IA17 |
| SIGLEC1 | Q9BZZ2 | SIGLEC14 | Q08ET2 | SIGLEC5 | O15389, O43699 |
| SIGLEC7 | Q9Y286 | SIGLEC8 | Q96PQ1, Q9NYZ4 | SIK1 | O00567 |
| SIRPA | P78324 | SIRPG | Q9P1W8 | SIT1 | Q9NP91 |
| SKAP2 | O75563 | SLC15A2 | Q16348 | SLC15A4 | Q8N697 |
| SLC16A10 | Q8TF71 | SLC1A3 | P43003 | SLC22A23 | Q3SY84 |
| SLC24A4 | Q8NFF2 | SLC25A1 | P53007 | SLC25A6 | P12236 |
| SLC27A1 | Q6PCB7 | SLC27A3 | O14975 | SLC30A1 | Q9Y6M5 |
| SLC35B3 | Q9H1N7 | SLC39A3 | Q9BRY0 | SLC4A1 | P02730 |
| SLC4A10 | Q6U841 | SLC6A12 | P48065 | SLC7A7 | Q9UM01 |
| SLC7A8 | Q9UHI5 | SMG7 | Q92540 | SMG9 | Q9H0W8 |
| SMPD1 | P17405 | SMPD3 | Q9NY59 | SNRPN | P63162 |
| SNU13 | P55769 | SOCS1 | O15524 | SOCS2 | O14508 |
| SOCS3 | O14543 | SORBS3 | O60504 | SORT1 | Q99523 |
| SOS2 | P42766 | SP100 | P23497 | SP110 | P23497 |
| SPON2 | Q9BUD6 | SPRED1 | Q7Z699 | SPSB2 | Q99619 |
| SPTLC2 | O15270 | SQLE | Q14534 | SQOR | Q9Y6N5 |
| SQSTM1 | Q13501 | SRGAP2 | O43295, O75044 | ST3GAL3 | Q11203 |
| ST3GAL4 | Q11206 | ST7 | Q9Y561 | STARD10 | Q9Y365 |
| STARD8 | Q92502 | STAT1 | P42224-1 | STAT2 | P52630 |
| SYK | P43405 | TAB3 | Q8N5C8 | TAGLN2 | P37802 |
| TAP1 | Q03518 | TAP2 | Q03519 | TARBP2 | Q15633 |
| TAX1BP3 | O14907 | TBL1X | O60907 | TBL3 | Q12788 |
| TCF7 | P36402 | TCF7L2 | Q9NQB0 | TCN1 | P20061 |
| TCN2 | P20062 | TCTN3 | Q6NUS6 | TEAD4 | Q15561, Q15562 |
| TECR | Q5VVJ2 | THBD | P07204 | THRA | P10827-2 |
| TIA1 | P31483 | TIFA | Q96CG3 | TIMELESS | Q9UNS1 |
| TIMM10 | P62072 | TIMP2 | P16035 | TKTL1 | P29401 |
| TLE2 | Q04725 | TLR1 | Q15399 | TLR2 | O60603 |
| TLR4 | O00206 | TLR5 | O60602 | TLR7 | Q9NYK1 |
| TLR8 | Q9NR97 | TLR9 | Q9NR96 | TMED7 | Q9Y3B3 |
| TMOD2 | Q9NZR1 | TMX3 | Q96JJ7 | TNFAIP6 | P98066 |
| TNFRSF10B | O14763, Q9UBN6 | TNFRSF17 | Q02223 | TNFRSF21 | O75509 |
| TNFRSF25 | Q93038 | TNFSF10 | P50591 | TNFSF13 | O75888 |
| TNFSF13B | Q9Y275 | TNFSF14 | O43557 | TNNC2 | P02585 |
| TNRC6A | Q8NDV7 | TOM1L2 | O60784 | TOMM6 | Q96B49 |
| TOMM7 | Q9P0U1 | TOP2A | P11388 | TOR1B | O14657 |
| TOR4A | Q9NXH8 | TPM1 | P06753 | TPM2 | P07951 |
| TPX2 | Q9ULW0 | TRAF3 | Q13114 | TREML1 | Q86YW5 |
| TREX1 | Q9NSU2 | TRIM14 | Q14142 | TRIM21 | P19474 |
| TRIM22 | Q8IYM9 | TRIM25 | Q14258 | TRIM27 | P14373 |

| Input | UniProt Id | Input | UniProt Id | Input | UniProt Id |
|---|---|---|---|---|---|
| TRIM34 | Q9BYJ4 | TRIM38 | O00635 | TRIM5 | Q9C035 |
| TRIM56 | Q9BRZ2 | TRIM6 | Q9C030 | TRIM69 | Q86WT6 |
| TRIP12 | Q14669 | TRMT61B | Q9BVS5 | TSC22D3 | Q99576 |
| TSPAN33 | Q86UF1 | TSPO | P30536 | TSPOAP1 | O95153 |
| TST | Q16762 | TTC26 | A0AVF1 | TTLL4 | Q14679 |
| TTYH3 | Q9C0H2 | TUBA4A | P68366 | TXK | P42681 |
| TXNDC5 | Q8NBS9 | TYMP | P19971 | TYMS | P04818 |
| U2AF1L4 | Q8WU68 | UBA6 | A0AVT1 | UBA7 | P41226 |
| UBE2C | O00762 | UBE2G1 | P62253 | UBE2J2 | Q8N2K1 |
| UBE2L6 | O14933 | UBOX5 | O94941 | UBXN1 | Q04323 |
| UFD1 | Q92890 | UFL1 | O94874 | UHMK1 | Q8TAS1 |
| UHRF2 | Q96PU4 | UNC93B1 | Q9H1C4 | UPF3B | Q9BZI7 |
| URM1 | Q9BTM9 | USP1 | O94782 | USP15 | Q9Y4E8 |
| USP18 | Q9UMW8 | USP25 | Q9UHP3 | USP41 | Q3LFD5 |
| USP8 | P40818 | UXT | Q9UBK9 | VCAN | P13611 |
| VCPIP1 | Q96JH7 | VEGFB | P49765 | VIM | P08670 |
| VPS51 | Q9UID3 | VRK2 | Q86Y07 | VRK3 | Q8IV63 |
| WARS1 | P23381 | WIPF1 | O43516 | WLS | Q5T9L3 |
| WNT16 | Q9UBV4 | WNT2B | Q93097 | WSB1 | Q9Y6I7 |
| XAF1 | Q6GPH4 | XCL2 | Q9UBD3 | XPO1 | O14980 |
| XRN1 | Q8IZH2 | ZBP1 | Q9H171 | ZC3HAV1 | Q7Z2W4 |
| ZDHHC9 | Q9Y397 | ZEB2 | O60315 | ZFYVE9 | O95405-1 |
| ZNF2 | Q9BSG1 | ZNF333 | Q96JL9 | ZNF446 | Q9NWS9 |
| ZNF451 | Q9H7R0 | ZNF467 | Q7Z7K2 | ZNF484 | Q5JVG2, Q8N8J6 |
| ZNF496 | Q96IT1 | ZNF586 | Q9NXT0 | ZNF595 | Q8IYB9 |
| ZNF600 | Q6ZNG1 | ZNF668 | Q96K58 | ZNF684 | Q5T5D7 |
| ZNF74 | Q16587 | | | | |

| Input | Ensembl Id | Input | Ensembl Id | Input | Ensembl Id |
|---|---|---|---|---|---|
| ABCA1 | ENSG00000165029, ENST00000374736.7 | ABCG1 | ENSG00000160179 | ACSL1 | ENSG00000151726 |
| ACTA2 | ENSG00000107796 | ADAR | ENSG00000160710 | ALOX15 | ENSG00000161905 |
| APOD | ENSG00000189058 | ARF1 | ENSG00000143761 | ATF3 | ENSG00000162772 |
| AXIN2 | ENSG00000168646 | B2M | ENSG00000166710 | BCL2A1 | ENSG00000140379 |
| BCL2L14 | ENSG00000121380 | BRCA1 | ENSG00000012048 | BST2 | ENSG00000130303 |
| CASP1 | ENSG00000137752 | CAT | ENSG00000121691 | CAV1 | ENSG00000105974 |
| CCL2 | ENSG00000108691 | CCL20 | ENSG00000115009 | CCNA1 | ENSG00000133101 |
| CCNA2 | ENSG00000145386 | CCR1 | ENSG00000163823 | CD163 | ENSG00000177575 |
| CD274 | ENSG00000120217 | CDC45 | ENSG00000093009 | CDKN1A | ENSG00000124762 |
| CETP | ENSG00000087237 | CISH | ENSG00000114737 | CLDN5 | ENSG00000184113 |
| CLOCK | ENSG00000134852 | CXCL10 | ENSG00000169245 | CXCR4 | ENSG00000121966 |
| DBP | ENSG00000105516 | DHCR7 | ENSG00000172893 | EEPD1 | ENSG00000122547 |
| EGR1 | ENSG00000120738 | EIF2AK2 | ENSG00000055332 | EPHA4 | ENSG00000116106 |
| EXTL3 | ENSG00000012232 | EZH2 | ENSG00000106462 | FANCD2 | ENSG00000144554, ENST00000419585 |
| FCGR1A | ENSG00000150337 | FCGR1B | ENSG00000198019 | FGF9 | ENSG00000102678 |
| FLT4 | ENSG00000037280 | FTH1 | ENST00000273550 | GAMT | ENSG00000130005 |
| GBP1 | ENSG00000117228 | GBP2 | ENSG00000162645 | GBP3 | ENSG00000117226 |
| GBP4 | ENSG00000162654 | GBP5 | ENSG00000154451 | GBP6 | ENSG00000183347 |
| GBX2 | ENSG00000168505 | GCLC | ENSG00000001084 | GFPT1 | ENSG00000198380 |
| GHRL | ENSG00000157017 | HBG1 | ENSG00000213934 | HBG2 | ENSG00000196565 |

| Input | Ensembl Id | Input | Ensembl Id | Input | Ensembl Id |
|---|---|---|---|---|---|
| HERPUD1 | ENSG00000051108 | HLA-A | ENSG00000206503 | HLA-B | ENSG00000234745 |
| HLA-C | ENSG00000204525 | HLA-DQA1 | ENSG00000196735 | HLA-DQA2 | ENSG00000237541 |
| HLA-DRA | ENSG00000204287 | HLA-DRB4 | ENSG00000227357 | HLA-G | ENSG00000204632 |
| HLA-H | ENSG00000206341 | HNRNPA2B1 | ENSG00000122566 | HSPA1A | ENSG00000204389, ENSG00000215328, ENSG00000234475, ENSG00000235941, ENSG00000237724 |
| HSPA6 | ENSG00000173110 | ID2 | ENSG00000115738 | ID3 | ENSG00000117318 |
| IFI27 | ENSG00000165949 | IFI30 | ENSG00000216490 | IFI35 | ENSG00000068079 |
| IFI6 | ENSG00000126709 | IFIT1 | ENSG00000185745 | IFIT2 | ENSG00000119922 |
| IFIT3 | ENSG00000119917 | IFIT5 | ENSG00000152778 | IFITM1 | ENSG00000185885 |
| IFITM2 | ENSG00000185201 | IFITM3 | ENSG00000142089 | IFNG | ENSG00000111537 |
| IGHE | ENSG00000211891 | IGHG1 | ENSG00000211896 | IGHG4 | ENSG00000211892 |
| IKBIP | ENSG00000166130 | IL1RN | ENSG00000136689 | IL4R | ENSG00000077238 |
| IL6 | ENSG00000136244 | IL6R | ENSG00000160712 | IP6K2 | ENSG00000068745 |
| IRF1 | ENSG00000125347 | IRF5 | ENSG00000128604 | IRF7 | ENSG00000185507 |
| IRF9 | ENSG00000213928 | ISG15 | ENSG00000187608 | ISG20 | ENSG00000172183 |
| ITGA4 | ENSG00000115232 | ITGB1 | ENSG00000150093 | KDM1B | ENST00000297792 |
| KDM4B | ENST00000381759 | KIT | ENSG00000157404 | KLF4 | ENSG00000136826 |
| LIF | ENSG00000128342 | LMNB1 | ENSG00000113368 | MEF2C | ENSG00000081189 |
| MT2A | ENSG00000125148 | MUC1 | ENSG00000185499 | MX1 | ENSG00000157601 |
| MX2 | ENSG00000183486 | MYBL2 | ENSG00000101057 | NAMPT | ENSG00000105835 |
| NCAM1 | ENSG00000149294 | NPAS2 | ENSG00000170485 | NR1D1 | ENSG00000126368 |
| OAS1 | ENSG00000089127 | OAS2 | ENSG00000111335 | OAS3 | ENSG00000111331 |
| OASL | ENSG00000135114 | OR2AG1 | ENSG00000279486 | PML | ENSG00000140464 |
| PPIA | ENSG00000196262 | PRKCB | ENSG00000166501 | PSMB8 | ENSG00000204264 |
| PSME2 | ENSG00000100911 | PTCH1 | ENSG00000185920 | PTGDS | ENSG00000107317 |
| PTPN2 | ENSG00000175354 | RBBP8 | ENSG00000101773 | RGCC | ENSG00000102760 |
| RGL1 | ENSG00000143344 | RORA | ENSG00000069667 | RORC | ENSG00000143365 |
| RPLP0 | ENSG00000089157 | RPS2 | ENST00000343262 | RRM2 | ENSG00000171848 |
| RSAD2 | ENSG00000134321 | S100B | ENSG00000160307 | SCO2 | ENSG00000130489 |
| SERPINH1 | ENSG00000149257 | SESN1 | ENSG00000080546 | SHH | ENSG00000164690 |
| SLC27A1 | ENSG00000130304 | SOCS1 | ENSG00000185338 | SOCS2 | ENSG00000120833 |
| SOCS3 | ENSG00000184557, ENST00000330871 | SP100 | ENSG00000067066 | SQLE | ENSG00000104549 |
| SQSTM1 | ENSG00000161011 | STAT1 | ENSG00000115415 | TNFRSF10B | ENSG00000120889 |
| TNFRSF21 | ENSG00000146072 | TOP2A | ENSG00000131747 | TRIM14 | ENSG00000106785 |
| TRIM21 | ENSG00000132109 | TRIM22 | ENSG00000132274 | TRIM25 | ENSG00000121060 |
| TRIM34 | ENSG00000258659 | TRIM38 | ENSG00000112343 | TRIM5 | ENSG00000132256 |
| TRIM6 | ENSG00000121236 | TYMS | ENSG00000176890 | UXT | ENSG00000126756 |
| VIM | ENSG00000026025 | XAF1 | ENSG00000132530 | ZEB2 | ENSG00000169554 |

| Input | ChEBI Id |
|---|---|
| MAL | 15589 |

### 7. Identifiers not found

These 576 identifiers were not found neither mapped to any entity in Reactome.

| | | | | | | | |
|---|---|---|---|---|---|---|---|
| ABHD14A | ABTB3 | ADAP2 | ADGRE4P | ADGRL1 | AFF1 | AFG3L1P | AFTPH |
| AGAP1 | AHNAK | AHSA2P | AIDA | AIG1 | AIRE | ALKBH7 | AMIGO1 |
| AMN1 | ANKRD17 | ANKRD22 | ANKRD23 | ANXA2P1 | ANXA2R | ANXA3 | ANXA4 |
| AP5B1 | APOL2 | APOL4 | APOL6 | ARL5B | ARMCX6 | ARMT1 | ASCL2 |
| ASF1B | ASMTL | ASPM | ASPRV1 | ATP6V0E2-AS1 | ATRN | AZI2 | BAIAP3 |
| BATF2 | BAZ1A | BAZ2B | BCAR3 | BCL11B | BCL2L2 | BCL7B | BOK |
| BRSK1 | BSCL2 | BTBD3 | BTN2A3P | C10orf105 | C12orf4 | C12orf57 | C14orf132 |
| C15orf48 | C17orf58 | C1orf198 | C21orf91 | C3orf14 | C5orf24 | C8orf31 | C9orf72 |
| CA11 | CACHD1 | CALHM6 | CAMK2N1 | CAND2 | CAPRIN1 | CARD16 | CARD17 |
| CARD17P | CARD6 | CASP4LP | CCDC124 | CCDC146 | CCDC17 | CCDC90B | CCDC96 |
| CCL18 | CCM2 | CCNJL | CCNY | CD248 | CD6 | CD69 | CD7 |
| CD82 | CDCA7 | CDHR1 | CDK20 | CDR2L | CENPBD2P | CENPV | CEP85L |
| CFAP410 | CFAP44 | CFAP58 | CFAP58-DT | CHRM3-AS2 | CIDECP1 | CKAP2 | CKS2 |
| CLIC3 | CLIC4 | CLK4 | CLPP | CMAHP | CMPK2 | CMTR1 | CNIH4 |
| CNP | CNTNAP3 | COCH | COLQ | CPNE2 | CPQ | CRIP2 | CSRNP1 |
| CTDSPL | CTRL | CUTA | CUX2 | CXCR2P1 | CXorf21 | CXorf65 | Calcineurin |
| DALRD3 | DBNDD1 | DCANP1 | DCBLD1 | DDX60 | DDX60L | DEAF1 | DEXI |
| DHRS11 | DIPK2A | DISC1 | DLEU2L | DMTF1 | DMXL2 | DNAH6 | DNAJB12 |
| DNAJB14 | DNAJB5 | DONSON | DPCD | DUSP22 | EEF1A1P6 | EEF1GP1 | EEIG2 |
| EFCAB12 | ELAPOR1 | EMC9 | ENKD1 | EPB41L4A-AS1 | EPB42 | ERGIC3 | ETV5 |
| ETV7 | EVA1B | EVA1C | EWSR1 | EXOC3L1 | EXOG | F8A1 | FAM102A |
| FAM153B | FAM177A1 | FAM177B | FAM209B | FAM225A | FAM53C | FAM72B | FAM76B |
| FAM8A1 | FASTKD1 | FAT4 | FAXDC2 | FBXO39 | FCGBP | FCGR1CP | FCGR2C |
| FCRL5 | FEZ1 | FLVCR2 | FNDC3A | FRMD3 | FTH1P3 | GALM | GASK1B |
| GATAD1 | GBP1P1 | GCNT2 | GFOD2 | GLRX2 | GOLGA2P10 | GOLGA6L1 | GOLGA8B |
| GORAB | GPA33 | GPATCH2 | GPBP1 | GPR162 | GPRASP1 | GRAMD1B | GRIPAP1 |
| GTF2IP1 | GUCD1 | HDGFL2 | HES4 | HESX1 | HLA-DRB6 | HLA-J | HLA-K |
| HMGB1P3 | HOPX | HOXB7 | HSPBP1 | IFI44 | IFI44L | IGHGP | IGHV1-18 |
| IGHV3-15 | IGHV3-21 | IGHV4-61 | IGHV5-51 | IGKV1-27 | IGKV3D-15 | IGLL5 | IKZF4 |
| INKA2 | IRF1-AS1 | IRX3 | ITIH5 | ITPRIPL2 | Immunoglobulin | JPX | KATNAL1 |
| KBTBD4 | KIAA0319L | KIAA0895L | KIAA1109 | KIAA1958 | KLHDC7B | KLHDC8B | KLRC3 |
| KRT10-AS1 | KRTCAP2 | L-tryptophan | LACTB | LAMP5 | LBH | LEF1-AS1 | LENG8 |
| LGALS9B | LINC00487 | LINC00857 | LINC00937 | LINC00999 | LINC01002 | LINC01061 | LMO4 |
| LRMDA | LRRC15 | LRRC23 | LRRC26 | LRRC37A4P | LRRC56 | LRRN3 | LTK |
| LY9 | LYRM9 | LYSMD2 | MACROD2 | MAEA | MANSC1 | MAP3K7CL | MAP4K1 |
| MAP7 | MARCHF1 | MARCHF5 | MAST2 | MCF2L-AS1 | MCRIP2 | MELK | MICAL2 |
| MID2 | MKI67 | MMP28 | MOB3A | MOB3B | MOK | MORC2-AS1 | MOXD1 |
| MPZL1 | MPZL2 | MS4A1 | MS4A4A | MS4A7 | MTPN | MTRF1 | MTSS1 |
| MTURN | MTUS1 | MUSTN1 | MXI1 | MYO7B | MYOF | MZB1 | N4BP2L2 |
| NAIP | NAP1L3 | NBDY | NCOA7 | NDRG2 | NELL1 | NEXN | NFE2L3 |
| NIPSNAP3A | NLRP2 | NLRP7 | NOP53 | NPTXR | NSG1 | NUDT16L2P | NUFIP2 |
| NUSAP1 | ODAD4 | ODF3B | ONECUT2 | OR52K3P | OST4 | OTUD6B | PABPC4 |
| PARP12 | PARP15 | PARP3 | PATL2 | PAWR | PCDHGB6 | PDCD2L | PDIA3P1 |
| PDXP | PDZD4 | PDZK1IP1 | PGAM4 | PGAP3 | PGAP6 | PHACTR1 | PHF11 |
| PHLDB1 | PHLDB2 | PIK3IP1 | PITPNC1 | PLEKHB1 | PLLP | PLSCR1 | PLXDC1 |
| POGLUT3 | POLR2J2 | PPHLN1 | PPM1H | PPP1R32 | PPP3R2 | PRAF2 | PRF1 |

| | | | | | | | |
|---|---|---|---|---|---|---|---|
| PRKCQ-AS1 | PRORSD1P | PRRG4 | PRRT2 | PRSS30P | PRSS33 | PRXL2A | PRXL2C |
| PSME2P2 | PSTPIP2 | PTBP2 | PTGES3P1 | PTMA | PTP4A1 | R3HCC1 | RABGAP1L |
| RAD54B | RASD1 | RASGEF1B | RASSF3 | RBM15B | RBM45 | RBPMS2 | RECQL |
| REEP6 | REM2 | RETREG1 | RFLNB | RFX2 | RGPD1 | RIMS3 | RN7SL1 |
| RNF126P1 | RNF148 | RNF5P1 | ROPN1B | RPH3A | RPL10P3 | RPL13AP5 | RPL3P4 |
| RPL7P1 | RPL7P9 | RPP25L | RPS4XP22 | RPSAP9 | RSPH9 | RTN2 | RTP4 |
| RUBCN | RUFY3 | RUFY4 | RWDD4 | SALL2 | SAMD14 | SAMD4A | SAMD9 |
| SAMD9L | SASH1 | SASS6 | SCAMP5 | SCARNA9 | SCFD2 | SCML1 | SCYL1 |
| SDR39U1 | SEC14L1 | SECTM1 | SELENBP1 | SEMA4C | SERPINF1 | SESTD1 | SGTB |
| SH3TC1 | SH3YL1 | SHFL | SIGLEC17P | SIPA1L2 | SLC25A39 | SLC26A8 | SLC41A3 |
| SLF1 | SLFN11 | SLFN12 | SMCHD1 | SMCO4 | SNHG29 | SNHG32 | SNHG5 |
| SNORA32 | SNORD32A | SNTA1 | SNX10 | SP140 | SPATS2L | SPG11 | SPG21 |
| SPIN4 | SPINK2 | SPNS3 | SPOCK2 | SPRYD3 | SRBD1 | SRFBP1 | SRGAP2B |
| SRGAP2C | SRRD | SS18 | SSH3 | STAC3 | STIL | STK16 | STMN3 |
| STX11 | SULF2 | SUSD1 | SYTL2 | TAGLN | TARDBP | TARP | TASL |
| TBC1D8 | TCEA3 | TCP11L1 | TDRD7 | TENT5A | TFEC | TGFB1I1 | TGM2 |
| TGM3 | THNSL1 | TIGIT | TIPRL | TLCD3A | TMEM123 | TMEM127 | TMEM140 |
| TMEM144 | TMEM176A | TMEM176B | TMEM178B | TMEM191A | TMEM191C | TMEM203 | TMEM204 |
| TMEM207 | TMEM258 | TMEM259 | TMEM268 | TMEM62 | TMEM87B | TMEM8B | TMEM97 |
| TMIGD2 | TMPRSS15 | TMTC1 | TMX4 | TOP1MT | TOR2A | TP53INP2 | TPRA1 |
| TPT1 | TRABD2A | TRAFD1 | TRANK1 | TRAV1-2 | TRDC | TRDV2 | TRGC1 |
| TRGV9 | TRIM6-TRIM34 | TSPAN18 | TTC21A | TTC39C | TXNDC12 | UBE2Q2P6 | UBL7 |
| UBQLNL | VAMP5 | VNN3 | VOPP1 | VPS50 | VWCE | WDFY1 | WDR5B |
| WDR73 | XKR8 | XKRX | XPO7 | YBX3P1 | YIF1B | YTHDC2 | ZCCHC2 |
| ZFC3H1 | ZFP57 | ZNF117 | ZNF174 | ZNF185 | ZNF358 | ZNF438 | ZNF575 |
| ZNF674 | ZNF814 | ZNFX1 | ZNRD2 | ZSCAN16 | ZSCAN18 | ZUP1 | kynurenine |
